# Quadratic trends: a morphometric tool both old and new

**DOI:** 10.1101/2023.03.23.533997

**Authors:** Fred L. Bookstein

## Abstract

The original exposition of the method of “Cartesian transformations” in D’Arcy Thompson’s great essay *On Growth and Form* of 1917 is still its most cited. But generations of theoretical biologists have struggled ever since to invent a biometric method aligning that approach with the comparative anatomist’s ultimate goal of inferring biologically meaningful hypotheses from empirical geometric patterns. Thirty years ago our community converged on a common data resource, samples of landmark configurations, and a currently popular biometric toolkit for this purpose, the “morphometric synthesis,” that combines Procrustes shape coordinates with thin-plate spline renderings of their various multivariate statistical comparisons. But because both tools algebraically disarticulate the landmarks in the course of a linear multivariate analysis, they have no access to the actual anatomical information conveyed by the arrangements and adjacencies of these locations as they combine in pairs or higher numbers into substructures. This paper explores a geometric approach circumventing these fundamental difficulties: an explicit statistical methodology for the simplest nonlinear patterning of these comparisons at their largest scale, their fits by what Sneath (1967) called quadratic trend surfaces. After an initial quadratic regression of target configurations on a template, the proposed method ignores individual shape coordinates completely, replacing them by a close reading of the regression coefficients accompanied by several new diagrams, notably the exhaustive summary of each regression by an unfamiliar biometric ellipse, its circuit of second-order directional derivatives. These novel trend coordinates, directly visualizable in their own coordinate plane, do not reduce to any of the usual Procrustes or thin-plate summaries. The geometry and algebra of these second-derivative ellipses seem a serviceable first approximation for applications in evo-devo studies and elsewhere. Two examples are offered, one the classic growth data set of Vilmann neurocranial octagons and the other the Marcus group’s data set of midsagittal cranial landmarks over most of the orders of the mammals. Each analysis yields startling new findings inaccessible to the current GMM toolkit. A closing discussion suggests a variety of ways by which innovations in this spirit might burst the current strait-jacket of Procrustes coordinates and thin-plate splines that together so severely constrain the conversion of landmark locations into understanding across our science.

## I. Introduction

The main thrust of this paper can be distilled down to the pair of diagrams in Figure 1. They present a novel ordination of the *quadratic trend descriptor* intended to help rebuild our current toolkit for the landmark data sets of geometric morphometrics (GMM). The paper’s main empirical contribution, Section IV, is a new pattern analysis filling a major gap in the current GMM tookit. This first figure conveys the basic idea of the new ordination method: conversion of two-dimensional landmark configuration data into an explicit representation of just its quadratic trends.

**Figure 1.**
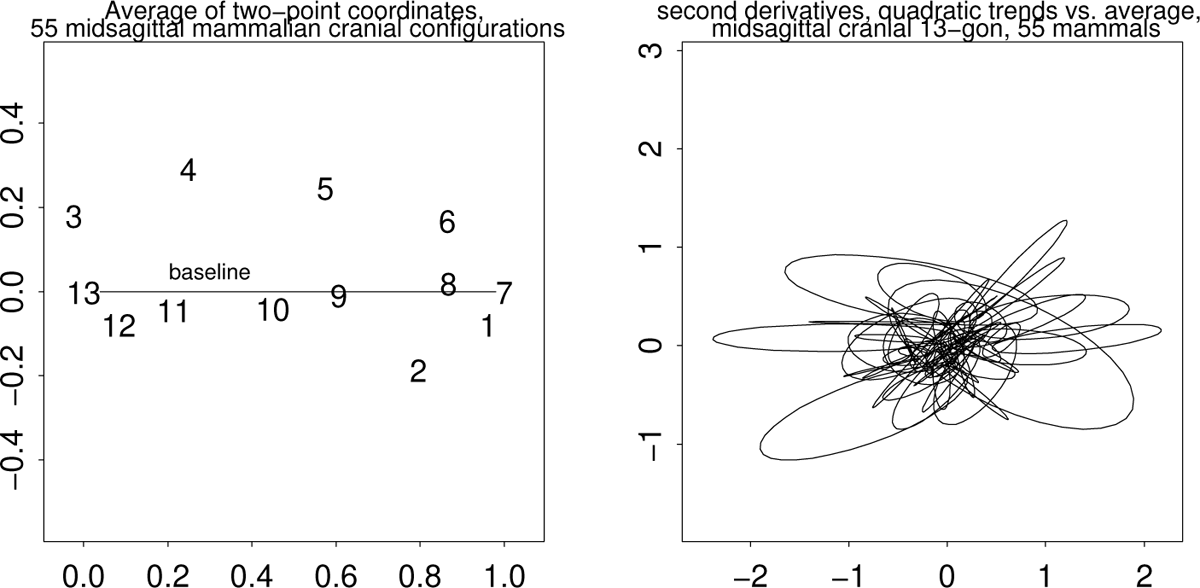
Principal methodological theme of this paper: a novel ordination of the *quadratic trend descriptor* apposite to the landmark data sets of geometric morphometrics, here as fitted in Section IV to a 13-point midsagittal cranial landmark configuration for each of 55 mammal specimens from 23 orders. Each ellipse traces the second derivative of the quadratic trend fit around the circle of directions from the sample average with respect to a convenient posteroanterior baseline (posterior foramen magnum to premaxilla). (left) The average of the 55 configurations of 13 midsagittal cranial landmarks in this example. The thirteen midsagittal cranial landmarks are as follows: 1, anterior symphysis of mandible; 2, posterior symphysis of mandible; 3, inion sagittal; 4, frontal-parietal sagittal; 5, frontal-nasal sagittal; 6, tip of nasal sagittal; 7, tip of premax sagittal; 8, premax-maxillary sagittal; 9, maxillary-palatine sagittal; 10, posterior palate; 11, basisphenoid-basioccipital; 12, anterior foramen magnum; 13, posterior foramen magnum. (right) All 55 ellipses of directional second derivatives for the 55 quadratic trend fits from the configuration at left.

The data set on which the figure is based comprises the 13-point midsagittal subset of a 35-landmark cranial configuration originally exploited in Marcus et al. (2000). While that representation relied on Procrustes shape coordinates, the registration here, in keeping with the recommendation of Bookstein (2023), instead uses a two-point coordinate system (Bookstein, 1986, 1991) with baseline from posterior foramen magnum to tip of premaxilla in this sagittal plane. Each comparison is from the configuration in the left-hand panel here, the average of these 55 representatives of 23 mammalian orders, to one of the individual configurations.

When the data of each 13-gon are fitted as a quadratic trend over that two-point average, there result the 55 four-panel figures collected in the Supplement: for each specimen, a Cartesian grid of the fitted trend, a polar grid of the same, a tracing of a half-unit circle deformed quadratically, and, as collected in the right-hand panel of Figure 1, the quadratic part of each fit visualized as explained in Section II by the ellipse of its second derivatives in every direction with respect to that two-point baseline. Linear multivariate analysis of these fits can proceed either by analysis of their six coefficients or instead by an equivalent eight-coordinate data set, the “cardinal diameters” (easterly, northeasterly, northerly, northwesterly) of their ellipses, likewise to be introduced in Section II; but non-linear analyses offer considerable power as well. There result new pattern analyses of these trends per se, the curving of coordinate lines whose graphical power has so intrigued all of us since D’Arcy Thompson.

This theme of visible curving was replaced (unfortunately, in my opinion) over the development of today’s GMM by an inappropriate surrogate arising instead from multivariate analyses (particularly principal-component analyses) of the otherwise disarticulated shape coordinates of the Procrustes approach. An earlier graphical innovation, my thin-plate spline deformation grid, has proved insufficient to restore the missing articulation with the anatomical sciences. In their place I foreground these ellipses as a first step in the replacement of the current GMM toolkit by a successor capable of generating hypotheses more closely aligned with the language of organismal anatomy. A praxis for feature analysis of the gradients and other nonlinear aspects of form-comparisons, such as this article sketches, could be a crucial component of the return of GMM to any future state-of-the-art biometric toolkit for either evolutionary or developmental studies.

### Beginning with D’Arcy Thompson

The new biometric methodology this paper introduces realizes a very old programme: the production of hypotheses about the causes or consequences of organic form from geometric observations of those forms as represented in line drawings. Such a thrust is over a century old, deriving from the first edition (1917) of the celebrated essay *On Growth and Form* by the polymath D’Arcy W. Thompson. For three-quarters of a century after that initial provocation — right through 1993 — the literature of this purpose remained disorganized, a range of approvals and disapprovals every decade or so without much of a consensus. D’Arcy Thompson’s much-quoted original example still stands as the clarion announcement of his purpose, and as he was a master of English prose style, it is best to quote him in his own words. From the most readily available edition (1961, abridged by John Tyler Bonner), pages 275–6 and 300–301:

> The deformation of a complicated figure may be a phenomenon easy of comprehension, though the figure itself have to be left unanalyzed and undefined. This process of comparison, recognizing in one form a definite permutation or deformation of another, apart altogether from a precise and adequate understanding of the original ‘type’ or standard of comparison, lies within the immediate province of mathematics. … When the morphologist compares one animal with another, point by point or character by character, these are too often the mere outcome of artificial dissection and analysis. Rather is the living body one integral and indivisible whole, in which we cannot find, when we come to look for it, any strict dividing line even between the head and the body, the muscle and the tendon, the sinew and the bone. Characters which we have differentiated insist on integrating themselves again, and aspects of the organism are seen to be conjoined which only our mental analysis had put asunder. The co-ordinate diagram throws into relief the integral solidarity of the organism, and enables us to see how simple a certain kind of correlation is which had been apt to seem a subtle and a complex thing.
>
> But if, on the other hand, diverse and dissimilar fishes can be referred as a whole to identical functions of very different coordinate systems, this fact will of itself constitute a proof that variation has proceeded on definite and orderly lines, that a comprehensive ‘law of growth’ has pervaded the whole structure in its integrity, and that some more or less simple and recognizable system of forces has been in control. It will not only show how real and deep-seated is the phenomenon of ‘correlation’ in regard to form, but it will also demonstrate the fact that a correlation which had seemed too complex for analysis or comprehension is, in many cases, capable of very simple graphical expression.

And then his most celebrated example, still on t-shirts to this day:

> [One DWT figure] is a common, typical *Diodon* or porcupine-fish, and [in another DWT figure] I have deformed its vertical co-ordinates into a system of concentric circles, and its horizontal co-ordinates into a system of curves which, approximately and provisionally, are made to resemble a system of hyperbolas. The old outline, transferred in its integrity to the new network, appears as a manifest representation of the closely allied, but very different looking, sunfish, *Orthagoriscus mola.* This is a particularly instructive case of deformation or transformation.

No, it is not “particularly instructive” in any contemporary sense — for one thing, neither circles nor hyperbolas are among the curves that characterize either the thinplate splines of the current consensus or the quadratic trends explored in this paper. But Thompson’s insistence on a “simple graphical expression” has fired the imagination of many of us over the century since his argument first appeared, while a like number of counterarguments have appeared to caution the enthusiasm of the same readers. An early counterargument was Karl Przibram’s (1923:14), insisting that no comparisons of this sort could be regarded as biologically credible unless and until they could be reproduced repeatedly in an experimental setting (such as the laboratories of his Vienna ‘Vivarium,’ Müller, ed., 2017). Julian Huxley (1932) promulgated the differential model log(*y*) = *a* log(*x*) for distances *x* and *y* only to realize that if such a model applied to, e.g., the sides of a rectangle, with different *a*’s, it didn’t apply to the diagonal. Peter Medawar (1945) circumvented the difficulty by diagramming only the sides of rectangles, not their diagonals, in his contribution to a Thompson Festschrift using human body growth as an example (but see also Richards and Kavanagh 1945, the contribution following on his in the same volume). Robert Sokal and Peter Sneath (1963) present several earnest attempts in the spirit of Thompson but end up concluding (pages 82–83), “No general and simple methods seem yet to have been developed for extracting the factors responsible for such transformations. … It is not easy to see how many separate factors are needed to express more complicated examples” where “not only are the grid lines deformed in several ways, but the deformation is different in different parts of the skull. What would be useful would be a way of extracting the minimum number of factors that would account for the difference in form. … It would probably be at first necessary to mark operationally homologous points [today’s ‘landmarks’] on the diagrams before feeding them into the computer.”

Four years later, this same Peter Sneath published the first attempt to properly quantify the issues here. He adopted a technique then under active exploitation in geology, the method of trend surfaces (now a component of the subdiscipline known as geostatistics) to apply to two variables over the same map, which could be taken as the coordinates of the corresponding points of the image of another organism entirely. Sneath (1967) offers a “partial solution” to the problems set down in his earlier collaboration with Sokal. He claims success in a first goal, “to estimate numerically the overall similarity between two figures, i.e., the gross difference in shape,” and proceeds to demonstrate using “sagittal sections of four hominoid skulls.” The “overall similarity” he suggests is close to Procrustes distance as the current consensus has it, and the “trend surfaces” he computes for his quadratic case are the same as those of this paper. Unfortunately, he pays no attention to the coefficients of those formulas, noting only the displacements at each landmark in turn. Each of the six comparisons among his four specimens is diagrammed and verbalized separately. There is no summary ordination, merely a hope that the sum of squared differences in the coefficients might serve as some sort of inverse index of “phenetic affinity.”

This was roughly the state of the art ten years later when I published my doctoral dissertation (Bookstein, 1978) that, picking up on an alternate theme of the 1940’s, attempted to customize a coordinate system for the description of the change per se, not the forms being compared. The biorthogonal grid method, however, had the same flaw as Sneath’s: it applied to only a single transformation — a single pair of forms — at a time.

### The “morphometric synthesis” of 1993

Shortly afterwards, however, a series of innovations resulted in the “morphometric synthesis” that this paper is now trying to replace. The first of these contributions was my announcement of two-point shape coordinates (Bookstein, 1986), a statistical space that supports applications of conventional tactics like mean difference and regression to the shape of configurations of points by algebraic manipulations of their Cartesian coordinates in a rigorous, theorem-governed way. Shortly afterwards came a 1988 conference (Rohlf and Bookstein, eds., 1990), the journal announcement (Bookstein 1989), and finally the book form (Bookstein 1991) of the thin-plate spline model for deformation, which explicitly converts landmark-driven shape comparisons to grids over the picture of the organism. Meanwhile Kanti Mardia and John Kent (University of Leeds) had tightly tied a crucial additional multivariate tool, principal components analysis, to coordinate data via the Procrustes shape space David Kendall (1984) had announced via the mathematical literature. The fusion of these latter two tools in Jim Rohlf’s NTSYS software package was the core of the NATO Advanced Studies Institute of 1993 organized by Leslie Marcus at Il Ciocco, Italy (Marcus et al, eds., 1996). It was there that Rohlf and I announced the “morphometric synthesis,” which by then had included several other extensions (to semilandmarks, to symmetry). At about the same time, the corresponding rigorous probability theory that had been developed in parallel by Mardia, Kent, and their students Ian Dryden and Colin Goodall was beginning to appear in the formal statistical literature (cf. Kent and Mardia 1994), a theory soon codified in the graduate textbook *Statistical Shape Analysis* (Dryden and Mardia, 1998, second edition, 2016). The synthesis emphasizes Procrustes shape coordinates over the two-point version because of their much greater suitability for principal components analysis with respect to Procrustes distance.

This serendipitous confluence became the core computational engine for data graphics and ordination of samples across a wide range of biological investigations, particularly in anthropology and paleontology, fields not susceptible to Przibram’s challenge of laboratory confirmation. Over time it has become the theme of pedagogy directed at biologists over a wide range of levels of sophistication. Whether course or workshop, most of these curricula favor the same shared workflow — gather your landmark coordinates, convert to Procrustes shape coordinates by John Gower’s (1975) Generalized Procrustes Analysis, carry out any of the popular multivariate analyses there (principal components, canonical variates, multivariate analysis of variance or covariance, partial least squares) on samples (and, more recently, on their phylogenetic contrasts), diagram the analyses by thin-plate spline, and publish.

But over the course of this development Thompson’s original goal, the pursuit of simple explanations hinting at meaningful hypotheses, was subordinated to a reversed logic in which extant hypotheses were “tested” using morphometric arithmetic. Philosophers of science often refer to this pejoratively as the context of confirmation, not discovery. Today, thirty years on, the synthesis is overdue for major revision. The multivariate aspects of this context have been revolutionized in most other sciences by the advent of techniques of machine learning and artificial intelligence, while GMM’s tools have remained where they were created thirty or more years ago. To put matters bluntly, GMM techniques no longer produce surprising findings any more, findings that lead to unexpected hypotheses about the causes or consequences of organismal form. Thin-plate splines do not often meet Thompson’s criterion of being simpler than the data that drive them.

Thus it is time to revisit Thompson’s original goal, the production of interpretable diagrams simpler than the anatomies they compare. Bookstein (2023) has already noted how the Procrustes method, by prohibiting the investigator from rotating a Thompsonian coordinate grid, interferes with the generation of optimally simple accounts of findings. This paper is the second step in this programme: the construction of an explicit statistical method for the transformation grids that Sneath could already compute, by imitating what the geologists were doing, but did not know how to compare or extend to ordinations of samples. Once the new method is adjoined to an accessible statistical package, our community might discover its strengths and weaknesses quite a bit more rapidly than was the case for Procrustes tools and the thin-plate spline.

### Purpose and contents of this paper

Like the original paper on the thin-plate spline (Bookstein, 1989), this paper has two distinct purposes: to teach the mathematics driving the new praxis for ordinating quadratic growth-gradients and other quadratic trends, and also to present a pair of potentially classic examples, one involving growth and the other involving adaptive radiation, in order to hint at the kinds of biometric questions that might now enjoy the possibility of explicitly geometrical answers along with the diagrams that help convert those answers into biological hypotheses. The outline of the rest of this paper is as follows. Section II, which is mostly elementary college geometry, retrieves a simple fact about parabolas that can be made to apply to the quadratic case of the regressions Sneath was already demonstrating half a century ago. It may surprise the reader that the same elliptical shape we’ve used since the 1880’s to characterize linear regression has an equally promising role to play in these simplest *non*linear regressions, the quadratic trend fits. (I also demonstrate that Thompson made a completely avoidable mistake back in 1917 when he failed to consider polar coordinate grids as well as Cartesian grids for the diagramming of his comparisons.)

Section III shows how the method affords a reanalysis of one aspect of a classic data set (the growth of Vilmann’s neurocranial octagons) in such a way as to lead automatically to a report of its features, which turns out to match one of the special cases surveyed in Section II. Section IV is a more challenging reanalysis, one well beyond the usual bounds we set on diversity of data sets that will yield morphometric sense: the sample of over 50 mammal skulls originally assembled for just this sort of challenge by Marcus and his colleagues in 2000. The analysis by Marcus et al., in my view, was not fully a success — the finding was only that Procrustes analysis had something to say about mammal phylogeny, but not what that message actually was. The multivariate analysis in Section IV is novel in that once the quadratic trend is fitted to landmark configurations, the landmarks per se are completely ignored in favor of the features of those six-dimensional derived formalisms instead. A closing Discussion, Section V, is, I hope, the foreshadowing of a much more nuanced, more demanding use of GMM to generate actual biological understanding. It touches on the deeper question of exactly what quantities are appropriate for minimizing by some empirical biological model, but also on the importance of a pre-existing language of graphical reportage that must be present from the beginning whenever a quantitative method of describing biometric morphology is under development.

### II. Geometric fundamentals

#### A simple fact about parabolas

Much of the geometry needed for the new approach to quadratic trend analysis is elementary. A first underlying principle is simple indeed: the midpoints of chords of a fixed span over a parabola all lie at the same distance from the parabola underneath. Writing the parabola as *y* = *x*^2^, this is the identity (*x* + 1)^2^ + (*x* − 1)^2^ */*2 − *x*^2^ = 1, which is the same as the coefficient of *x*^2^ in the parabola’s formula. Figure 2 confirms the identity for several chords all of projected length 2 over the parabola *y* = *x*^2^*/*2. As you see from the equality of all the heavy vertical segments, the midpoint of each chord between *x*’s separated by 2 is at height ½ over the curve, the same as the coefficient of *x*^2^ in the formula — half of the second derivative in question, and constant everywhere along the parabola. The identity is more familiar to the mathematician after being multiplied by two: it is the equality of the second *derivative* of the parabola with its *second difference,* the formula 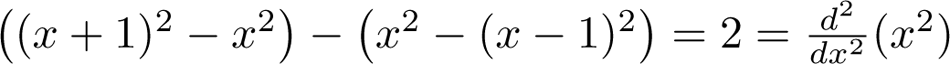.

**Figure 2.**
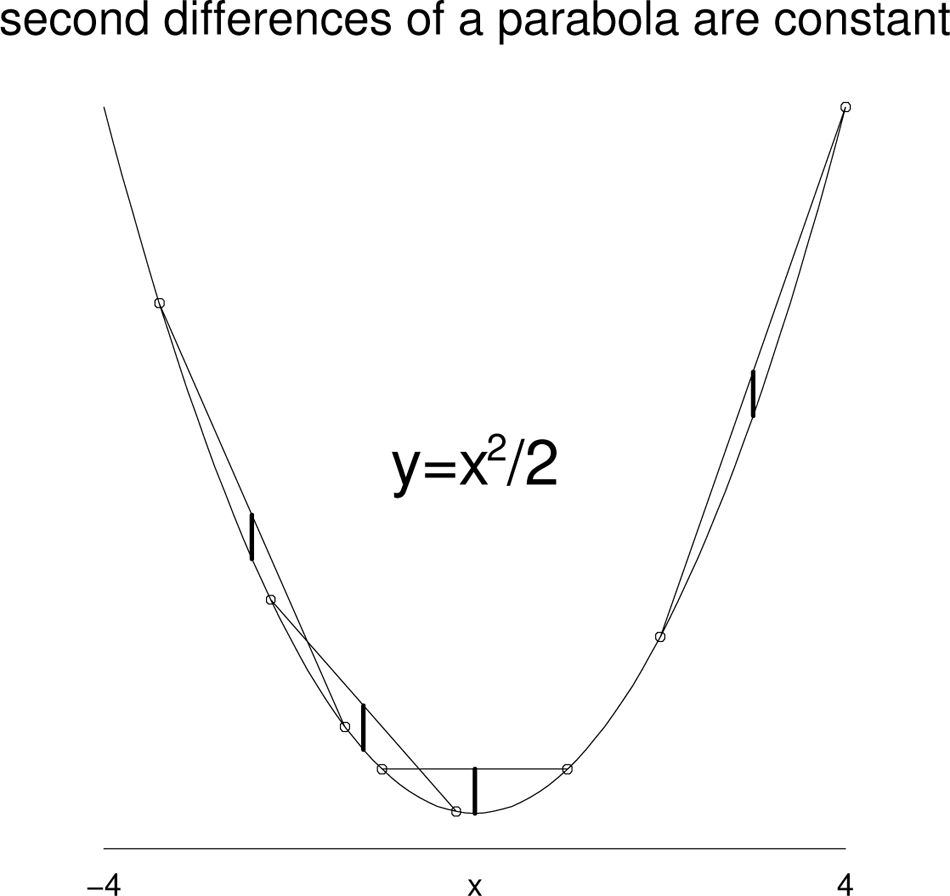
An ancient identity: the second difference of a parabola is a constant, equalling the second derivative of the curve’s formula. The proposition is easily confirmed once both sides are divided by two: invariance of the heavy segments, each the height over the parabola of the midpoint of any chord whose projection on the *x*-axis is a fixed interval (here, two units).

#### In two dimensions: ellipses and their cardinal diameters

The same proposition, appropriately reinterpreted, turns out to apply to quadratic trends (which, direction by direction, *are* parabolas) in two or any other higher count of dimensions. In every direction, the second *difference* is equal to the second *derivative* in that direction, which, for any quadratic trend (such as the quadratic regression of a target configuration on some template), is constant. But in higher dimensions another useful property emerges: the value of this second directional derivative lies on an exact ellipse (in two dimensions; in 3D it will be an ellipsoid). Figure 3 confirms this by a clever choice of coordinate systems borrowed from the morphometric methodology to be introduced presently. Begin with either lopsided oval curve, the one drawn in tiny plus signs or the one drawn in tiny times signs. Either serves as an arbitrary pedagogical choice of an example quadratic trend as rendered (and this step is crucial) by its effect on a unit circle as in the various examples of Sections III and IV. Here that curve is the example of no particular symmetries or other idiosyncrasies and with the identity as its linear term. Notice that the formula includes six coefficients, one each for *x*^2^*, xy,* and *y*^2^ for each of the two Cartesian coordinates of a grid point in two dimensions.

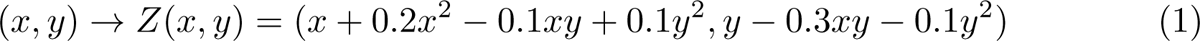

**Figure 3.**
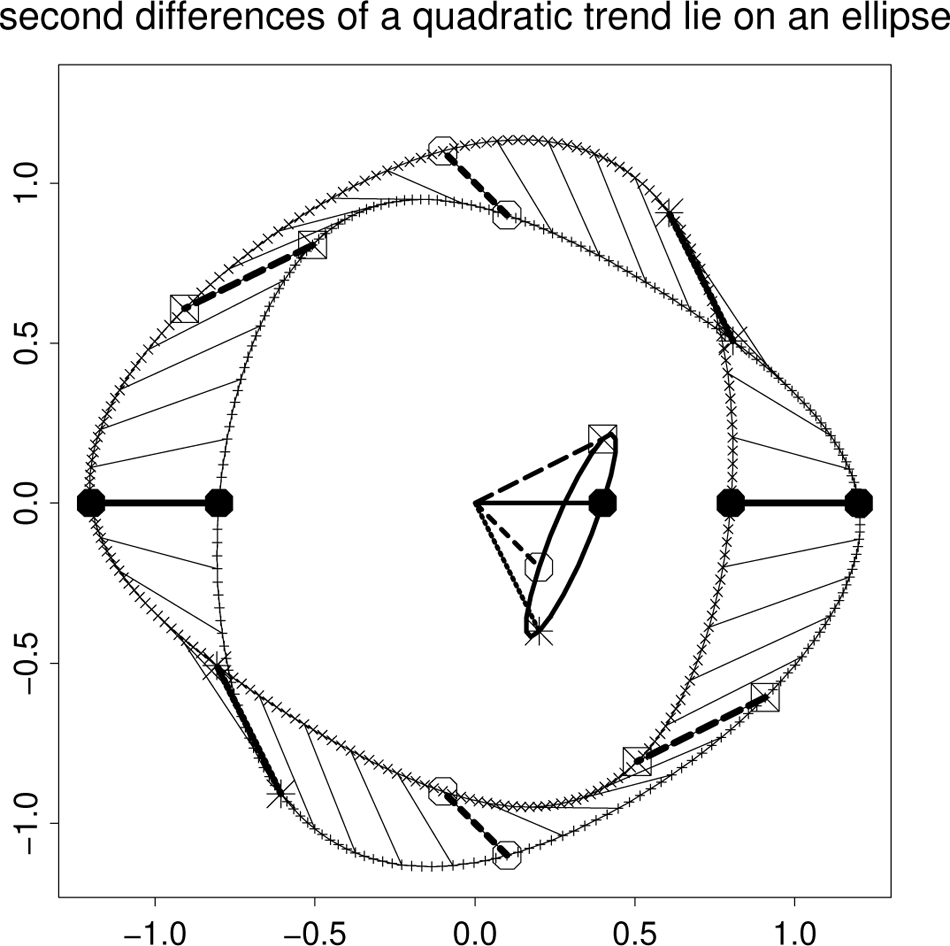
The same for a two-dimensional (quadratic) trend fit. Symbols for highlighted points along the curves will be introduced in the next figure. Curve of + signs: the effect of the quadratic trend on a unit circle around the origin of coordinates. Curve of × signs: the opposite curve, *x* and *y* replaced by −*x* and −*y*. See text.

The quadratic trend function leaves the origin (0, 0) unchanged, so, copying the formula for the parabola, the second difference across any diameter of the unit circle is just the sum of the values *Z*(*x, y*) and *Z*(−*x,* −*y*) in the direction (*x, y*) = (cos *θ,* sin *θ*) over the full circle of angles *θ* corresponding to points on the circle. But because all the quadratic terms in the formula are invariant when both *x* and *y* are multiplied by −1, the second difference *Z*(*x, y*) + *Z*(−*x,* −*y*) is the same as the simple difference *Z*(*x, y*) − *Z*^′^(−*x,* −*y*) where *Z*^′^ is the function −*Z,* reflection of the function *Z* in the origin.

Figure 3 draws the values of the function *Z* with tiny + signs and those of *Z*^′^ with tiny × signs, and their difference is drawn in light segments every 9^◦^ and heavy segments at the four cardinal directions (lines at 0^◦^, 45^◦^, 90^◦^, and 135^◦^ to the horizontal), identified by the same four symbols that will be assigned them in later figures of this paper. These heavy segments, translated to the origin (0, 0) of both coordinate systems, are four out of the continuum that traces the second-difference ellipse that losslessly, indeed redundantly, encodes the equation of the quadratic trend formula (1) driving it. While the curves of + signs and × signs are obviously not ellipses, the curve of their differences in this mixed registration must be, for the same reason that the second differences of an ordinary parabola (Figure 2) must be constant. Note that each point of the ellipse arises from two diametrically opposite segments of the original circuit.

Figure 4 shows how the analysis in Figure 3 can be attached to a comparison of landmark configurations in order to represent the quadratic trend of interest as an ellipse that can be submitted to explicit multivariate statistical analysis. For this example (and all the others of this section) the template is taken simply as a 3 × 3 grid of squares one unit on a side, a template that will be deformed by a quadratic formula written out in advance instead of being computed by a regression. The heading presents the coefficients of that quadratic trend in the same order as the example in Figure 3 or formula (1). Here that formula is as printed over the upper left panel. I cobbled this together to be mainly the *xy* term of the *x*-coordinate regression together with the *x*^2^ term of the *y*-coordinate regression, with a little effect of the other four terms. At above right is the better rendition of this same deformation, now as a polar coordinate grid (Bookstein 2022).

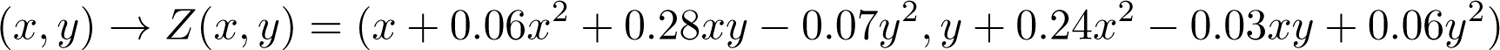

**Figure 4.**
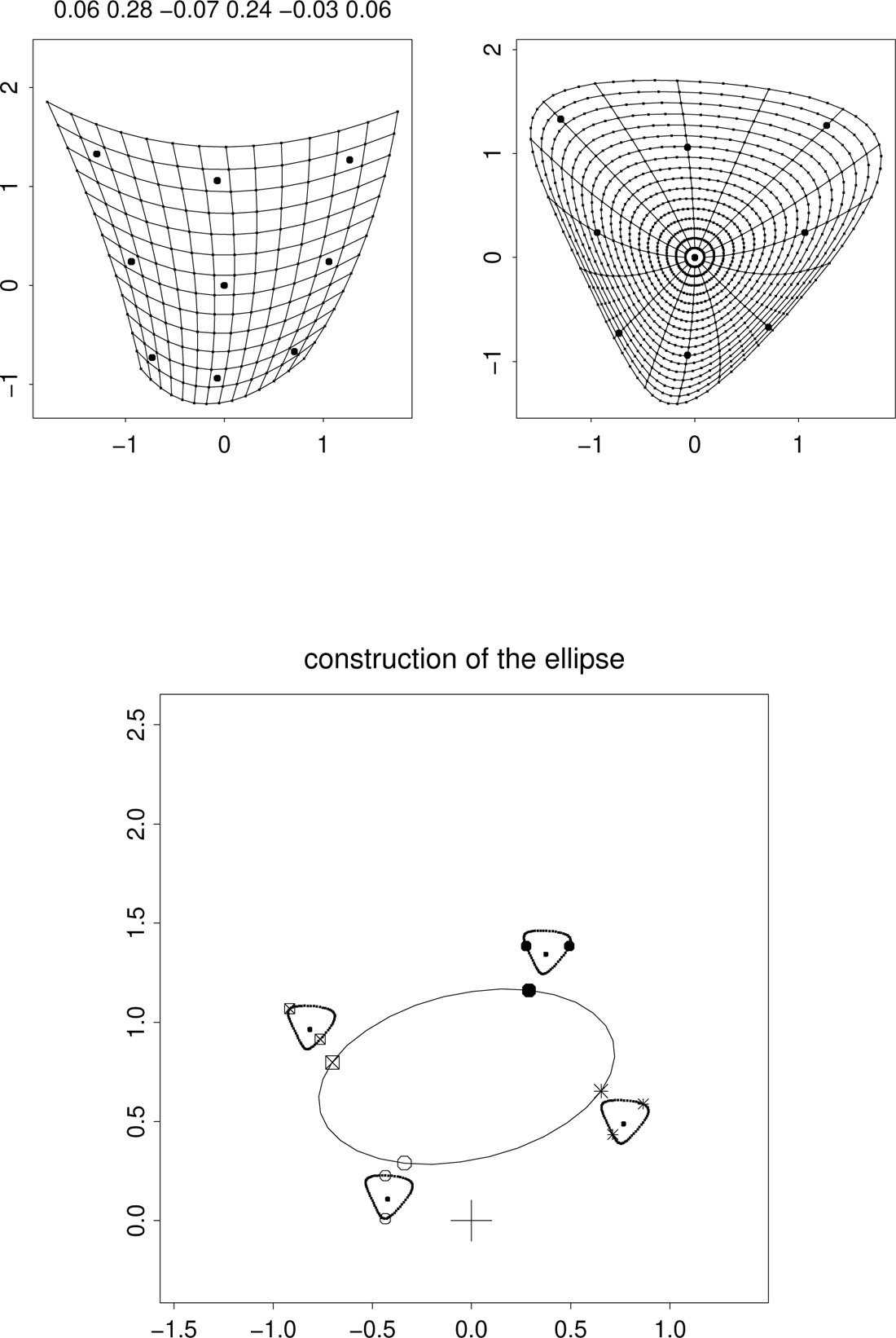
Schematic of the specimen-by-specimen analysis, here exemplified by a pure quadratic leaving the point (0, 0) fixed at the center of a 3 × 3 template. (above left) The target, an exact quadratic deformation of the template according to the trend with the coefficients printed above the diagram, represented in Cartesian coordinates over a 3 × 3 grid of cell size 1. (upper right) The same transformation now diagrammed in polar coordinates up through radius 1.5. (below) The ellipse of second derivatives of that quadratic trend, constructed as half the second differences of points at opposite ends of a diameter of the image of the unit circle with respect to the shared point at the origin, or, equivalently, the average of that pair of points with respect to the origin. The + marks the origin of coordinates.

Below is the conversion of the polar plot to the ellipse tracing all of its directional second derivatives. Around the enlargement of the deformed unit circle from the upper-right panel are vignettes of four of these halved second differences plotted with four different symbols as in Figure 3. Again these are the four *cardinal diameters* whose statistics will concern us in the examples of Sections III and IV. The figure clarifies the symbols that will be used to differentiate them: a bullet for the horizontal diameter, an open circle for the vertical diameter, an asterisk for the diameter running southwest-northeast on the template, and a times sign in a box for the northwest-southeast diameter at 90^◦^ to that one. Note how the second derivatives in the *x* and *y* directions lie at opposite ends of a diameter, and likewise those for the *x*+*y* and *x*− *y* directions, and that these diameters are what the geometer calls *conjugate,* meaning that each one is parallel to the ellipse’s tangent at the points of the other. (This property is the affine generalization of the situation for perpendicular radii of a circle.) See Figure 5 and its caption.

**Figure 5.**
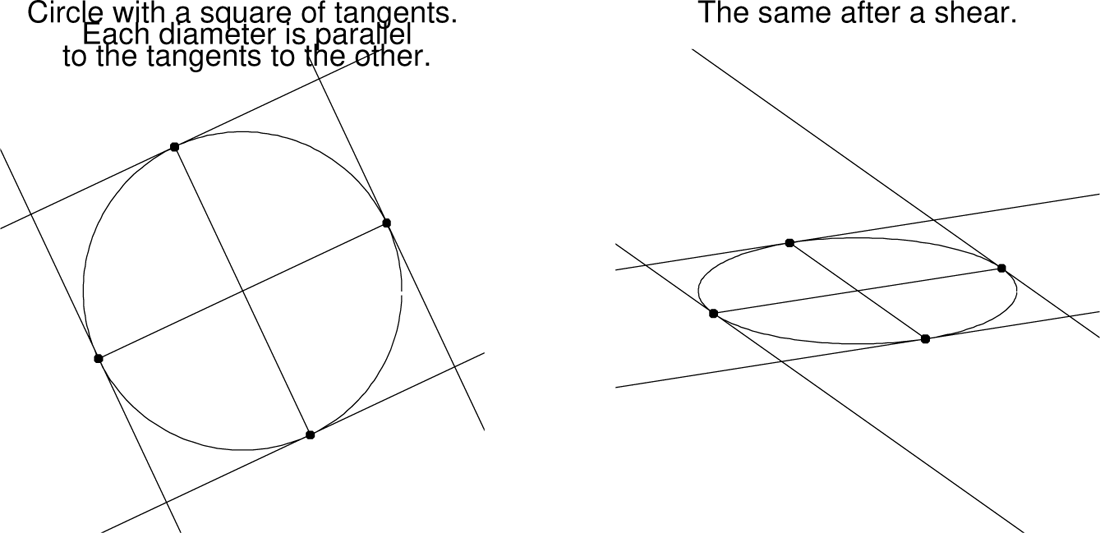
Conjugate diameters of an ellipse are pairs of diameters that were perpendicular in the circle from which the ellipse emerged as a shear. On the circle, each diameter is parallel to the tangents to the circle at the touching points of the *other* diameter, and this parallelism is unaffected by shear operations. (left) A circle with its circumscribed square. (right) The corresponding configuration after the circle is sheared into an arbitrary ellipse.

Put more abstractly: the six-dimensional feature space of quadratic trend grids of arbitrary landmark configurations over a common template is identical with the six-dimensional feature space of cardinal diameters of an ellipse inscribed on the same picture plane. The Euclidean formulas for distance are moderately different in those two spaces, but, obviously, neither one is “incorrect” — both are reasonable.

#### Trend formulas with just one or two terms

Because there are six free coefficients in formulas like (1) for the quadratic trends to be examined, it is worth drawing their effects singly and, more importantly, in pairs. Figure 6 uses Cartesian coordinates to show each single term twice, once with a positive coefficient and once with a negative coefficient. The more realistic Figure 7 switches to polar coordinates to show all of the possibilities involving *two* of these terms — several of the actual examples to follow in Sections III and IV will resemble one of these. The four panels here that look like stacks of circles with shifting centers correspond to projected images of a circular paraboloid, one of the standard quadric surfaces (Hilbert and Cohn-Vossen, 1932/1952).

**Figure 6.**
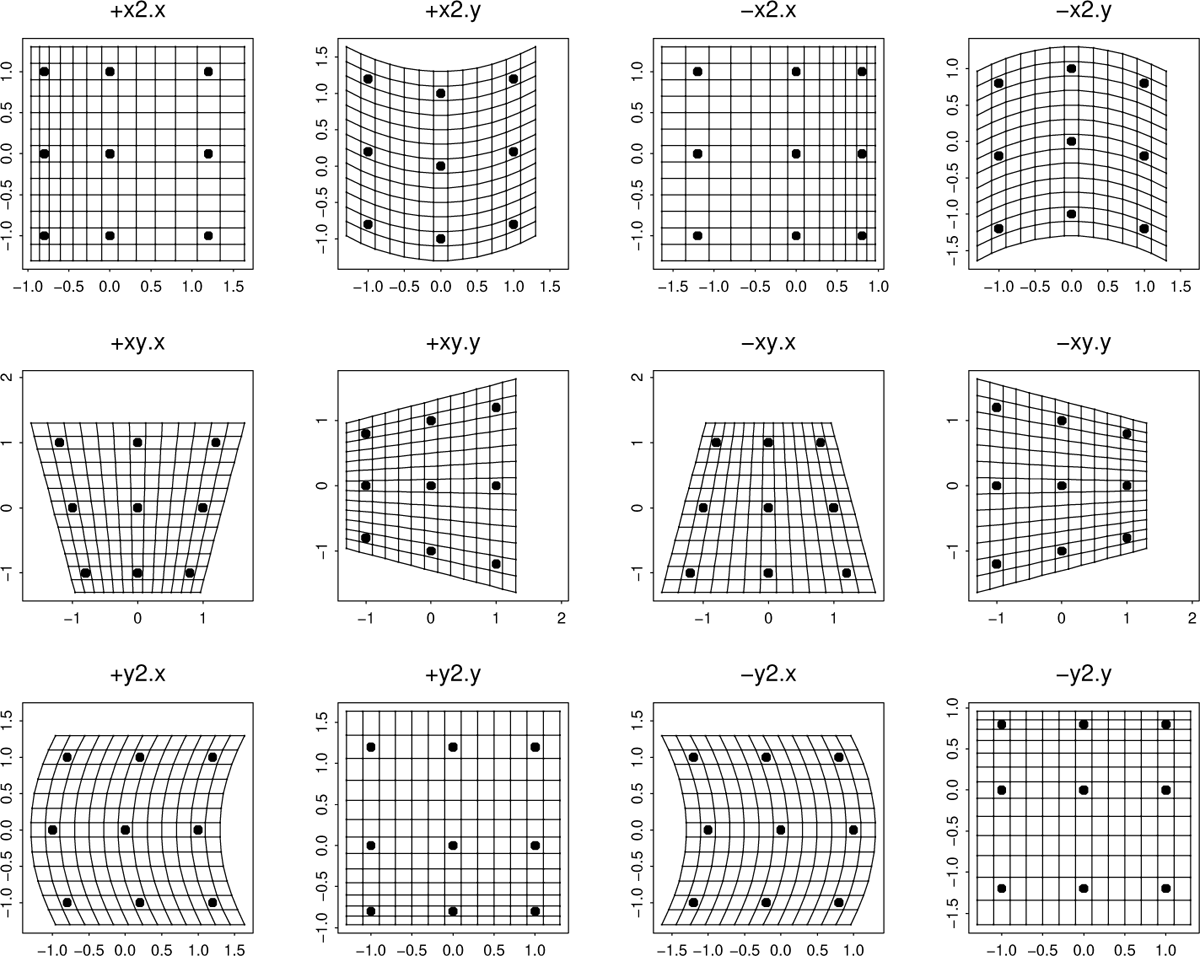
The six degrees of freedom of a quadratic trend fit, each plotted with both signs over a 3 × 3 template, in Cartesian coordinates. In the panel labels, “x2.x” stands for a regression term *rx*^2^ with *r* = 0.2 for the *x*-coordinate of a deformation, and likewise *y*2*.x* is a term *ty*^2^ for the deformed *x*; similarly *x*2*.y* and *y*2*.y* for *ux*^2^ and *wy*^2^; and finally *xy.x* and *xy.y* for terms *sxy* or *vxy* in one of the two coordinate regressions. Letters here correspond to terms in the regression formulas of equation (3) to come.

**Figure 7.**
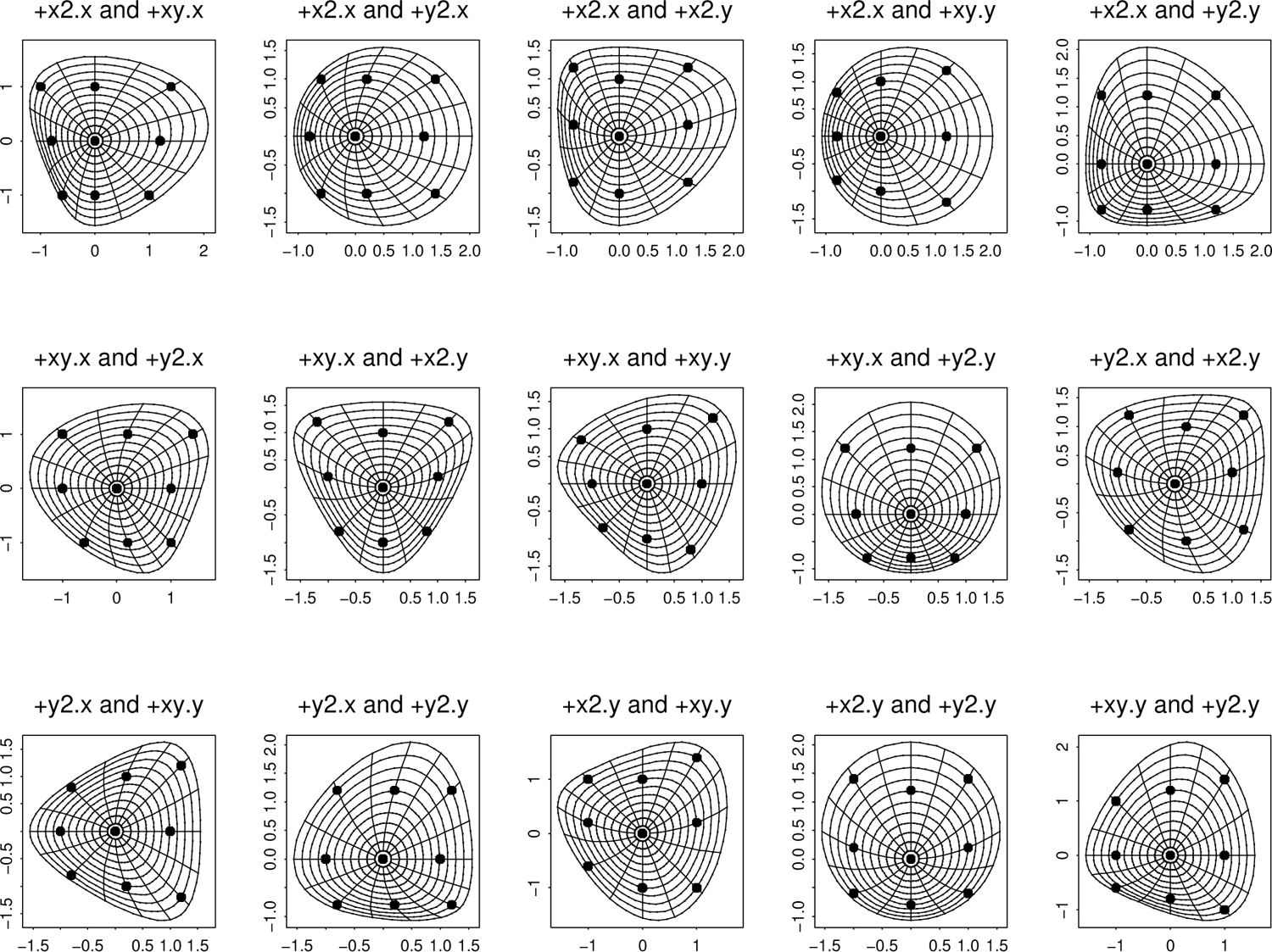
All pairwise combinations of the separate panels labelled with plus signs in Figure 6, now plotted more appropriately in the polar coordinate system of Bookstein (2022). Similar figures could be drawn for combinations of +−, −+, or −−.

For the analysis of the Vilmann growth process in Section III, we will need the rendering of the transformations of Figure 6 by the grid protocol of Figure 4. The resulting twelve “ellipses,” Figure 8, are actually line segments through the origin.

**Figure 8.**
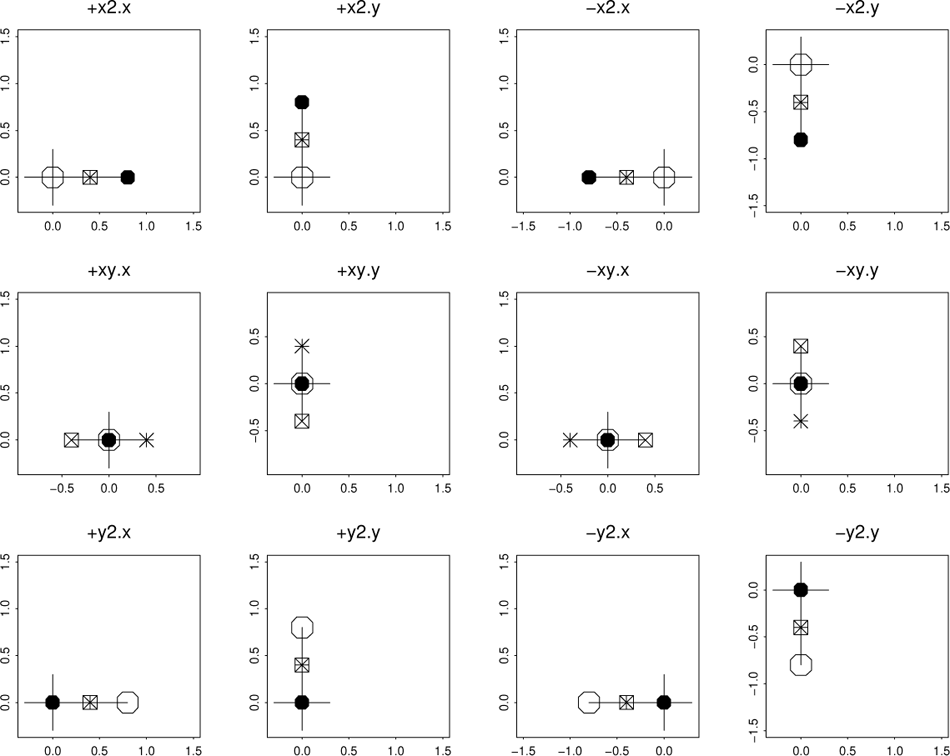
Single-term prototypes for the quadratic trend ellipses: the second-order derivative analysis for each of the frames in Figure 6. Plus sign: origin of coordinates. Other symbols show the second derivatives in the four cardinal directions as in Figure 4.

At this point we have already arrived at a diagram whose salient features can be used for reports of empirical findings wherever any *two* of the six coefficients of the quadratic trend map dominate the other four in absolute value. Figure 9 supplies such an ellipse for each two-coefficient panel in Figure 7. Those that are not points or circles appear as lines oriented at either 0^◦^ or 45^◦^ to the horizontal and vertical.

**Figure 9.**
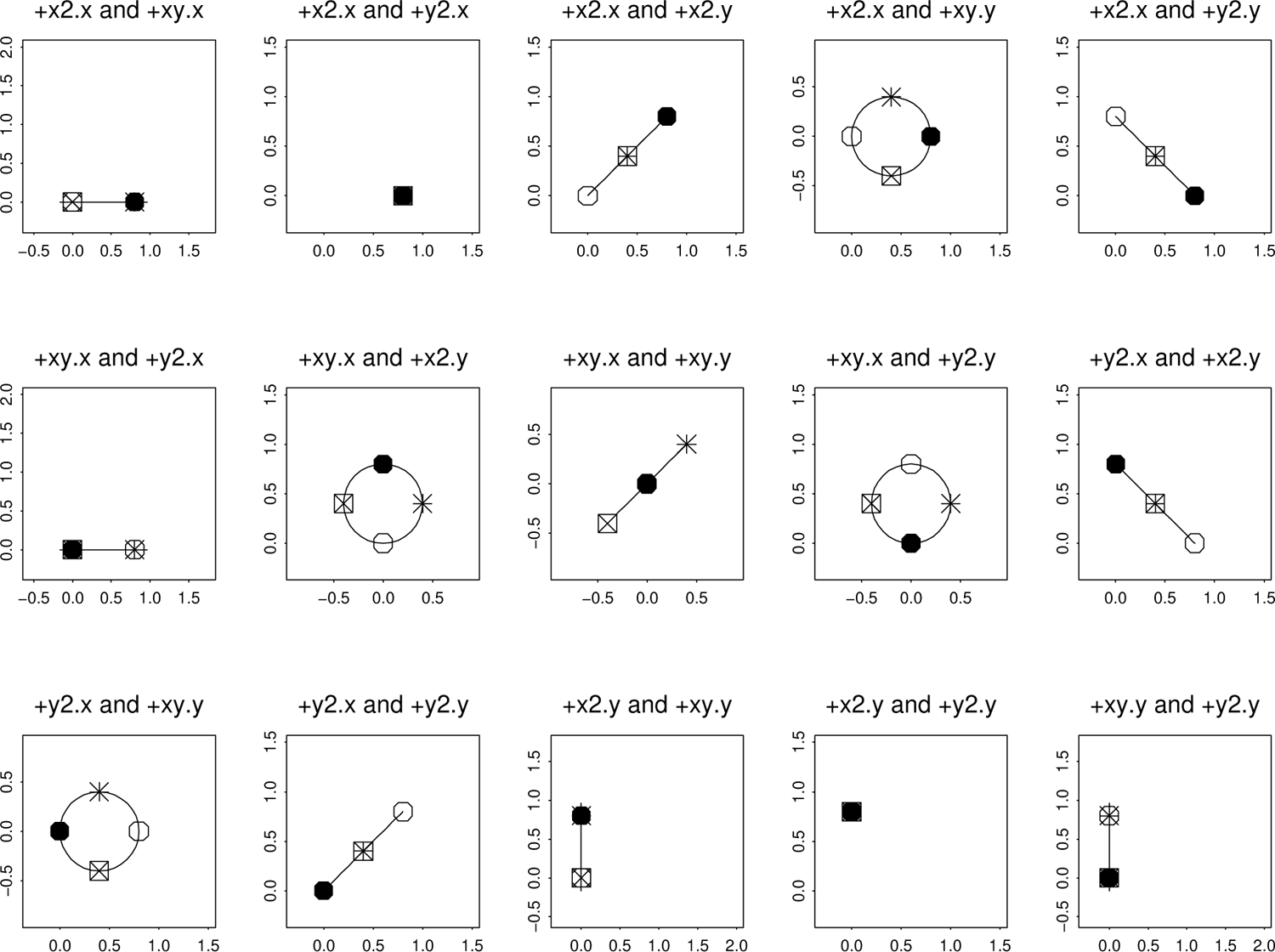
Ellipses for the transformation grids in Figure 7 are all either circles or line segments. As in Figure 8, the second directional derivatives in the *x*-direction and *y*-direction and their bisectors are plotted with the usual four symbols.

#### “Ellipses” are sometimes points or lines

Let us look a little more closely at Figures 8 and 9. In Figure 9, two of the frames display single points (overlaid with all four of the cardinal symbols). These are the frames labelled “+x2.x and +y2.x” and “+x2.y and +y2.y,” meaning, configurations with the coefficients of *x*^2^ and *y*^2^ in one direction equal to each other and all other coeffients zero. Why is this the case? The answer can be phrased either algebraically or geometrically. Geometrically, note in Figure 8 that the “ellipse” for the regression with single term *rx*^2^, upper left corner panel, is a line segment with the point for the *x*-direction at one end, the point for the *y*-direction at zero, and both points for the *xy* and −*xy* directions halfway between. For the analogous regression with just the term *ty*^2^, lower left corner, the ordination of directions is the same except the solid disk and the open disk are reversed. (The letters *r* and *t* here, as in the caption to Figure 6, correspond to terms in the regression formulas of equation (3) to come.) The sum of these two configurations replaces both of the disks by their average and leaves the other two symbols right where they are, but that is the same place as the average of the two disk symbols. Hence after averaging, all four symbols land in the same place, a point on the *y*-axis but not on the *x*-axis, as in row 1, column 2 of Figure 9. Algebraically, the transformation *rx*^2^ + *ty*^2^ with *r* = *t* sends all of the points (*x, y*) = ±(cos *θ,* sin *θ*) to *r*(cos^2^ *θ* + sin^2^ *θ*) ≡ *r,* independent of the angle *θ*.

Likewise one can confirm that combinations like the one with *r* = *s* and all other regression coefficients zero, upper left panel of Figure 9, yield “ellipses” that are lines with both of the second-derivative coefficients for the mixed derivatives at zero and the other two atop one another on the *x*-axis. There are four such combinations, versus two for the point-type of the previous paragraph. Geometrically, this is the sum of the labelled line segments in the first two rows of column one of Figure 8. Algebraically, these points are the averages of terms *r*(cos^2^ *θ* ± sin *θ* cos *θ*), where the second term cancels over the ± operation and the first one simply tracks the squared cosine function over its angular range.

But there is another way to get a straight line “ellipse” in this approach: any of the single-term regressions set down in Figure 8. There are two different types of this ordination, depending on whether the single term is the mixed expression *sxy* or *vxy* (row 2 of the figure) or instead one of the pure squares (row 1 or row 3). In the latter case, one of the disks is at the origin, the other disk is at some remove, and the symbols for the other two cardinal directions overlap halfway between. In the former case, both of the disks are at the origin, and the second derivatives for the ±45^◦^ directions lie on equal and opposite vectors.

The two pure types of ellipses, points and lines, combine, so that any of the lines in Figures 8 or 9 can be shifted to accommodate any point representing equal coefficients *r* and *t* or *u* and *w*. We will see examples of all these intriguing configurations in the next two sections.

For data in 3D, as I mentioned above, the sphere of directions would lead to a surface of second derivatives taking the form of an ellipsoid, not an ellipse, and analysis would proceed using diameters from all thirteen cardinal directions (three edge directions, six face diagonals, four body diagonals), not just the four of this presentation.

### III. A simple example: the Vilmann neurocranial octagons

The taxonomy of examples in Section II can serve as a typology of ideal types for the understanding of individual examples. This section does so for a familiar textbook data set, the “Vilmann neurocranial octagons” tracing around the midsagittal neurocrania of close-bred laboratory rats radiographed in the 1960’s by the Danish anatomist Henning Vilmann at eight ages between 7 days and 150 days and digitized some years later by the New York craniofacial biologist Melvin Moss. This version of the data is the one explored in my textbook of 2018: the subset of 18 animals with complete data (all eight landmarks) at all eight ages. The concern in this section is the contrast of the Procrustes-averaged shapes for the age-7 and age-150 animals (only the averages, no consideration of covariances).

#### Starting from a regression instead of from a model

The quadratic maps in Section II were all synthetic, in the sense that they illustrated an exactly quadratic correspondence

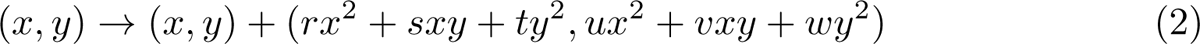

between two configurations of nine landmarks, one of which was an exactly Cartesian grid. This was the case even when the ultimate visualization, as in Figure 7 or Figure 9, was not itself Cartesian. We identified these coefficients *r* through *w* with half the second derivatives of the resulting mapping, but for any empirical study those coefficients need to be produced by some arithmetical manipulations based in the actual data. As Sneath suggested so long ago, that arithmetic is the standard least-squares analysis that applied statisticians in a great range of different disciplines exploit when it is adjudged sensible to “fit a model by least squares”: they are the coefficients *r* through *w* of the more highly parameterized *regression model* approximating the target configuration (*x*^′^*, y*^′^) as an exact polynomial function of the template configuration (*x, y*), the formula (*x*^′^*, y*^′^) = (*a* + *bx* + *cy* + *rx*^2^ + *sxy* + *ty*^2^*, d* + *ex* + *fy* + *ux*^2^ + *vxy* + *wy*^2^), and our task is to minimize the sum of squares of discrepancies of this predictor with the target locations (*x*^′^*, y*^′^) over the template configuration: minimizing

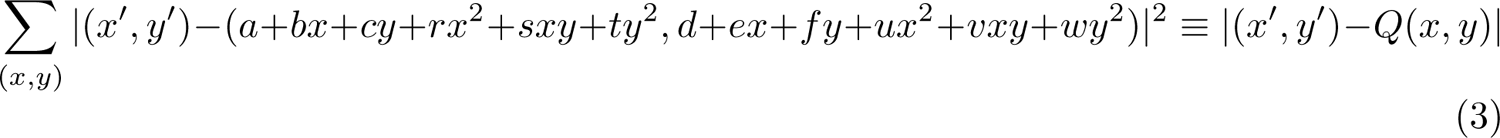

(this will be our definition of *Q*) in which all twelve coefficients are calculated to minimize the sum of squared lengths of the error term in the complex plane. Each coordinate is then itself the result of an ordinary multiple regression *x*^′^ ∼ *a* + *bx* + *cy* + *rx*^2^ + *sxy* + *ty*^2^, etc. In the examples of Sections III and IV we ignore the values of the constants *a* through *c* (and likewise the *d*, *e*, *f* that characterize the analogous regression for *y*^′^), examining only the coefficients of the quadratic part, which, according to the geometry of Section II, can be interpreted unambiguously as half the second derivatives of the fitted quadratic trend: at every point of the picture plane, we have 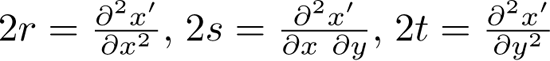, and similarly for the second partial derivatives *u*, *v*, *w* of *y*^′^.^1^ The calculus of the complex plane allows us to combine these two ordinary multiple regressions into the one quadratic trend analysis in two dimensions minimizing the sum of *both* families of squared errors, the one for *x*^′^ and the other for *y*^′^, because of the Pythagorean mystery that what we perceive as distance on the picture is actually the square root of the sum of these two squared arithmetical differences. (This observation is certainly not original; it is already explicit in Sneath’s paper of 1967.)

Near the end of the Discussion, Section V, I will return to this convenient equivalence. Until then, it is simply assumed that it makes biological sense to consider the parameters *r* through *w* to be sensible quantifications of what the biologist’s eye would already see as one meaningful aspect of a composite characterization of the difference in form of two organisms, each as represented for the purposes of that comparison by a configuration of finitely many landmarks. When landmarks are closer together, which is not the case for this example, one may think of the quadratic regression as a specialized version of a *smoothing* — a projection of the *x*− and *y*−coordinates of every landmark configuration on the five predictors *x*, *y*, *x*^2^, *xy*, *y*^2^ of the template. It is not the ordinary sort of smoothing of an image, convolution with a Gaussian, but a representation within one shared specifically smoothed subspace of second derivatives all constant.

#### Rotating coordinates helps interpretation

Under the assumption that landmark configurations can yield meaningful sets of coefficients *r* through *w*, Figure 10 begins at upper left with one version of the conventional analysis of this growth analysis, the straightforward graphical comparison of the Vilmann octagon averages at ages 7 and 150 days. From the quadratic fit (open disks) of the trend’s deformation of the age-7 average configuration to the age-150 average (filled disks) it is clear that the trend method has captured most of the relevant geometric signal here; the problem is rather to state in simple words and coefficients what the meaning of that signal actually *is.* (Analogous grids could have been produced using principal component 1 or the regression of the octagon’s shape coordinates on Centroid Size — see, in general, the range of contexts of this example in Bookstein 2018 — but the thrust in this section is the interpretation of the single comparison of one pair of averaged configurations.)

**Figure 10.**
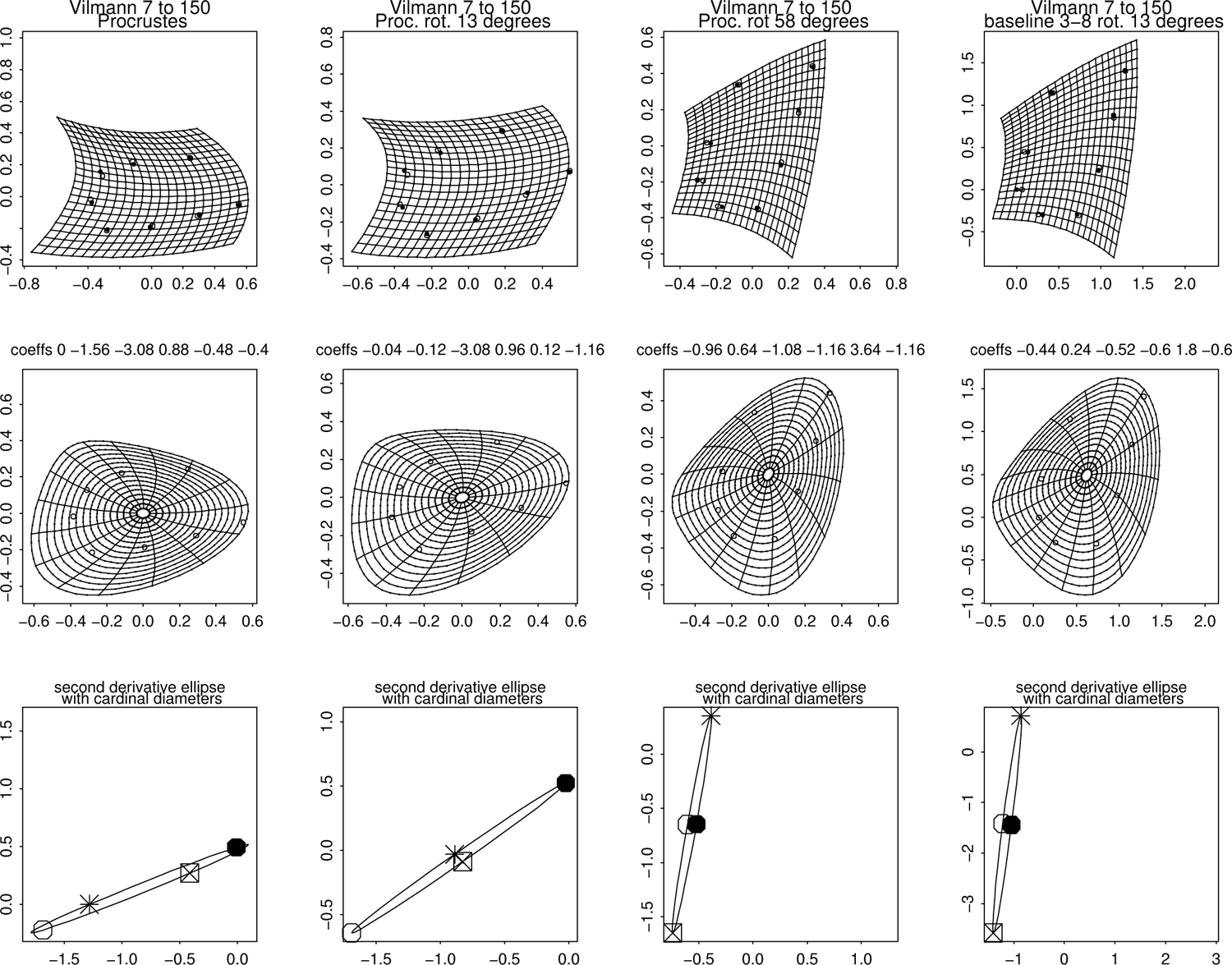
Analysis of the growth of the Vilmann neurocranial octagons averaged over the usual sample of 18 laboratory rats aged 7 days to 150 days. Columns, left to right: conventional Procrustes pose, rotation by 13^◦^ to superimpose the second derivatives along the diagonal directions (1, ±1), rotation by 58^◦^ that superimposes the second derivatives *∂*^2^*/∂x*^2^ and *∂*^2^*/∂y*^2^ along the coordinate axes instead, and rotation by 13*^c^irc* of a two-point registration (landmark 3 to landmark 8, Interparietal Point to Sphenoöccipital Synchondrosis) yielding exactly the same interpretation but without any use of the word “Procrustes” or any of the corresponding formulas or algorithms. Rows, top to bottom: Cartesian representation of the quadratic trend fit, polar-coordinate rendering of the same, and the second-directional-derivative ellipse (always the same shape) with the four cardinal diameters highlighted. In the upper two rows, the locations to which the points of the age-7 template are deformed by the grid are plotted in open circles; in the top row, their actual age-150 averages are shown as well by the solid circles.

The ellipse here, bottom left in the figure, appears to be close to a special case, as it has essentially only one dimension of variation — the minor axis is of length close to zero. We are free to rotate the coordinate system so that that minor axis falls on a meaningful diameter of the coordinate system in which the trend analysis was couched: explicit variation of the orientation of the Cartesian system used to convey the trend. Column 2 of Figure 10 offers one alternative, a rotation of 13^◦^, for which that null diameter connects the second derivatives at ±45^◦^ to the baseline. From the formula (25.3.26) of Abramowitz and Stegun 1964 it follows that the mixed second-order partial derivative of this quadratic trend is close to 0 for both the *x*− and the *y*-coordinates of the target configuration: the map is just a superposition of two processes each looking like any of the diagonal-dominant frames in Figure 9. (We can detect the dependence on just *x*^2^ and *y*^2^ in the top row, second column, of Figure 10, where neither system of grid lines is distinguishable from parallel translations of the same parabolic curve.)

That rotation zeroed the mixed partial derivatives. A different rotation, at 45^◦^ to that one, will shift the vanishing diagonal of the ellipse from the diagonal canonical diameter to the cardinal diameter with *∂*^2^*/∂x*^2^ equal to *∂*^2^*/∂y*^2^ for both dimensions of the target configuration. In this representation, furthermore, the ellipse has rotated close to orientation with a different, equally salient ideal type: it is nearly aligned with one of the coordinate axes of the plot. And in yet *another* potential special case, the uppermost point of this ellipse, for the second derivative in the (1, 1) direction, is close to the (0, 0) of this diagram. After rotating 45^◦^ to sum-and-difference coordinates, then, we find ourselves close to the situation in the lower right panel of Figure 9, an “ellipse” that is just a line, horizontal or vertical, anchored at the origin of its coordinate plane. An appropriate summary of this finding’s dominant feature would thus concentrate on that single mixed derivative. The situation (Figure 11) is the one I described decades ago (Bookstein, 1985) as the *bilinear map* leaving two families of straight lines straight. (The linear term of this trend fit cannot alter the straightness of those lines, although it may well modify their angle from the 90^◦^ characterizing their relation to the highly symmetrical template of a square.)

**Figure 11.**
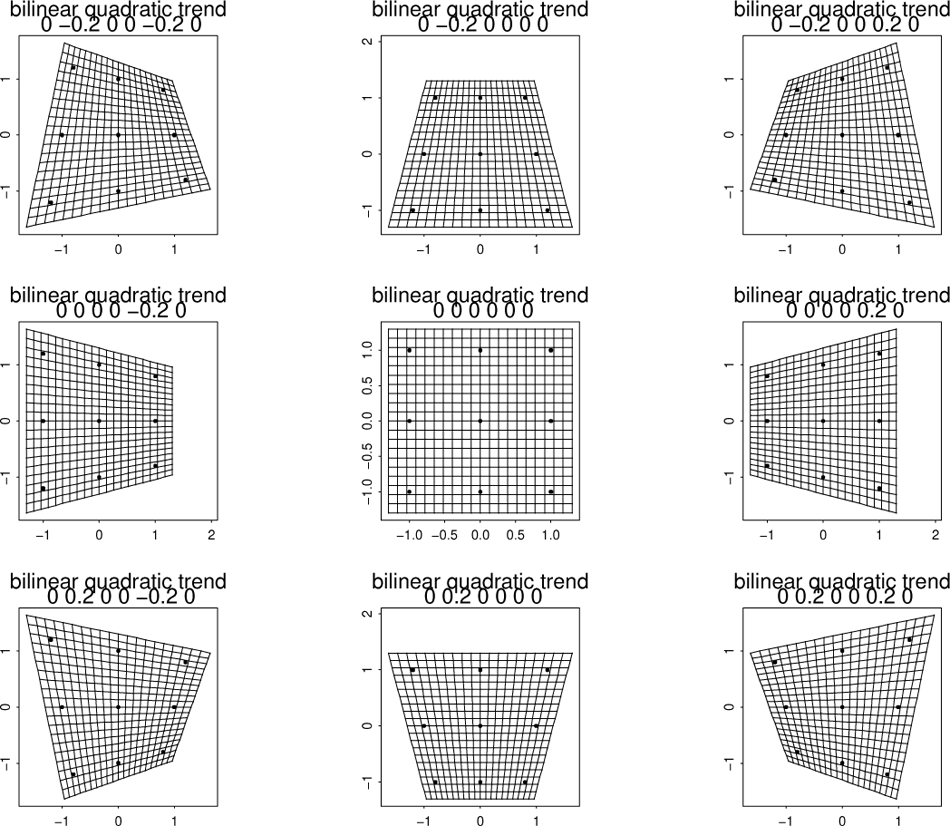
The *pure bilinear trend* of Bookstein (2023) is the quadratic trend of this paper with parameter string (0, ±.2, 0, 0, ±.2, 0) as in any of the corner instances here.

The bilinear map has an unexpectedly simple verbal report: opposite boundary segments are transformed linearly and the mapping deforms intersections of proportional transects across the template quadrilateral to intersections of the two new sets of segments connecting proportional aliquots of the target quadrilateral. Hence the map transforms two sets of straight lines on the template (in Figure 11, a square) into two other sets of lines that are likewise straight (but no longer parallel) on the target image. Most other straight lines map into parabolas. The grid at upper right in Figure 10 is close to the prototype in the second row, third column of Figure 11.

Note that the analysis in Figure 10 involved no thin-plate spline, nor did it rely on any details of the Procrustes analysis driving the configurations of the leftmost three columns in the top row. The finding is unchanged except for a translation and rescaling of the ellipse when, imitating the analysis in Bookstein (2023), we abandon the Procrustes framework for a two-point (Bookstein coordinate) representation as in the fourth column. The Procrustes procedure per se added nothing to the biological interpretation here, and in fact it seriously interfered with the interpretation of the finding, inasmuch as freedom to rotate the coordinate system of the reporting grid is crucial to understanding the deformation.

But how is that rotation to be described? Calling a pose “58 degrees rotation from Procrustes” is not helpful when that Procrustes pose itself bears no biological reference: such a reference position, conventionally aligned with the first principal axis of the landmarks of the template, is not accessible to the biologist’s intuition. In contrast, the figure’s description of the identical pose as 13 degrees from a specific interlandmark segment is a clear instruction. In terms of the prototypes in Section II, the analysis here is closely aligned with a combination of just two frames: for the vertically extended ellipse, the combination in row 2, column 2 of Figure 8; for the left shift of all those second derivatives in the *x*-direction, column 2 of row 1 of Figure 9.

#### Effect of baseline choice

The analysis in the rightmost column of Figure 10 rests on a seemingly arbitrary choice of baseline for the two-point construction: the segment from the Interparietal Point to the Sphenoöccipital Synchondrosis. (This was the ultimate recommendation of the earlier analysis in Bookstein 2023.) From Figure 10 we see that the ellipses of interest are invariant in size and shape, but only rotate with the coordinate system. That is reassuring, but it is more important to see the extent to which the analysis is stable against changes in the selection of the pair of points against which the baseline is constructed. Figure 12 continues the reassurance by superimposing those second-derivative ellipses for nine more different baselines, as shown in the inset diagram: not only Interparietal to Sphenoöccipital but also every segment linking one of Basion, Opisthion, or Interparietal Point to one of Bregma, Sphenöethmoid Synchondrosis, or Intersphenoidal Synchondrosis. At the top are drawn all ten of the resulting ellipses after they are rotated into the orientation at right in Figure 10. The plus sign is the origin of this plot, which is even more favorably placed than the example in Figure 10, i.e. closer to zeroing the second derivative in the *x*-direction. In the middle, the same cardinal diameters are displayed without the elliptical arcs connecting them, using the usual symbols for all except *∂*^2^*/∂y*^2^, which is labelled instead by the pair of landmarks serving as the baseline. Clearly the points for *∂*^2^*/∂x*^2^ are tightly clustered, and also those for the (*x* + *y*) direction. Those for the second derivative in the *y* direction or the (*x* − *y*) direction show more scatter on this plot but the scatter would not affect the interpretation of the growth pattern as bilinear. The baseline used at right in Figure 10 is the one numbered “38” here, which appears near the middle of the distribution of the *∂*^2^*/∂y*^2^ points of these ten alternatives.

**Figure 12.**
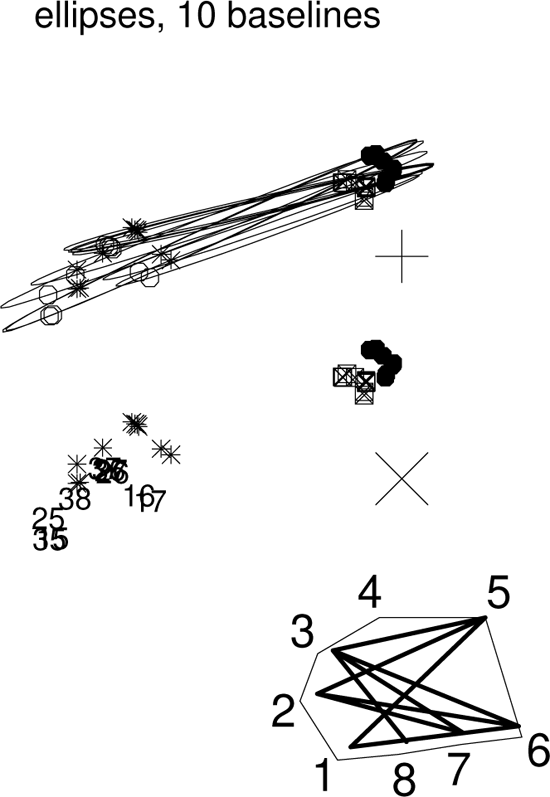
When rotated back into the original digitizing coordinate system, second-derivative ellipses of the longest baselines align with one another extremely well. (top) The quadratic trend ellipses to the ten selected baselines. with the standard four directional second-derivative symbols from Figure 4. Big plus sign, (0, 0), the origin of coordinates (all second derivatives zero). (middle) The same with ellipses suppressed and the *y*-direction second derivative point replaced by its two-digit baseline code. To avoid confusion the big plus sign is replaced by the big × sign. (bottom) The ten baselines: every join of one of the landmarks 1, 2, 3 to one of the landmarks 5, 6, 7 in the geometry of the average age-7 configuration, along with the 3-8 baseline from Figure 10.

Evidently when analytic results like these are rotated back into an appropriate digitizing coordinate system, second-derivative ellipses of the longest baselines align extremely well. Put another way, the Procrustes rotation has no scientific meaning — any requirement that this (or any other) orientation be standardized as part of a standard GMM dataflow makes no biometric sense. In a better toolkit, rotation would not be standardized, but instead those analyses will be highlighted which, like the quadratic trend ellipses here, do not depend on rotation — for which the rotation does not much affect the arithmetic of findings but so substantially affects the cogency of their reports. It is not that the Procrustes method disagrees with these versions – its ellipse, in column 1 of Figure 10, agrees with these. But nor did that Procrustes orientation gain us anything over the registration Moss originally applied to Vilmann’s images forty years ago. We want analyses for which Procrustes orientation, or any other orientation prior to analysis, is *irrelevant* to the reportage. That frees us to explore the grammar of the template coordinate grid perse, which can be a crucial component of a biological interpretation.

### IV. A realistic example: revisiting a mammal cranial data set

The data set on which Figure 1 is based is the 13-point midsagittal subset of a 35-landmark cranial configuration that arose as a revision of a data set originally exploited in Marcus et al. (2000). Its 55 specimens, all but the dolphin and the hyena from a representation kindly sent to me by Erika Hingst-Zaher in 2013, omit the deer-pig Babirusa from the 2000 sample but insert specimens of Kangaroo, Man, Sheep, and Boar. (The 2000 article also notes that the selected Elephant skull, along with those of Walrus and Manatee, had to be a young one in order to fit into their digitizing apparatus.) Figure 13 scatters the data in diverse analytic layouts. This particular revised data set was previously analyzed in Bookstein (2018, 2019) by a method different from the approach put forward here. In the upper right panel, the count of three partial warps the residualizations from which are scattered is set to correspond to the six degrees of freedom for the analysis by quadratic trend in the panel below it. Each partial warp score involves two degrees of freedom because in the GMM formulary it is a complex number. For an introduction to this way of notating the Cartesian plane see pages 370–371 of Bookstein 2018, or, for a much deeper pedagogical guide, Chapter 2 of Mumford et al. 2006.

**Figure 13.**
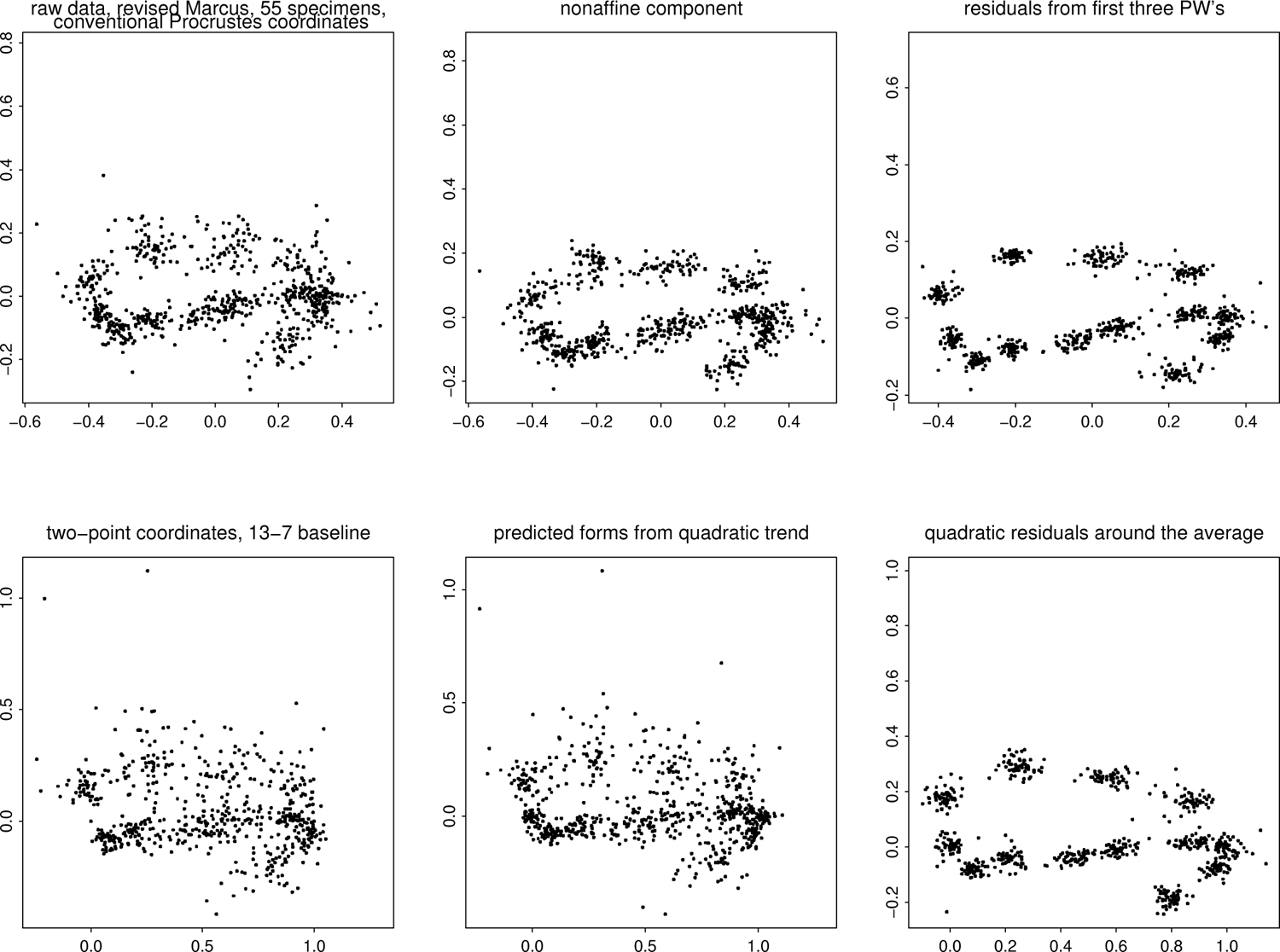
A selection of scatterplots relevant to the information in Figure 1 and those following in this section. All panels pertain to the distribution of the 13-landmark configurations for the 55 representatives of mammalian orders diagrammed individually in the figures of the Supplement. (upper left) The conventional Procrustes coordinates for these 55 13-landmark configurations. (upper center) The nonaffine component of that Procrustes scatter, adjusted for the linear (affine) aspect of their variation around the average from Figure 1. (upper right) Residuals from the first three partial warps of the conventional Procrustes-spline toolkit. (lower left) The two-point coordinates of the same raw data set to the longitudinal baseline used in Figure 1 and later figures. (lower center) Predicted forms from the quadratic trend analysis, showing roughly as much variation as the original data at their left. (lower right) Residuals from the quadratic trend analysis of this paper.

> *A note on notation.* When, as in this example, a large sample of target forms is to be regressed on the same template, the regressions can be written out all at once in a compact matrix expression of what the packages call a “multivariable regression.” But that notation, while useful for the programmer, does not simplify the exposition case by case — unlike the situation for principal components analysis, writing out all the shape coordinates of all the cases in one data matrix is not an insightful step toward understanding how their quadratic trends produce ellipses from the circle of directions — and does not substantially aid the task of visualizing the individual polar-coordinate grids and parameterizing the second-order ellipses at the core of the new methodology. Nor does the matrix notation help in the extension mentioned in Section V to phylogenetic contrasts, where the template would be different at every branch point of the phylogeny. So that alternative notation is not written out here; experienced R coders can construct it straightaway from equation (3) for themselves.

#### The four-panel dashboard

The Vilmann example of Section III involved only one single transformation, from the 7-day average form to the 150-day average. To extend this approach to larger samples it is helpful to have a formal dataflow, as laid out in the dashboards to follow. For the main text I have selected one exemplar (Bear) near the average of these configurations, another (Ondatra, muskrat) near one of the extremes, and a third one, Man, of parochial interest to most readers. For the full set of all 55 of these, please consult the Supplement. To understand the layout of any of the 55, review the three examples to follow here.

The key to Figures 14 through 16 is as follows. At upper left is a conventional Cartesian plot of the quadratic fit to the specimen from the average in Figure 1, with the coefficients *r* through *w* of the formula *Q*, equation (3), printed above. Solid dots, observed shape coordinates; open circles, predictions of the quadratic fit (including linear terms that do not contribute to the second derivative). At upper right the same deformation is rendered in polar coordinates around the centroid of the template. This grid is trimmed to avoid large expanses devoid of landmark data. At lower left is a close-up of the polar deformation within which is traced the warping from a circle of radius 0.5 in the template; the differences of averages of ends of diameters from the center here are the loci that, taken all together, comprise the ellipse at lower right. Cardinal directions of these diametral comparisons are keyed as in Figure 4. As none of the ellipses here are centered at (0, 0) except by accident, each diagram marks the origin of its coordinates (the purely linear transformation) by a large + sign.

**Figure 14.**
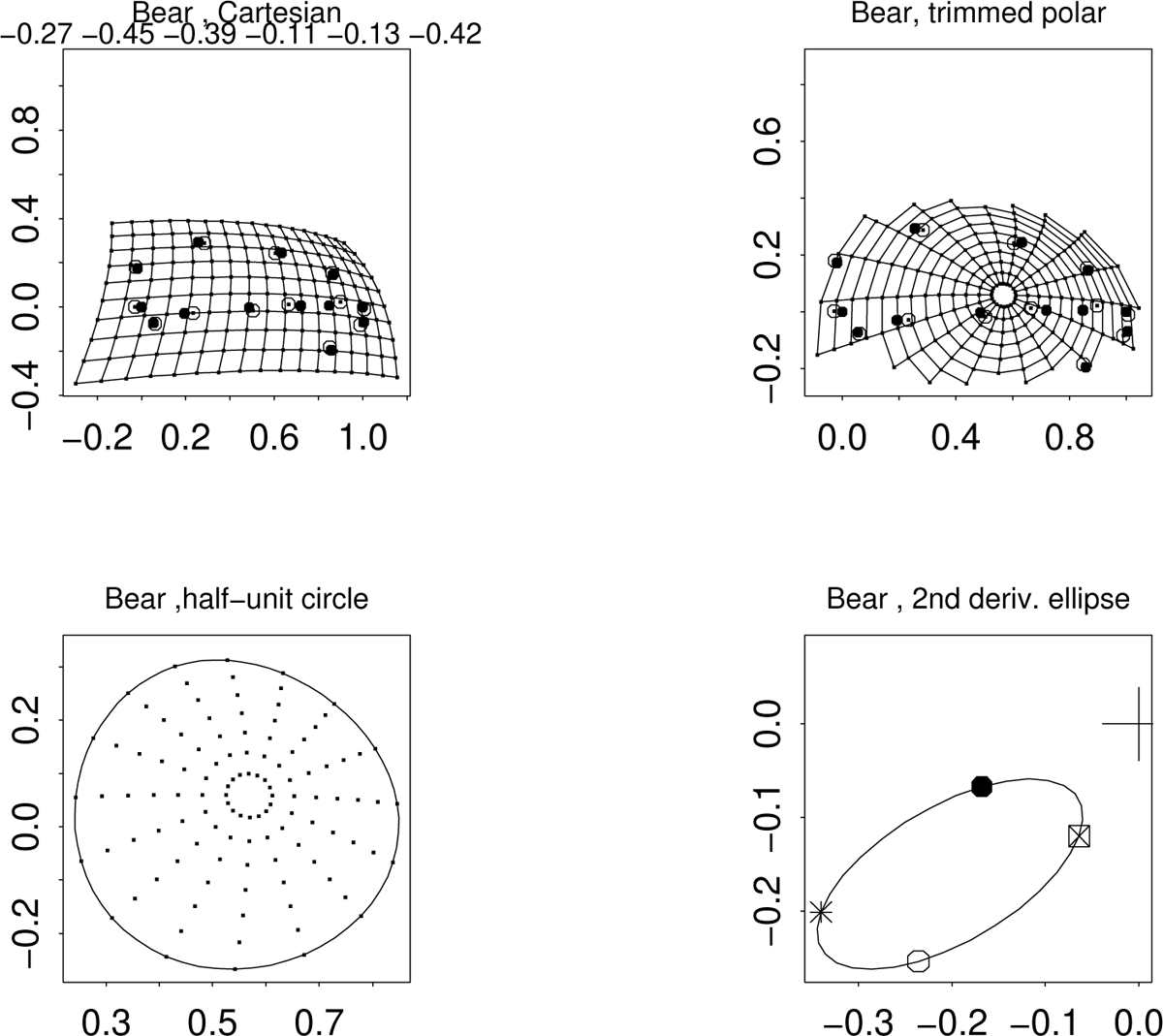
Dashboard for the quadratic trend fit of the average two-point shape coordinate configuration (Figure 1) to the Bear configuration, at a summed squared distance of 0.028 (fourth-closest) to the grand average in this data set. Comparing the axis scales of the lower-right panel here to those of the same panel in the next two figures, we infer that the ellipse here is not far from the prototype of one single point discussed in connection with Figure 9: not much quadratic warping at all.

**Figure 15.**
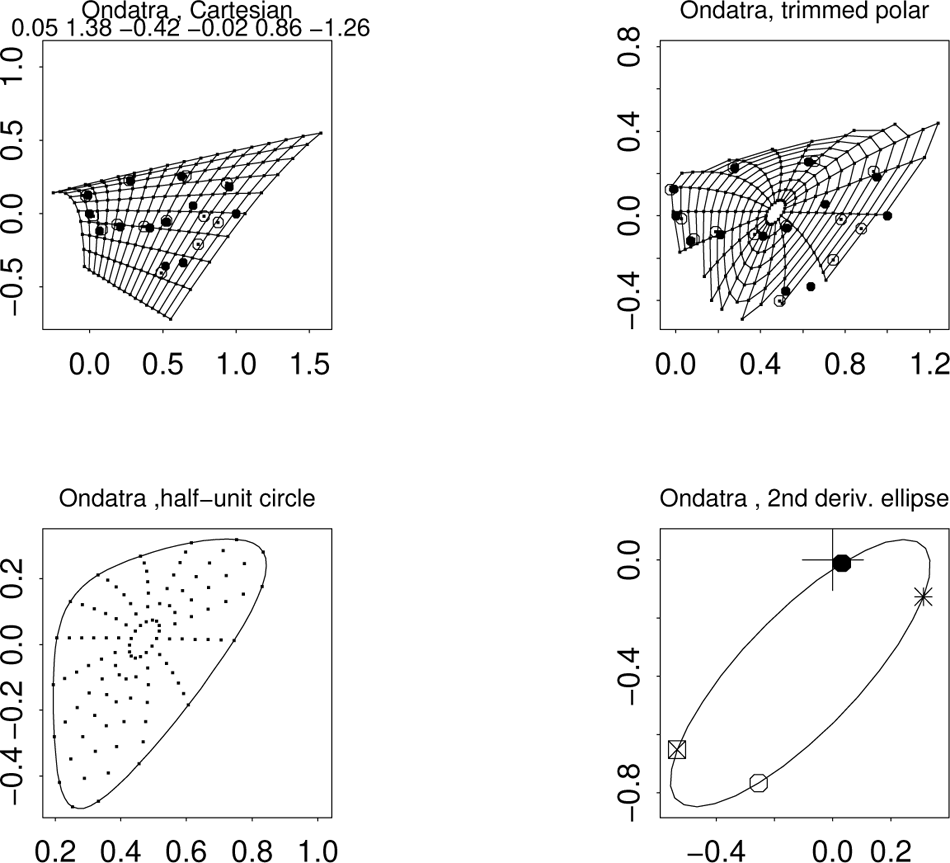
For the Ondatra (muskrat), distance 0.34 (fifth-largest) from the full sample average. Obviously the closed curve in the panel at lower left is not an ellipse, but still, as in Figure 3, the curve in the lower right panel is.

**Figure 16.**
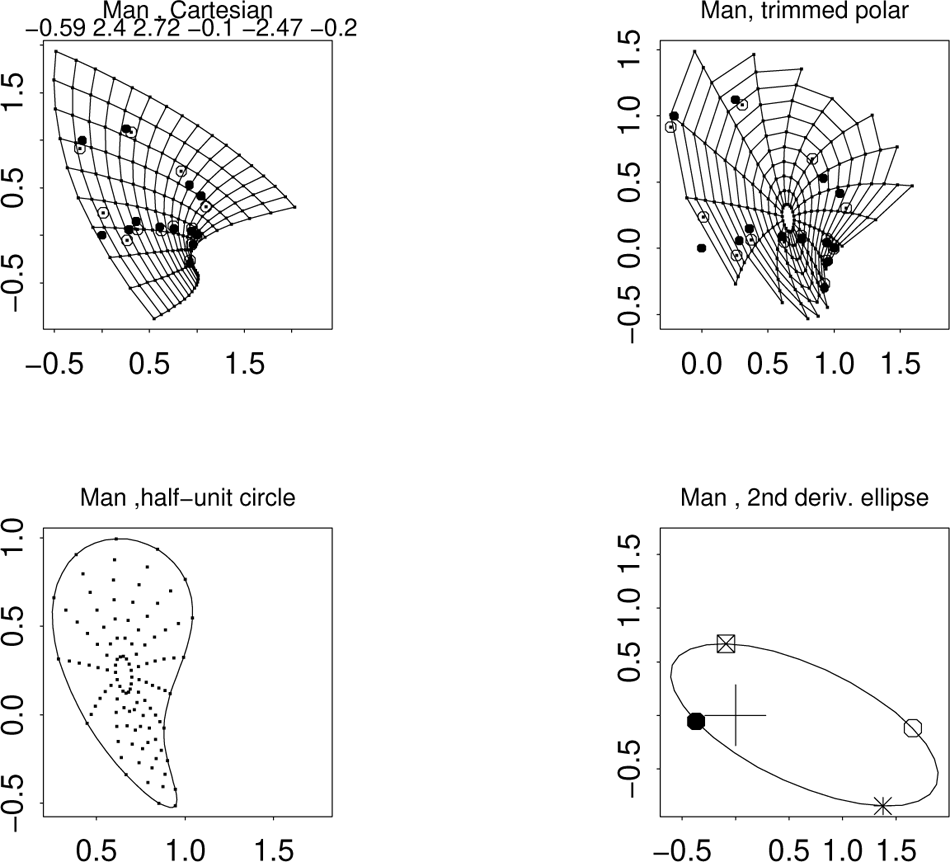
For the human specimen, farthest from the average at sum of squares 1.90, owing mostly to the extreme positions of the inion and the frontal-parietal suture here. The behavior of this second derivative in the vicinity of the mandible is interesting; it will concern us in more detail in Figure 23 below and in the “teardrop” analysis of Section V.

#### Reporting ellipses by their cardinal diameters

As rendered in Figure 1 the ellipses of Section II capture five of each quadratic fit’s six degrees of freedom (three coefficients for the horizontal shape coordinate, three more for the vertical). Here in Figure 17, which enhances a subset of the information in the right-hand panel of Figure 1, nineteen of the 55 ellipses that lay at or near the margin of the superposition there are named and the sixth degree of freedom indicated by symbols for the cardinal directions corresponding to the four directional second derivatives specified in the figure legend, icons already introduced in Section II and Section III. For each specimen, as explained in Section II, these directional derivatives apply across the entirety of the fitted quadratic grid. Directional derivatives along the baseline and at 90^◦^ to the baseline lie at opposite ends of a diagonal of their ellipse; the derivatives along the ±45^◦^ directions lie on the conjugate diagonal.

**Figure 17.**
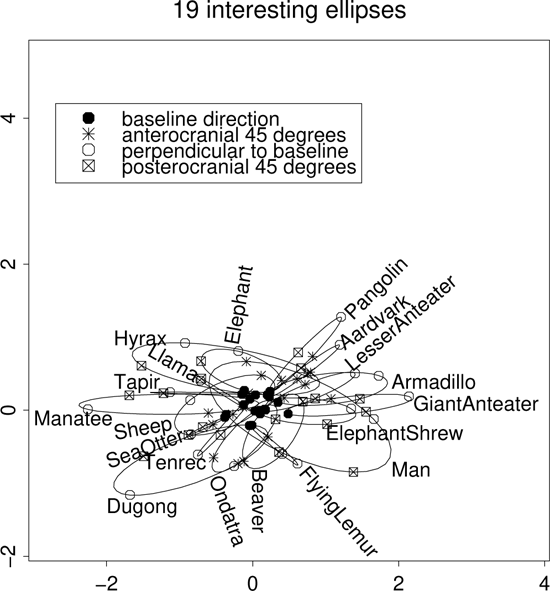
Nineteen ellipses from Figure 1 (right) that lie at or near the margin of the distribution there. See text.

Figure 18 conveys the same information less redundantly, by eliminating the elliptical curves entirely, leaving only the 19 pairs of cardinal diameters introduced in Section II. Like the vector of six coordinates introduced above in equation (1), these points bear a total of six degrees of freedom (since the midpoints of the two pairs characterizing each specimen must occupy the same location — a total of two linear constraints). The second derivatives along the baseline (filled circles in Figure 18) are tightly clustered around (0, 0) because the average shape of this configuration is highly elongated in this direction. The second derivatives in the perpendicular direction (open circles), derivatives vertical in this two-point system, are much more widely scattered. We shall see that the ordination of ellipses (trend gradients) here is moderately invariant to the choice of that baseline among reasonable alternatives. Nevertheless the diversity of these ellipses — the diversity of directional second derivatives of a quadratic trend fit unchanging in its template and in its least-squares formulation — is both novel and remarkable, and may be telling us something important about the evolvability of cranial form across Mammalia.

**Figure 18.**
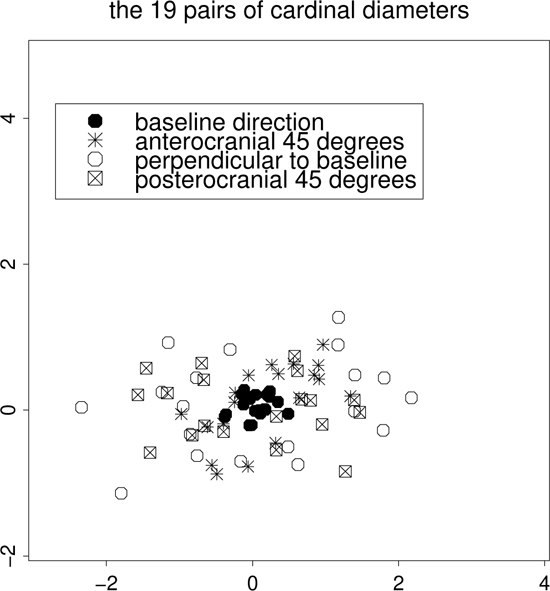
An even less cluttered rendering: just the cardinal diameters from Figure 17.

#### Principal components of coefficients; of diameters

To explore the diversity of these quadratic trend fits it is useful to begin with the complete display of their six-dimensional space. Figure 19 presents a view in the form of three two-dimensional projections each pairing one of the three coefficients — for *x*^2^, *y*^2^, or *xy* — for both of the Cartesian coordinates of this midsagittal plane. Plainly the lists on these three panels of the extremal forms are distinctive, in fact, nearly nonoverlapping: for *x*^2^, Baboon, Giant Anteater, Echidna, and Ornithorhyncus; for *y*^2^, the two cetaceans and again the Giant Anteater; and for *xy* the nearly collinear series Hyrax–Sheep–Man facing Beaver at the opposite corner of the scatter. We will shortly see echoes of all these positions in the multivariate analyses to follow.

**Figure 19.**
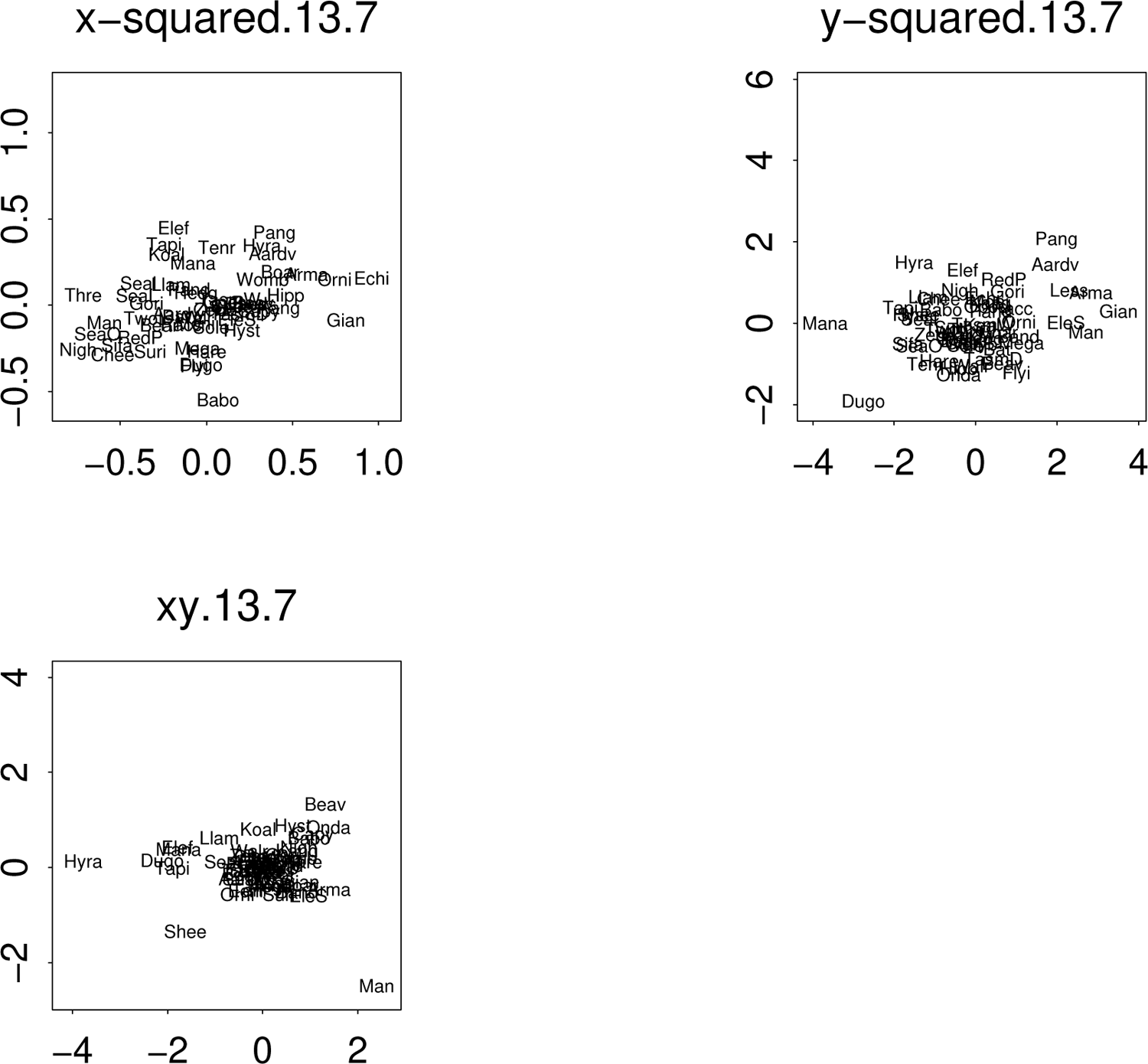
Scatters of the six dimensions of quadratic trend coefficients in three pairs, one for each of the patterns in Figure 7. Specimen names are the first four letters of their long strings as in the Supplement, except for the two beginning with Aard (five letters each) and the following four: TasmW, TasmanianWolf; TasmD, TasmanianDevil; Elef: Elephant. EleS: ElephantShrew. Note the difference in scale between the x-squared panel and the other two.

Figure 20 deals with diverse multivariate statistics of this new ordination by quadratic trend fits. The upper pair of ordinations, trend coefficients versus cardinal diameters, is aligned by canonical correlations .9809 and 0.8741, and the seven-species convex hulls of their scatters are identical as lists and nearly identical as shapes up to an affine transformation. But neither is acceptably in alignment with either of the Procrustes versions below: canonical correlations with the first two dimensions of the cardinal-diameters version are 0.8417 and 0.5779 vis-á-vis the full Procrustes shape coordinate space and 0.9018, 0.3576 with the nonaffine subspace there, which is ostensibly aiming at the same goal. Indeed the principal components (PC’s) of the canonical diameters are badly misaligned with their analogues in the Procrustes nonaffine setting — the correlation matrix of the two versions of the first three PC’s is 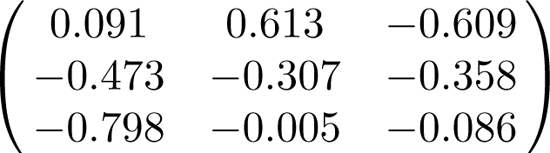. In the score plots, note, for instance, how much greater the separation of Hyrax from its neighbors is in the Procrustes plots compared to its relatively tame location in either quadratic-trend PCA.

**Figure 20.**
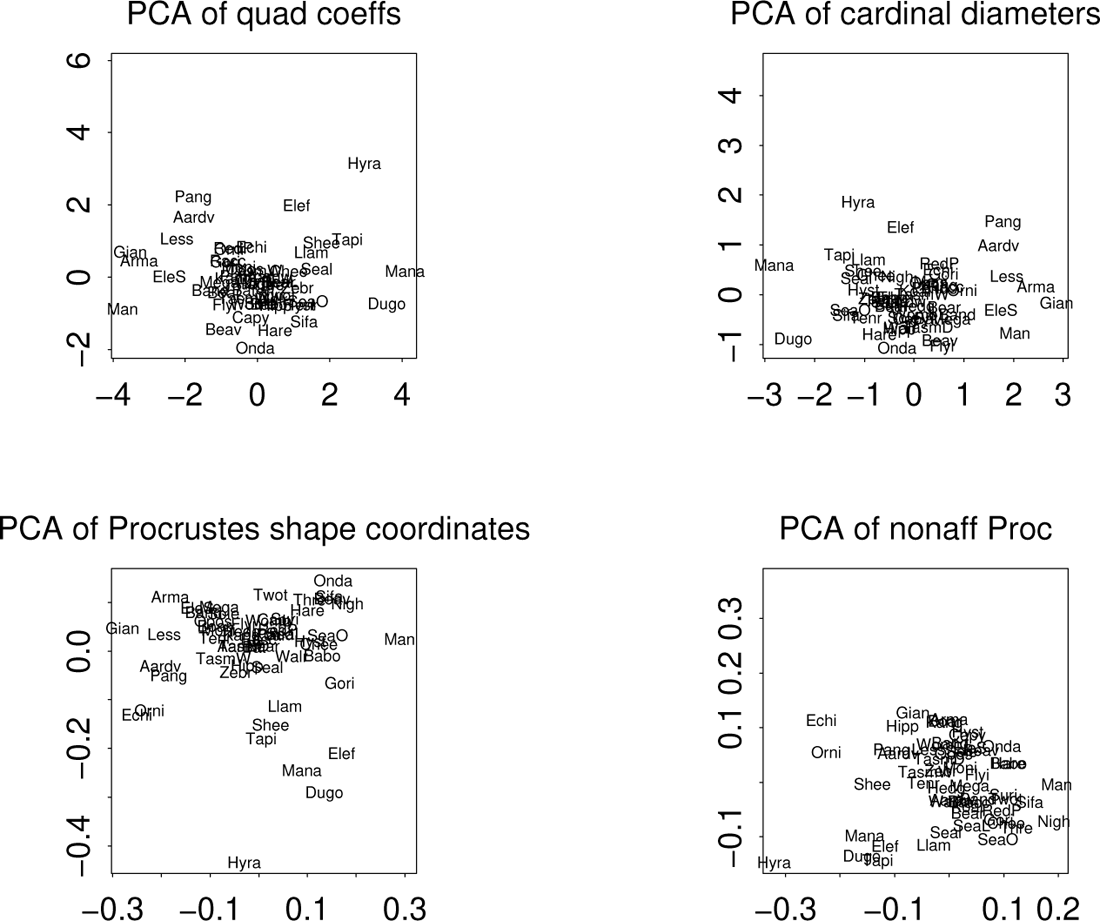
Four principal component analyses of the 55 × 13 Marcus et al. (2000) data set, 2013 version, plotted by the scatter of their first two scores. The upper left panel shows the first two principal components of the six dimensions laid out in Figure 19. The principal component analysis at upper right considers all 55 of the quartets of which 19 were displayed in Figure 18.

In conventional Procrustes-spline GMM, the results of a multivariate analysis are often diagrammed as a scatterplot amplified by marginal drawings that interpret the scatterplot’s axes as deformations. For a scatter that is not particularly Gaussian, like the ones here, the rendering by endpoints of axes is not as helpful as the more complete rendering by the full circuit of directions over the scatterplot (especially as the species anchoring this circuit around the outside of the hull are so suggestive of the overall variability of the class). Here the loci of these orienting specimens are identified by the first four or five letters of their common names as in the caption for Figure 19. These eight would outline the convex hull of the scatter were it not for the position for Elephant (“Elef”), but as explained in Marcus et al. (2000) this specimen had to be a juvenile in order to fit into their digitizing apparatus. Grids are generated by reconstructing the coefficients 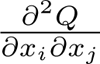, etc., of the quadratic trend from the entries for the four cardinal second derivatives in the scatterplots of Figure 14 ff.^2^ Grids opposite one another are not inverse maps but instead renderings of opposite coefficient vectors in their quadratic trend formula. Figure 21 renders these first two principal components as their effects on the 55-landmark average every 30^◦^ out of that average at an arbitrary multiple.

**Figure 21.**
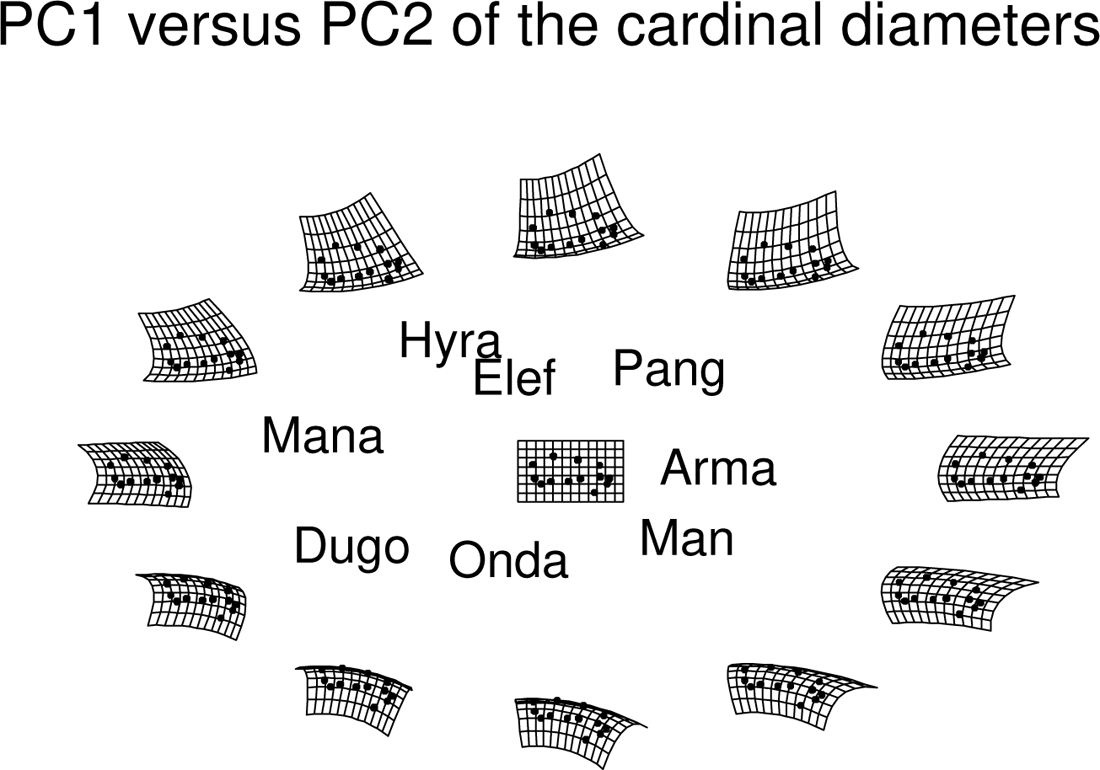
Interpretation of the axes of the upper right scatter in Figure 20 by extrapolations of the pertinent quadratic trend grids (in Cartesian format) every 30^◦^. The square grid at center bears the landmarks of the 55-specimen average from Figure 1. Affine terms are omitted.

Figure 22 represents the same twelve quadratic trend grids in the less familiar polar coordinate system we have already encountered in previous figures. What is intuited in Figure 21 as relative enlargement of the upper left (neurocranial) quadrant of the grid in the direction on which Man lies, for instance, is seen more cogently here in Figure 22 as a rotation apart of the relevant *radii* — compare the angulation of “ribs” in this region between the directions of Man and Manatee, for instance. From either of these two figures it is clear that the vertical reorientation of the face so clear in Figure 16 is not captured by this first pair of quadratic trend principal components. It may be seen instead in Figure 23, the analogous plot just for principal component 4 alone, on which Man is a striking outlier. The effect of this component is both to shorten the lower jaw and to straighten the facial angle with respect to the rest of this midsagittal configuration, two characters among the familiar synapomorphies of *Homo sapiens.* At the other end, the substantial negative scores for Beaver and its neighbors Hystrix, Capybara, and Ondatra are consistent with the large residuals at the landmarks of the Ondatra lower jaw in Figure 15.

**Figure 22.**
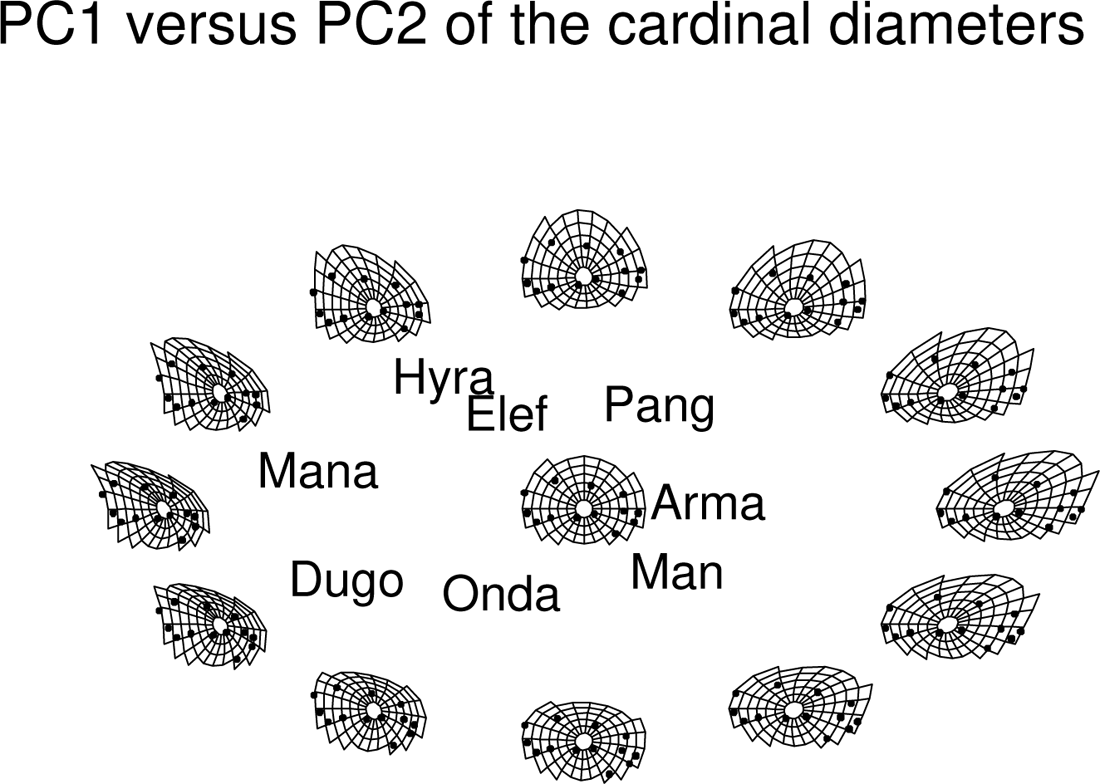
The same with the axes interpreted as trimmed polar coordinates instead, after the fashion of Figures 14, 15, 16 or those in the Supplement. Again affine terms have been omitted.

**Figure 23.**
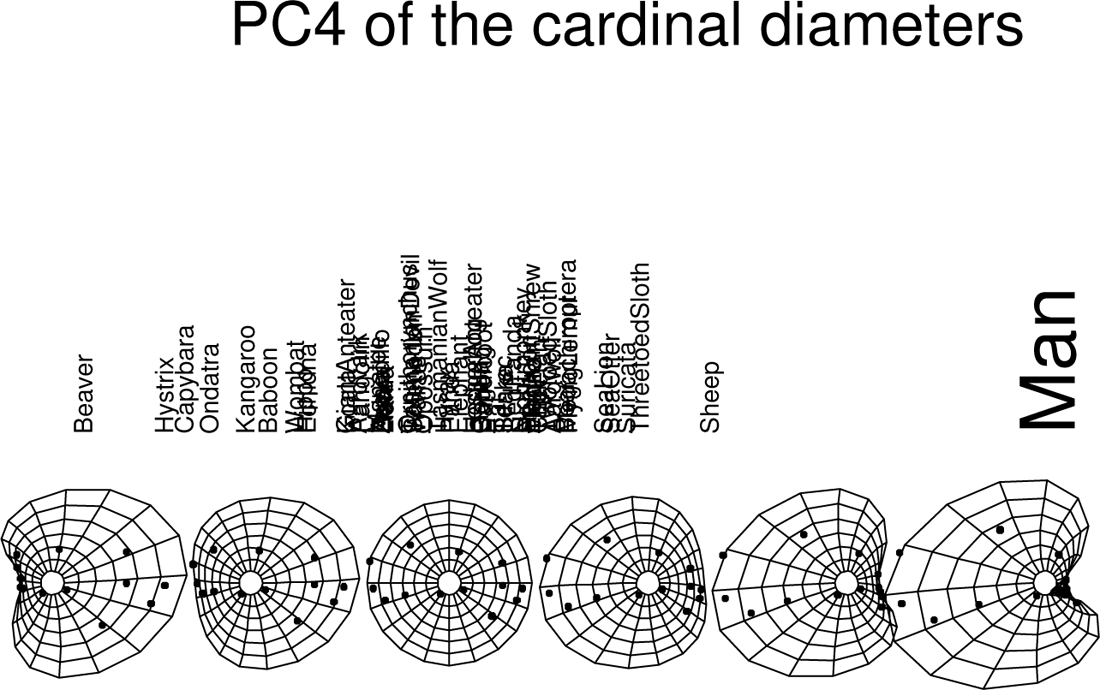
Principal component 4, ranging from Beaver to Man.

#### Ellipse axes, ellipse centers

In Figure 1 or Figures 17–18 there is strikingly variability in several aspects of these ellipses. The orientation of their cardinal diameters with respect to the Cartesian directions of the two-point registration has concerned us in connection with Figure 10 of the previous section, but there is additional information in the lengths of the ellipses’ own semiaxes. (A semiaxis is the distance from the ellipse’s center to one of the endpoints of its axes; it is half the axis length per se.) Figure 24 is an ordinary scatterplot of these two lengths for each of the 55 specimens in this data set. (As an alternative quantification one might consider the product of these two distances, which, when multiplied by *π,* is the area of the ellipse.)

**Figure 24.**
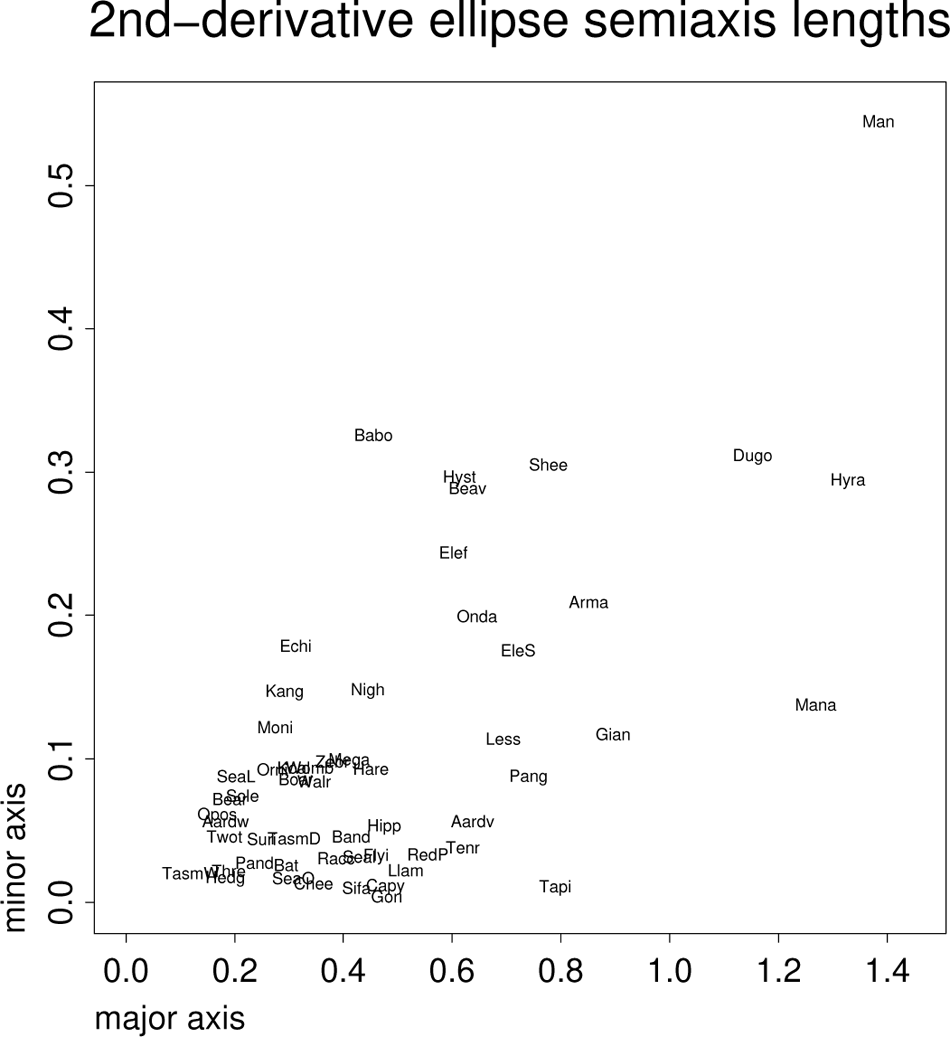
A suggestive bivariate quantification: semiaxis lengths of the second-order derivative ellipses. See text.

Some observations are clear from the figure. *Homo sapiens* has by far the largest of the minor semiaxis lengths (and also the largest ellipse area). But Hyrax, Manatee, and Dugong all have nearly the same *semimajor* axis length — the range of variation in one specific direction across the midsagittal cranium. These three examples are thus quite directional in their range of second derivatives (cross-cranium gradients of derivative). In the other direction, Baboon appears to have the most isotropic of these distributions — in the Supplement its ellipse of second derivatives is closest to a circle.

At the other extreme, the animal called Tasmanian Wolf (TasmW) here, a thylacine, has the smallest of these ellipses – it is closest to a purely linear transformation of the average configuration in terms of these quadratic trend fits. But several other species along the lower border of Figure 24 — notably Tapir, the primates Sifaka and Gorilla, and also Cheetah, Sea Otter, and Capybara — have ellipses that almost reduce to the flattened lines of Figure 8. (We saw this pattern already in Section III for the growth analysis of Vilmann’s laboratory rats, a rodent in the same family as the capybara.) The unidimensionality of the ellipses along this border suggests an actual *constraint* of their evolution — e.g., for the tapir there may be considerable developmental reorganization involved in liberating the nose for its preferred herbivore diet. In contrast, the position of Man in this scatter is consistent with our formal classification as a neotenous species, one whose heterochronies have not yet had “developmental time” to emerge. On the fourth PC axis, Figure 23, Man is most different from Beaver; in the PC plots of higher explained second-derivative variation, Figure 20, we are instead most contrasted with Tapir, Hyrax, Manatee, and Dugong. Thus the principal component analysis of Figures 20 through 22 seems quite sensitive in its numerical pattern to the pattern of semiaxis lengths in Figure 24. Geometrically this is no surprise, as the axis lengths are confounded with the separation of their endpoints, which, whenever they arise as cardinal diameters, are within the scope of linear combinations of these reference features.

Figure 25 looks more closely at the specific situation of Tapir here, the rightmost point along the lower margin of Figure 24. This particularly skinny ellipse is oriented along the horizontal in Figure 1, the long axis of the template configuration as a whole. The *y* term in its circuit of second derivatives nearly reduces to a single value (coefficients *u* = *w* and *v* = 0) and its *x* term has two equal coefficients, *s* and *t*, which is one of the special cases discussed in connection with Figure 9 (specifically, the negative of the configuration in row 2, column 1 of Figure 9). The constancy of the regression *ux*^2^ + *vxy* + *wy*^2^ for the warped *y*-coordinate implies a constant positive gradient in every direction, hence, the increasing second derivative in *x* as one passes one’s eye up the page — these grid parabolas become steadily sharper with height. The dominance of the open disk and its neighbor in the lower-right panel corresponds to a principal direction of increasing separation aligned halfway between their directions — roughly the direction at 20^◦^ counterclockwise of horizontal. This effect is *negative,* meaning that the spacing of verticals is closer toward the right side of the grid (positive *x*’s) than the left, principally in the direction aligned with the longer diagonal of the deformed rectangles projecting from the form as in the panel at lower left. In the polar plot, upper right panel, this gradient of radial spacing is particularly clear. That the roof of the calva is shortened is obvious, but that could have been accomplished in any of several geometrically different ways insofar as the midpoint of that reduced calva stayed fixed over the cranial base, shifted anteriorly, or shifted posteriorly. Here it apparently shifts mainly posteriorly, as one clearly sees in the polar grid plot at upper right.

**Figure 25.**
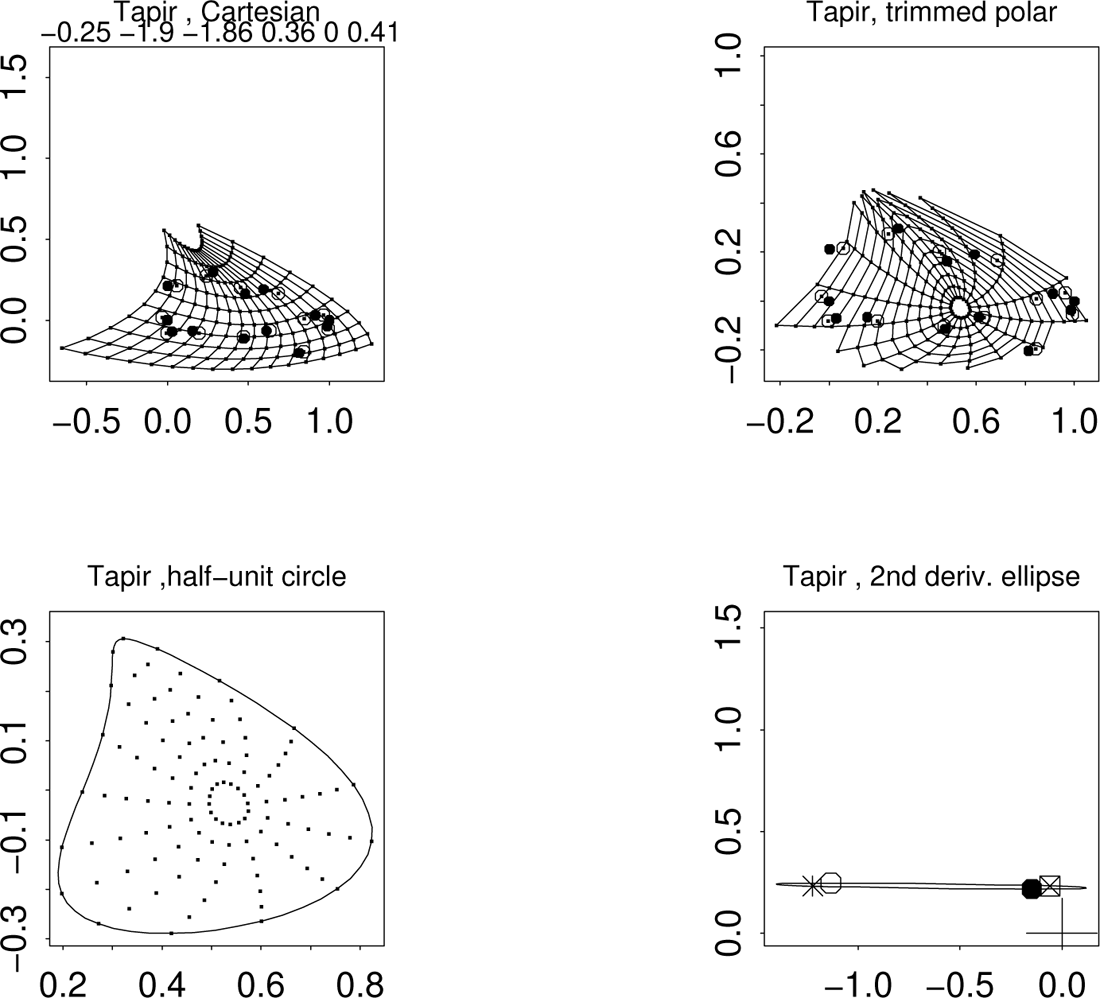
Four-panel plot akin to earlier Figures 14–16 for the tapir, the species furthest right along the lower margin of Figure 24. See text.

The simplicity of this analysis strongly suggests one search for a simple “cause” — a simple functional gradient accounting for the transformation of the template (a plausible ancestral state) with the constraints *s* = *t, u* = *w, r* = *v* = 0 of the regressions here. It affords an interesting contrast with the corresponding analysis for an entirely different animal, the manatee, Figure 26. There are strong similarities here — juxtaposition of the 45^◦^ and 90^◦^ pairs, horizontality of the ellipse in the coordinate system of Figure 1. Notwithstanding that the rank-order of the six regression coefficients *r* through *w* is different, the overall impression of this quadratic trend is remarkably similar, though more intense in the sea creature. That “teardrop” shape of the half-unit circle plot, lower left panel in both of these figures, may well be a novel character parameterizing the feasibility of extreme forms across a wide subset of the mammalian radiation. (Think of the teardrop as the combination of suppressed angular spacing along with enhanced radial spacing in the same polar sector — concentration of an intensified second derivative of a trend coefficient in one relatively narrow direction together with a suppression in the perpendicular direction.)

**Figure 26.**
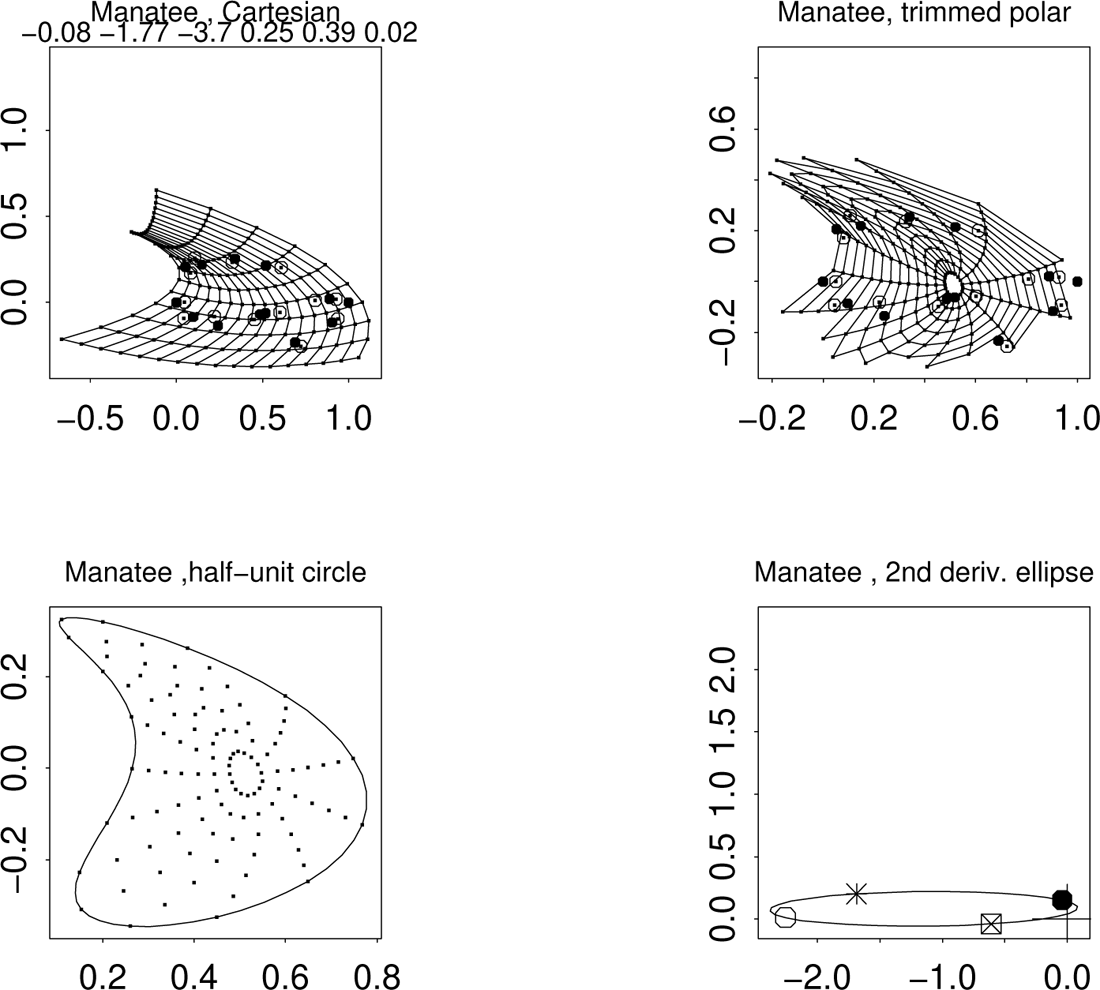
The same for the manatee. Note the similarity to Figure 25 in spite of the very different ecology of the creatures.

Either Tapir or Manatee, similarly extreme according to the quadratic-trend feature space, can be profitably compared to a contrasting situation, a rotation at 45^◦^ of those cardinal diameters. In the upper left panel of Figure 20 the PC1–PC2 score for the elephant (“Elef”) is relatively isolated along the vertical (PC2) axis, whereas the point for Manatee is where one expects it (far out along the PC1 axis). This divergence of positions corresponds to the appearance of the polar-coordinate panels at upper right in Figure 25 versus Figure 27 below. For the manatee, there is a sharp convergence of polar radii along a northwesterly direction; for the elephant, along a northerly direction instead. The elephant’s half-unit circle plot, lower left in Figure 27, shows a version of the teardrop pointed vertically rather than to the northwest, corresponding to the rotation of its cardinal diameters around the ellipse at lower right. (There has also been an elevation of this ellipse above the horizontal axis of its panel, corresponding to the positive second derivative of the spacing of the horizontal curves in the upper left panel.) In keeping with Sneath’s vision of factors it would be reasonable to search embryologically for some putatively one-dimensional form-factor that accounts fairly simply for this shape change, in spite of its extremely large magnitude, as a plausible two-parameter change of the quadratic trend coefficients. Teardrops may indeed be one good candidate for the rhetoric of form-factors that Sneath hoped for in his 1967 article. I return to this possibility in the Discussion.

**Figure 27.**
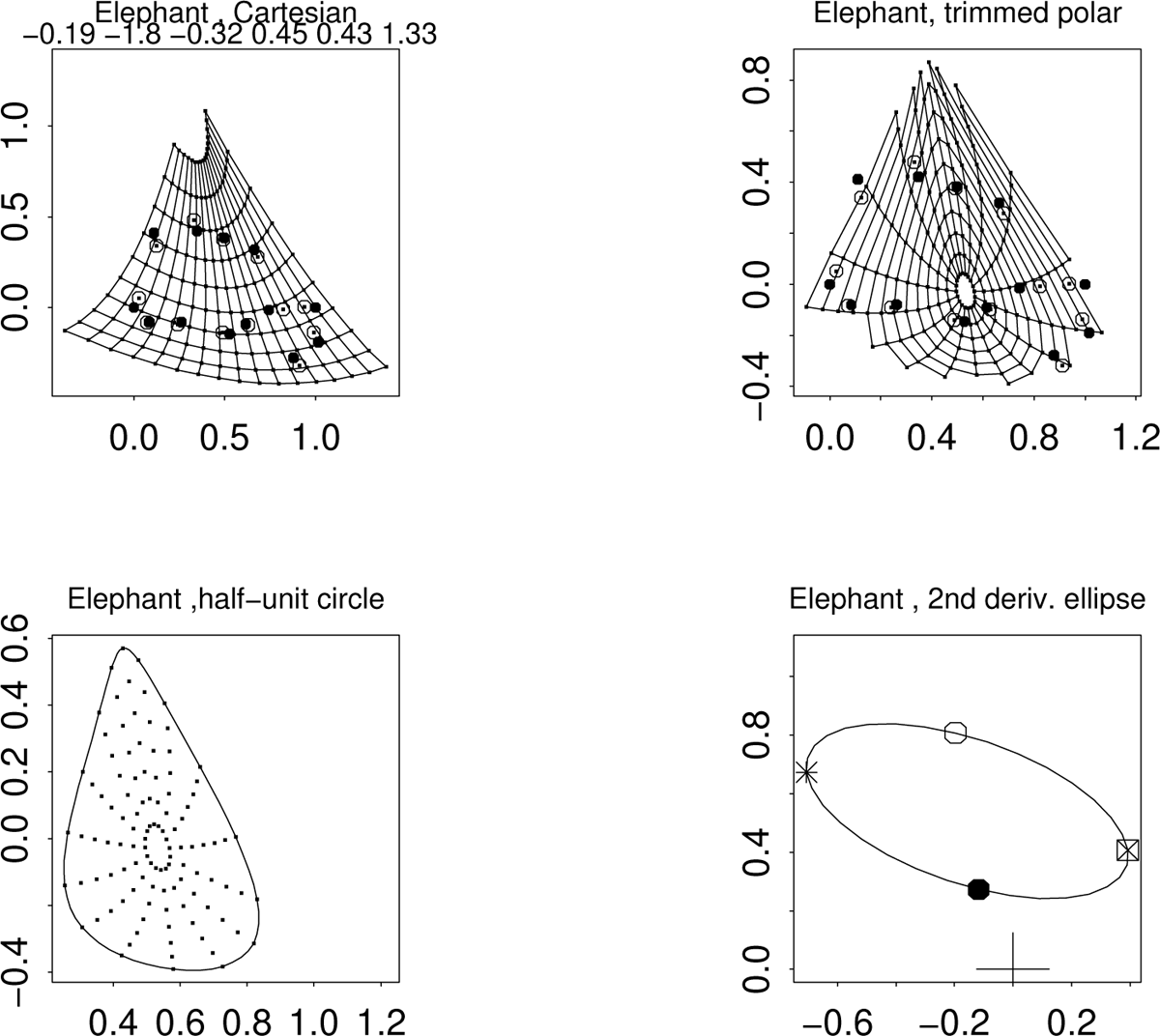
The same for the elephant.

It is fair to inquire about the uncertainty of this lengthy series of analyses against that initial analytic decision, the choice of a longitudinal baseline. Figure 28 complements the right panel of Figure 1 by an alternative for this highly diverse sample. It is a source of considerable comfort that the obvious features of Figure 1 are replicated here to a great extent in spite of the quite different functional contexts of the baseline points chosen. (This alternate baseline makes an angle of 33.5^◦^ with the nearer of the axes of the usual baseline; the maximum possible such deviation is 45^◦^.) In particular, the large ellipses are still large in general, and the general arrangement of these large shapes is similar on the page up to a linear transformation. The canonical correlations of this pair of ordinations are 0.909 and 0.716; the *R*^2^ of predicting the ellipse cardinal points on the right from their coordinates on the left is 0.852. The first two canonical correlations of the first four sets of PC’s are 0.983 and 0.980.

**Figure 28.**
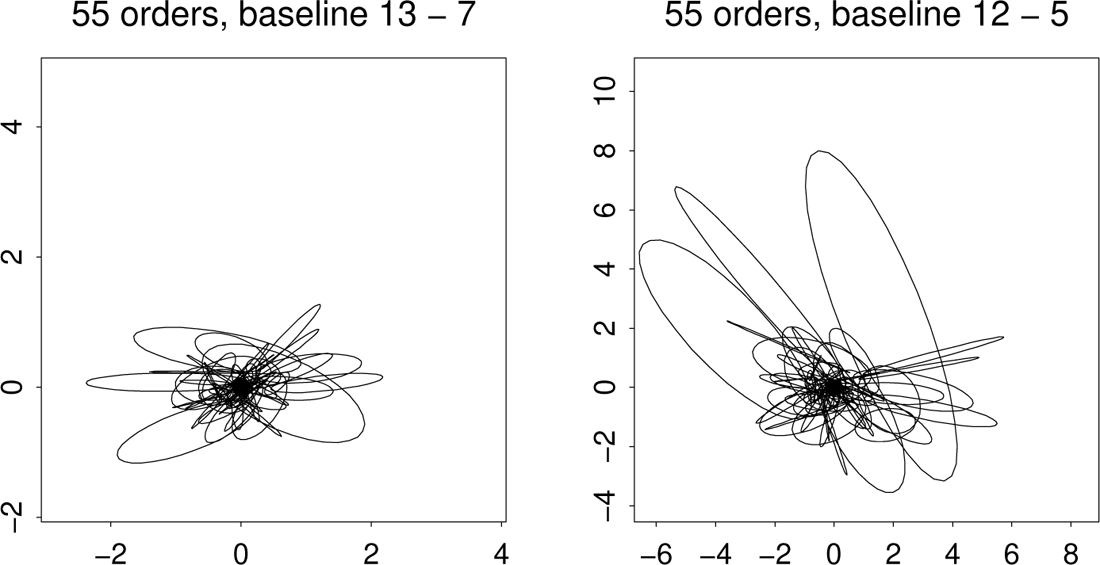
Critique of Figure 1 (right) by change of baseline to 12–5 in the numbering of Figure 1 (left), anterior foramen magnum to fronto-nasal sagittal, which makes an angle of 33.5^◦^ with the baseline of the other figures in this section.

Complementary to the display of the intraspecimen range of second derivatives in Figure 24, the axis lengths of their ellipses, is the information content of that ellipse’s center. Analytically this is the same as a morphometric quantification introduced decades ago, the concept of *roughness* explained in Bookstein (1978). The roughness of a grid transformation at a grid point is defined as the second-order approximation of the discrepancy (corrected for cell size) between the deformation of the centroid of a grid cell and the centroid of the deformation of its vertices. According to the formula on page 109 of the 1978 reference, this discrepancy is the vector whose *x*-coordinate is the sum of the two unmixed second partial derivatives of the *x*-coordinate of the deformation, while the *y*-discrepancy is the sum of the same two second derivatives of the *y*-coordinate. The mathematician would notate these components as the Laplacians (Δ*x*^′^, Δ*y*^′^) where each Δ is the sum of the two unmixed second partial derivatives of the deformed coordinates separately: the expansion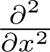^3^. So this Laplacian vector equals the sum of the vectors that are plotted with the filled disks and the open disks in Figure 20, and that sum, in turn, is twice the average of those two disk locations, which is to say, the center of the ellipse. These are plotted in Figure 29 (left) for each of our 55 species. (As a corollary, we uncover another characterization of the quadratic trend representation: grids having a^3^

**Figure 29.**
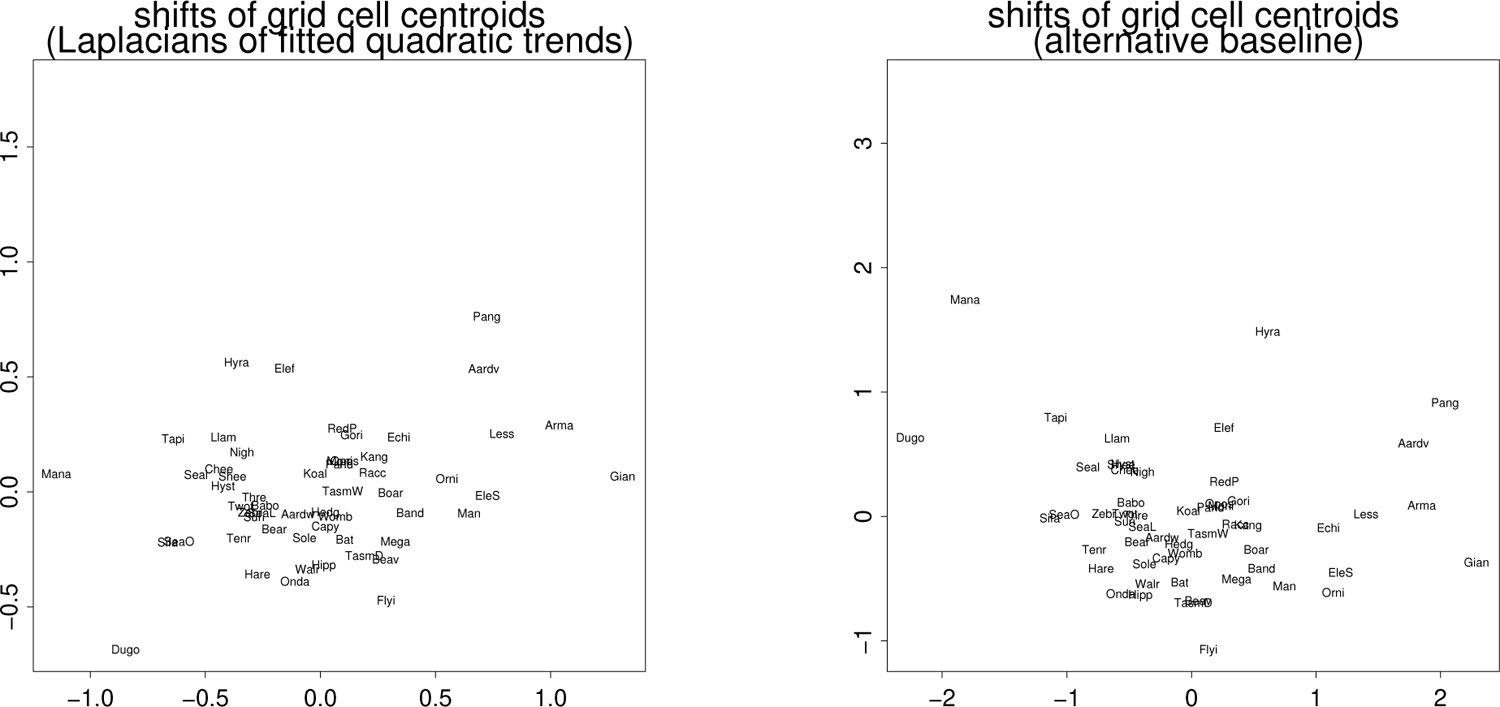
Complement to Figure 24: scatterplot of the centers of the ellipses in Figure 1 or Figure 28. The location of each point is the vector of separation between the quadratic prediction of the centroid of a grid square and the centroid of the prediction of the corners of that square. Labels are the short names of the 55 species as in Figures 19, 20, or 24. (left) For the baseline of Figure 1. (right) For the alternate baseline at right in Figure 28. The extreme taxa are the same.

This ordination is intriguing. Near (0, 0) we find the names of the species for which the deformation fitted by the quadratic comparison to the average is most nearly linear: Koala, Hedgehog, Tasmanian Wolf, Wombat, Aardwolf, Panda, Capybara, Monito. Around the outline of the distribution are the species for which the center of the ellipse is farthest from (0, 0): Armadillo, Lesser Anteater, Elephant Shrew. and (most extreme in either baseline direction) Giant Anteater versus Manatee. All these extremes represent shifts along the *x*-axis, the baseline from Figure 1, which is so much longer than the perpendicular skull height dimension. The species that are most shifted in the orthogonal direction are fewer — Hyrax, Flying Lemur, and Elephant – while Dugong, Pangolin, and Aardvark show substantial roughness in both directions. Any of these extremes might qualify as a morphogenetic innovation or synapomorphy in analyses of how these skull forms relate to their phylogeny. This interpretation is stable against the change of baseline in Figure 28 — the alternative scatter of ellipse centers, Figure 29 (right), shows the same list of outliers. In neither scatter is the position of the center of the ellipse for Man particularly unusual, not even for a primate (see also Bookstein, 2018, Figure 3.20c). It is not the average of these second derivatives over the circle of directions but their directionality per se that is unusual for our species among the mammals (Figure 24).

#### Retrieving the linear component

Be assured that the second-derivative ellipses of this new analysis are not confounded with the first derivative term that the exposition has hitherto ignored. The computation via second differences (Figure 4) renders that linear term algebraically irrelevant except for scaling for baseline length. Nevertheless it is worth exploring the posssibility of correlations between one or another parameterization of these ellipses and the four parameters of that previously overlooked linear term. A survey of a variety of these approaches yields an interesting pattern of correlations with ellipse form of two aspects of those linear terms, the regression of the specimen *x*-coordinate on the template *x*-coordinate (a coefficient that ranges from 0.300 for Hyrax to 0.407 for Boar) and the analogue in the *y*-dimension (which ranges from 0.128 for Giant Anteater to 0.667 for Man). From a range of possible diagram styles I have selected the version in Figure 30, which duplicates the loci of Figure 28 (left) but modifies the line type in two different ways: at left, by comparison to the median of that regression coefficient for *x*; at right, for *y*. In the left panel, the ellipses drawn in line correspond to the species for which the predicted *x*-coordinates of the configuration fall generally to the left of their position in the template (the average), and the opposite for the ellipses drawn as point series; in the right panel, it is the ellipses for species with predicted *y*-coordinates lying above the averages that are drawn in line. It appears that these ellipses do indeed cluster to some extent by characteristics of the associated linear component. (The correlation between the two linear coefficients, x.x and y.y, is a full −0.7, and the correlation between y.y and the northeasterly second derivative in the *x*-direction is nearly as large at −0.63.) Such associations are likely to serve as one productive context for further morphometric surveys of *Mammalia*.

**Figure 30.**
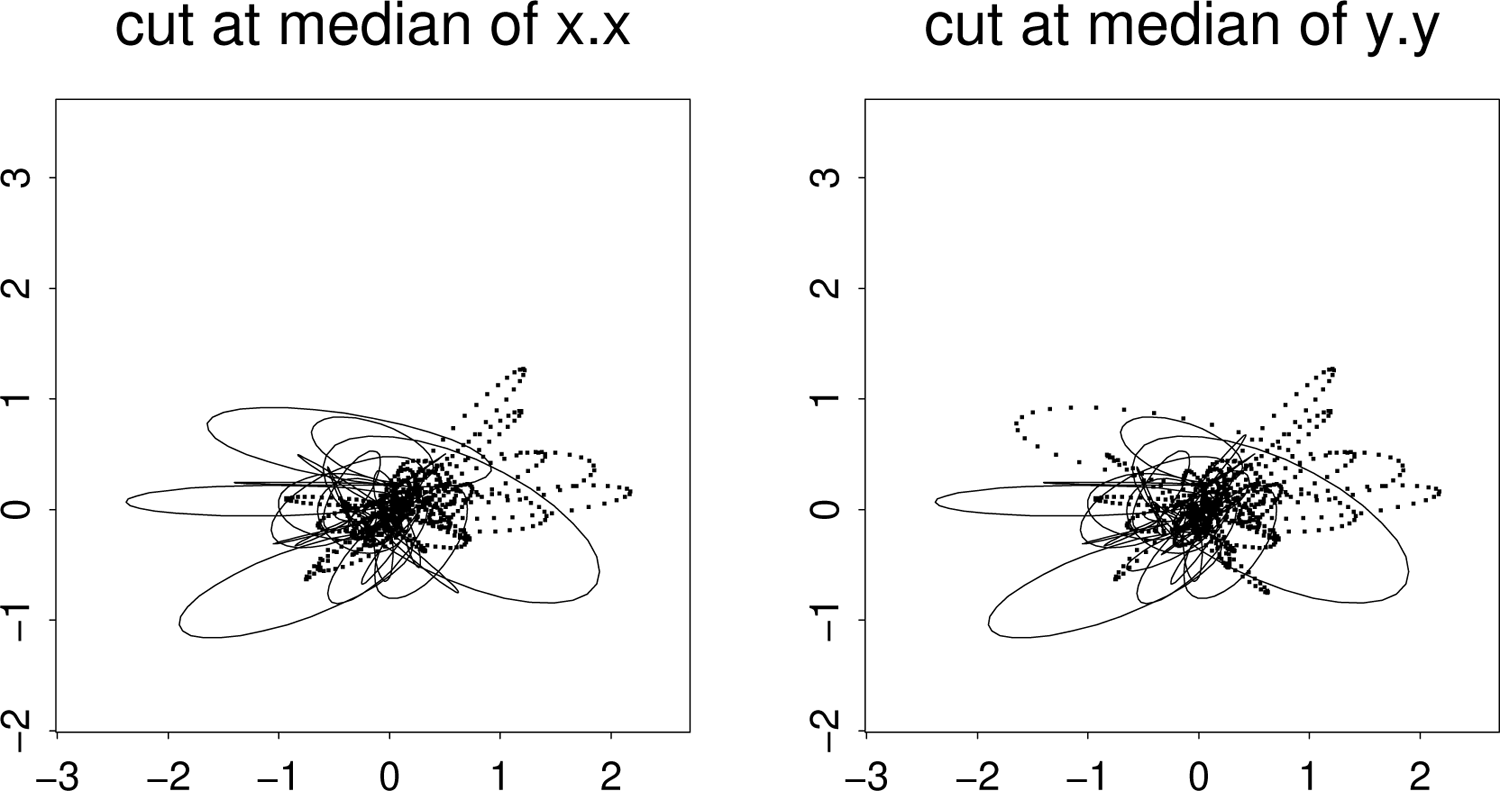
Subsetting the 55 2nd-derivative ellipses by parameters of the first derivative. (left) Ellipses for species with the regression coefficient for landmark *x*-coordinate on template *x*-coordinate less than than the median drawn in line, others as point strings. (right) Lines for regression coefficient of configuration *y* on template *y* greater than the median, points for species with coefficients below. The panels are similar overall.

The analysis here of the 55-mammal data set is quite different from that published by Marcus and colleagues and different also from the analysis in my own textbook of 2018. The original publication (Marcus et al., 2000) emphasized mainly the matrix of Procrustes distances among specimens along with the implications of those distances for a cladogram or a phylogeny. I have severely critiqued Procrustes distance in a range of papers over the last decade (see Bookstein 2015, 2016, 2018) and need not repeat the critique here except for a summary: Procrustes distance is not a meaningful biological quantity, as it depends too much on the investigator’s subjective choice of landmarks and, by treating all landmarks with complete algebraic symmetry, offers no access to prior anatomical or functional insights that might direct interpretation of multivariate analyses of shape coordinates. For example, the ordination in Figure 24 of the directionality of a quadratic trend, a quantity that likely relates to evolvability in more general senses, has no equivalent in the standard morphometric toolkit.

In general, quadratic regressions may be expected to organize some potentially important aspects of landmark configuration data more effectively than either the Procrustes maneuver or the various thin-plate spline visualizations of the resulting shape coordinates whenever the biological phenomenon under exploration (in Section III, the pattern of rodent neurocranial growth; in Section IV, the huge diversity of geometric arrangements of the adaptively radiated mammalian cranium) involves large-scale effects on these configurations. The scatterplot at lower right in Figure 13 assures us that most of the variation over this class in this 13-landmark configuration is captured by the trends here that, beyond any linear terms, support the simple parameterization displayed in Figure 1. The ellipses that organize these circuits of directional derivatives suggest a space of just six degrees of freedom better for biological interpretation (i.e. more anatomically organized) than either the thin-plate spline or the multivariate analysis of the whole set of 22 or 26 shape coordinates can manage. Neither the circuits of directional second derivatives nor the major axes of the ellipses that trace them are accessible from the conventional Procrustes–thin plate GMM toolkit. In the presence of diversity as great as what is laid out in Figure 1, we must protect ourselves from the utter arbitrariness of the original data resource here, the finite set of disarticulated landmark locations, which were identified more for the convenience of the graduate student doing the digitizing than for any promise of cogency for the resulting evo-devo hypotheses. The quadratic regressions draw the original landmark data, whatever their count (as long as it is greater than 6), into a summary designed to suggest explanations of a morphodynamic or biomechanical sort, explanations plausibly associated with the embryology or ecology of these highly diverse genera.

In this way the quadratic trends rescue D’Arcy Thompson from the sheer obscurity of the method (if any) by which his line drawings were generated, while at the same time their distillation into those six-parameter ellipses of second derivatives rescues Peter Sneath from his unfortunate preoccupation with distances and sums of squares. Together these formalisms, one old and one new, may help to deprecate GMM’s current focus upon Procrustes shape coordinates and thin-plate splines, both of which seemed much more promising back in 1993 than they seem today. Manatee and Tapir are satisfactorily close in the principal component scatterplots of the regression coefficients and the cardinal diameters, Figure 20. But adjacency in principal component plots is much less informative than similarity in a-priori geometric subspaces dealing explicitly with a-priori patterns of landmark rearrangement, the spaces where patterns lead to morphodynamic or functional explanations. Then closeness in principal component projections is much less informative than closeness in explicit character spaces defined a-priori — in the comparisons of Figures 25 through 27 this is that “teardrop shape” of the half-circle plots at lower left.

We have seen how the parameters of this paper’s ellipses seem meaningful beyond their algebraic identity as linear combinations of shape coordinates. Three different presentations of the complete space of these shape representations have been introduced. One iconography, Figure 19, is the full set of six regression coeffients in the approriate Cartesian setting of three complex numbers. Another representation, Figure 17, shows the ellipses in full relevant detail for a perspicuous subsample of 19 interesting forms, while a third view, combining Figure 18 (the scatter of just the cardinal points) with Figures 24 and 29 (the geometric parameters of the ellipses per se), parameterizes these ellipses in geometric rather than algebraic terms. Each of these involves just six large-scale parameters regardless of the count 2*k* − 4 of dimensions of the actual shape space, and each interpretation is intrinsically more cogent than anything offered in the conventional GMM toolkit. (Recall how Section II guided us in interpretations of the patterns of the largest three regression coefficients out of the sextet.) Restricting shape space to this six-dimensional subspace, in other words, achieves D’Arcy Thompson’s purpose — a simpler version of shape comparison regardless of how complicated the template may be — while also serving as a potentially feasible character space such as Peter Sneath was hoping for: geometric patterns that could be exogenously tested for meaning phylogenetically, morphodynamically or functionally.

#### A potential character: the “teardrop”

Toward the end of Section IV I suggested that the emergence of a teardrop in the half-unit polar plot of second directional derivatives might actually be such a potential character. That Sneath’s hope for a feature space was mostly ignored during the entire late-20th-century development of the current “morphometric synthesis” was an unfortunate oversight on the part of all its developers, including me. Figure 31 hints at a potential phylogenetic use of this quadratic trend approach in Sneath’s spirit. At left I have compiled all 55 of the half-unit outline curves from all the four-panel figures like the exemplars scattered throughout this essay. (All 55 of the curves may be reviewed in the Supplement.) There is a central skein of curves that differ little from circles, plus a variety of apparent deviations showing sharper curvature. It is convenient to parameterize these deviations by one classic measure of curvature, the change in direction between each of the line-elements of those deformed circles and their neighboring element, divided by the length of the chord that spans the pair. When we characterize our 55 species by the maximum of this improvised index, the top fifteen selections are an interesting list of species. The right panel of Figure 31 draws these 15 deformed circles by themselves, now after a centering just to simplify the graphic, and labels each with its short species name at the vertex of maximum curvature using the index just explained.

**Figure 31.**
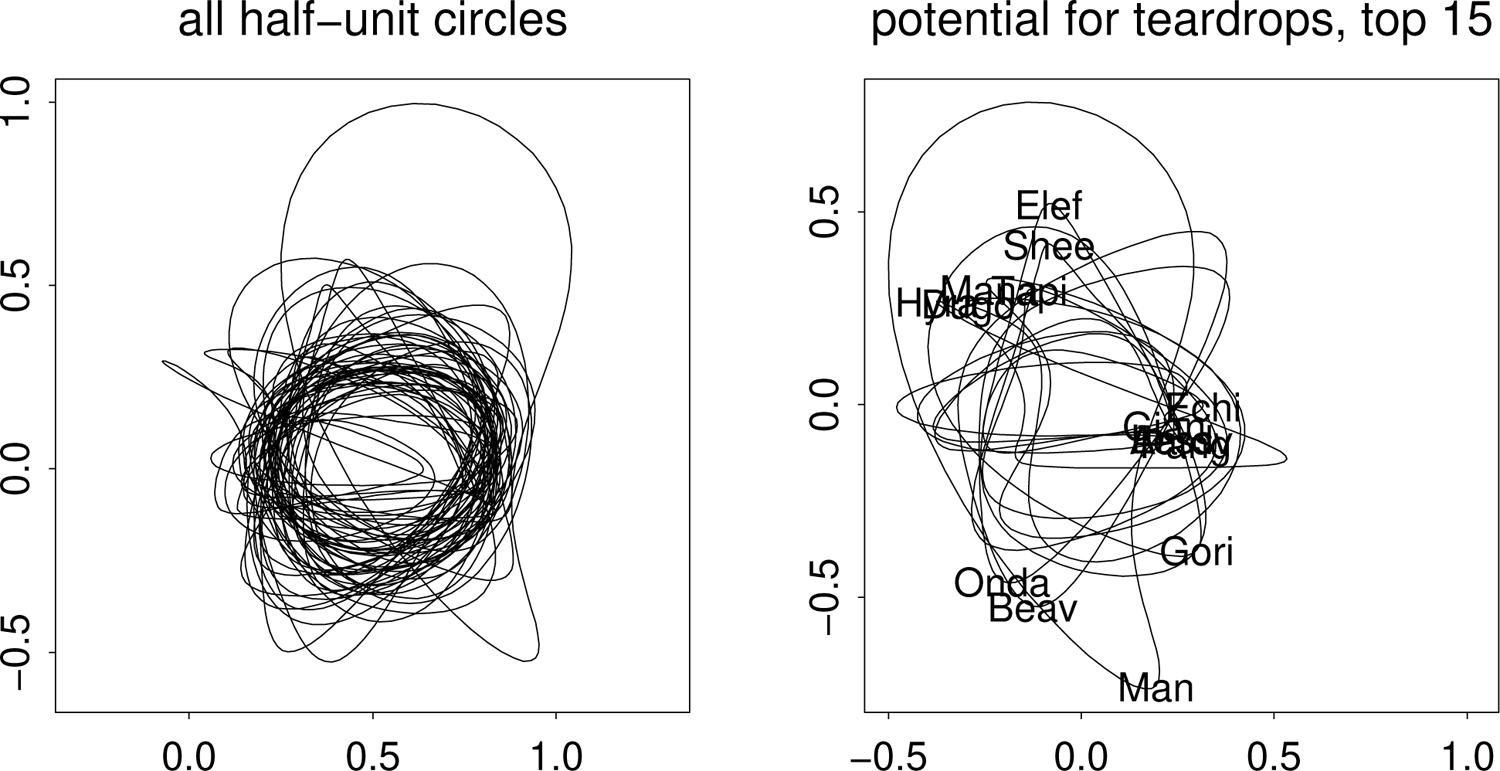
Analysis of the polar plots of 55 specimen-specific quadratic trend regressions. (left) Overlay of all 55 instances, collected from the Supplement. (right) The fifteen out of the 55 having the highest peak curvature scores as explained in the text. Each short species name is printed at the point of peak curvature. The illegible name list at upper left is Hyra(x), Tapi(r), Mana(tee), Dugo(ng); that at center right is Echi(dna), Gian(t Anteater), Less(er Anteater), Pang(olin), Aard(vark). Elef: Elephant. Shee: Sheep. Gori: Gorilla.

The fifteen vertices can be reviewed in five groups that are quite suggestive of biological interpretations, as follows. Toward the top of the diagram are two forms, Sheep and Elephant, which peak in a direction perpendicular to the long axis of Figure 1. Counter-clockwise from them is a cluster of four forms of which two, Manatee and Tapir, aroused our special interest in Section IV in connection with the zeroing of one of their ellipses’ semiaxes. A third, Dugong, is ecologically similar to Manatee, while Hyrax (associated with the most deformed polar circle out of all 55 of Marcus’s exemplars) is phylogenetically related to Elephant in the preceding cluster. Continuing counterclockwise, Ondatra (muskrat) and Beaver clearly overlap in their ecology. Our circuit next encounters a pair of singletons, Gorilla and Man; in view of their phylogenetic proximity, the separation might be attributed to *H. sapiens’s* neoteny. A final cluster of five species (Pangolin, Echidna, Aardvark, Lesser Anteater, Giant Anteater) collects some the forms with snout very highly compressed vertically (a cluster that would incorporate four more forms of the next five highest curvatures, including Elephant Shrew, Armadillo, Tenrec, and Bandicoot, not shown). Taken as a whole, the clustering here seems quite distant from a random pattern, but ought to be considered to represent some combination of niche and evolvability over the whole class of mammals. In my opinion, some less improvised version of this clustering may well justify deeper explorations.

Note how very nonlinear a computation this was. For each of the 55 specimens we located the maximum around the polar circuit of a one-dimensional summary of a two-dimensional regression prediction spanning the full configuration of landmarks but parameterized as a function of direction, not position; and inasmuch as what we are taking the maximum of is already a second derivative (for that is what our index of curvature is approximating), the result per se must be understood as having located the zero of a *third* derivative. In comparison with the hidden sophistication of this tactic, details of the Procrustes versus the two-point registration (a choice mandated in order that there be shape coordinates to be regressed) are trivial. The quadratic regression itself, equation (3), is not linear in the coordinates of either the target specimen or the template, owing to the presence of the terms in *x*^2^*, y*^2^, and *xy*; the polar graphic is linear in the regression coefficients but nonlinear in its argument (*x, y*) = ½ (cos *θ,* sin *θ*), where the constraint locking the radius at 1/2 happens to be half the distance between the landmarks of that arbitrarily chosen baseline.

The cardinal diameters are linear in the quadratic regression coefficients, and so their principal components, Figures 20 through 23, are likewise accessible from the data set of these vectors, though the necessary matrix notations would be clumsy. But the extraction of the principal axes of the second-derivative ellipses, Figure 24, was already beyond the capabilities of that standard toolkit, as the ellipses in question are not based on data-driven covariances, and the detection of the clusters in Figure 31 is clearly beyond any standard GMM software system — it is, indeed, a task for the natural intelligence of the investigating biologist, one who is aware of the currently accepted phylogeny of these same specimens. Clearly linear multivariate analysis alone is not adequate preparation for the biometrics of landmark configurations. Of course there are many other branches of geometry applicable to biometrics as well — tensor calculus, differential geometry of surfaces, catastrophe theory, projective metrics, fractal geometry, hyperbolic geometry, to name just a few. It is no reason to privilege the linear vector geometry of covariance matrices among this multitude of toolboxes that we have learned how to teach it.

#### Trends versus interpolating splines: the contrasting roles of landmarks

Turn now from discussion of the dimensions of this simplified shape space to a more detailed examination of the grid figures themselves, the rendering of the picture area in-between the landmarks of the data set. Equation (3) of Section III declares that the transformation grid we want is the minimizer of a sum of squares over what is represented as a prediction error at each landmark in turn: the displacement of the predicted location of a landmark from its location as actually observed in some specimen. That equation writes the numerical task here as the computation of numbers *a, b, c, d, e, f, r, s, t, u, v, w* that together minimize

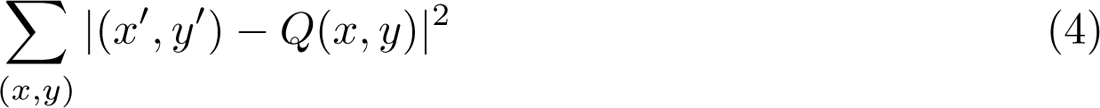

where *Q* is the quadratic trend function from equation (3), (*x, y*) and (*x*^′^*, y*^′^) are the landmark locations of template and target, respectively, and as before the vertical bars stand for ordinary distance on the picture plane. Six of the resulting values, the coefficients *r, s, t, u, v, w,* of *Q*’s quadratic terms, are intended as characters of the specimens, and the sum is over the list of the landmark locations themselves.

Remembering that the values *r* through *w* are half the actual second derivatives of the fitted mapping *Q*, it is instructive to juxtapose the assignment of minimizing expression (4) to the task that is solved by the conventional thin-plate spline as it has been exploited since the early 1990’s all over the GMM community. This is the task of choosing the function *Q out of the restricted family of possible Q’s constrained a-priori to exactly match the values at the landmarks — Q*(*x, y*) = (*x*^′^*, y*^′^) *at each pair of locations —* that minimizes the somewhat different-looking expression

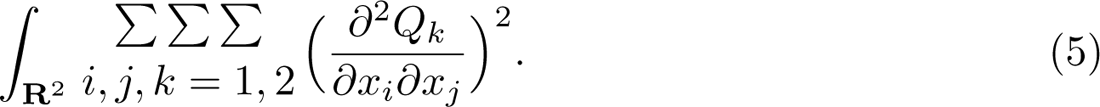

The symbol **R**^2^ here stands for the whole Cartesian plane, out to infinity, and the subscripts *i, j, k* range only from 1 to 2. The *i* and the *j* stand for the two Cartesian dimensions of the template, which can be the same (*x* and *x*, or *y* and *y*) or different (*x* and *y* or the opposite, which give the same 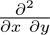, as it happens), and *k* is 1 or 2 for the Cartesian coordinates of the target.

What is being minimized in equation (5) isn’t the error of fit of the function *Q* at the landmarks — that error is identically zero as the essential restriction on *Q* in the statement of this other problem — but is instead a property of those second derivatives in equation (5) that were taken to be constant coefficients in the formula for *Q* of expression (4): their *own* sum of squares, integrated (added up) over the whole picture, should be a minimum. This is no longer any sort of error at the landmarks — it is, rather, a quite different kind of error that would arise if we compared the function *Q* to one that had all of its second partial derivatives zero, which is to say, an exactly linear map. Formula (4) cares about the landmark locations; formula (5), about the grid in-between and all the way to the edges of the picture and beyond. The formulas that give us this second kind of optimizing *Q* are very clever indeed — mathematicians derived them only in the 1960’s — but the values of their coefficients (the vectors *L*^−1^*H* of Bookstein 1989 and other presentations) are of no empirical interest at all, only the integral of summed squared second partials that engineers quickly recognized as the bending energy of an idealized thin plate. And in formula (5) the locations (*x*^′^*, y*^′^) of the target landmarks are not of any explicit interest, either: only that the function *Q*(*x, y*) exactly reproduces each of them.

Then several formal relationships between equations (4) and (5) are noteworthy. Either optimum is *rotatable* — when one of the Cartesian coordinate systems is rotated, both the minimizing spline (5) and the quadratic regression (4) rotate directly with (*x*^′^*, y*^′^) and inversely with (*x, y*). And both approaches can be proven to supply unique global minimizers except under singular circumstances (e.g., all landmarks in a line). But the differences are more numerous, and more salient. Most importantly, there is no “error” *at* landmarks in (5), the quantity whose sum of squares is minimized in (4). Instead, what is minimized in (5) is the integral of the variation from point to point of what could be regarded as a model for variation of the regression coefficients *b, c, e, f* over position — the squared partial derivatives of the *linear* part of the fitting function *Q*, which constitute the second partial derivatives that the definition of the function *Q* in formula (3) sets to be constant everywhere. And while equation (4) involves a sum over the landmarks alone, expression (5) integrates over not just the interior of the landmark configuration but all the way out to infinity (which is why these maps *must* become linear toward infinity — *r, s, t, u, v, w* all have to drop to zero pretty fast for their integrals to be finite at all, let alone minimized).

Sums of squares of position discrepancies and integrals of sums of squares of second partial derivatives are not equivalent either numerically or conceptually. The coefficients of *Q* that serve as characters in the quadratic trend approach are mere nuisance variables in the spline approach, never examined or subjected to multivariate analysis until they are summed after each is multiplied by the very peculiar formula *r*^2^ log *r* where *r* is the same Pythagorean distance as in formula (4), but now applied to an entirely different argument, driven not by the configuration of landmarks but instead by the continuum of gridded points themselves as they relate to the various data points of the template one by one. But in the quadratic-trend graphics, the error of fit landmark by landmark is explicit in the relation between the filled disks and the open disks in contexts like Figure 14, while the six second derivatives of *Q*, hinted at in the variation of the shapes of the little grid squares across the diagram, become explicit when the polar coordinate construction is assessed by the second-difference method of Figure 4.

In short, the two approaches to landmark configuration analysis, polynomial trend fits versus thin-plate splines, suit wholly different explanatory styles and bioscientific purposes. The coefficients that the trend method uses to export biometric meaning are discarded immediately after computation by the spline. The second partial derivatives of *Q*, the relevant constants of the quadratic trend fit, are the minimand of the spline method, which refers to the embedding space of the landmarks in a manner to which the regression equation has no access. And the error of fit of the trend method is forced to exactly zero in the spline method. This paper argues that, in view of the arbitrariness of the landmark lists driving GMM data sets from the outset, one of these assignments is much more conducive to subsequent biological insight than the other is.

The displays in Figure 13 already permitted us to quantify the net modeling power of this paper’s quadratic trend analyses vis-à-vis an analogous attempt at large-scale modeling, the *partial warps* of the thin-plate approach (Bookstein, 1989, 2014, 2018). The upper right panel, for the spline-based approach, reduces the landmarkwise variances of the upper left panel by a factor of 0.65; the equivalent comparison along the lower row, for this paper’s quadratic approach, reduces the summed variances by a factor of 0.85. The analyses in the upper row allow the coefficients of the prediction function to vary but require them all to attenuate sharply toward zero away from the origin; those in the lower row leave those coefficients constant, instead optimizing the predictive accuracy of that single shared quadratic regression. Clearly the constant-coefficient approach inherited from Sneath (1967) dominates the varying-coefficient approach of the thin-plate consensus in this context of predictive accuracy per se. The discrepancy between the approaches is understandable — the regression approach is, after all, a least-squares optimization already, one that would become exact after 14 more terms are added, but these additional terms involve higher powers of *x* and/or *y* that are correlated with the quadratic terms already present and so are unlikely to add much accuracy overall; whereas the first three components of the principal warp decomposition are suboptimal for any decomposition except bending energy, but that is not a sum of squares of anything referring to the actual morphology of the organism, bounded as it is in extent.

#### A different thin-plate spline

The quadratic trends analyzed and diagrammed so far in this paper are regression fits that leave unexplained variation at each landmark. In the other approach to combinations of squares, the thin-plate spline formalism, the map is an interpolation rather than a regression, and the sum of squares that is being minimized is not a net prediction error but rather a sort of complexity, the version the literature calls “bending energy.” For the conventional thin-plate spline, bending energy is one quantification (albeit perhaps a peculiar one) of the departure of its *Q* from a linear map. But ever since the initial promulgation of these splines there has existed an alternative, the *quadratic thin-plate spline,* that likewise fits each landmark exactly, by analogy with our conventional thinplate spline, but for which the drift term (the polynomial part that is not a function of the distance of a grid point from every landmark location in turn) is a quadratic map rather than a linear one. This is the thin-plate spline that minimizes a different kind of “bending” energy, namely,

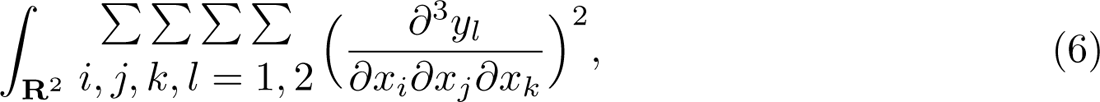

sum of the integrals of the squared *third* derivatives of the map (*y*_1_*, y*_2_) of *x*_1_ and *x*_2_. And the additional contribution of each landmark, to be multiplied by some complex number, takes the form *r*^4^ log *r* instead of *r*^2^ log *r.* For the theory of this spline, which is closely analogous to the usual linear-trend spline of the conventional GMM toolkit, see Kent and Mardia, 2022. I have exploited it once before, in Bookstein 2004, where, however, it was applied to a comparison of forms that differed hardly at all.

At upper right in Figure 32 is the Vilmann comparison as in Section III but using this exact fit with quadratic drift (not “trend”) curve analogous to the quadratic fit with regression errors that was displayed in Figure 10. Clearly it is failing to cope with the situation along the cranial base (right-hand margin of the figure in this orientation), where it appears to be rolling up the paper on which the image is printed rather than telling us anything useful about the gradient of the interpolation there. The same impression of an unreal scrolling is apparent in applications to several of the Marcus mammalian crania. Three examples are displayed here, for three of the forms extreme in some of the panels in Figure 20: Elephant, Beaver, and, of course, Man.

**Figure 32.**
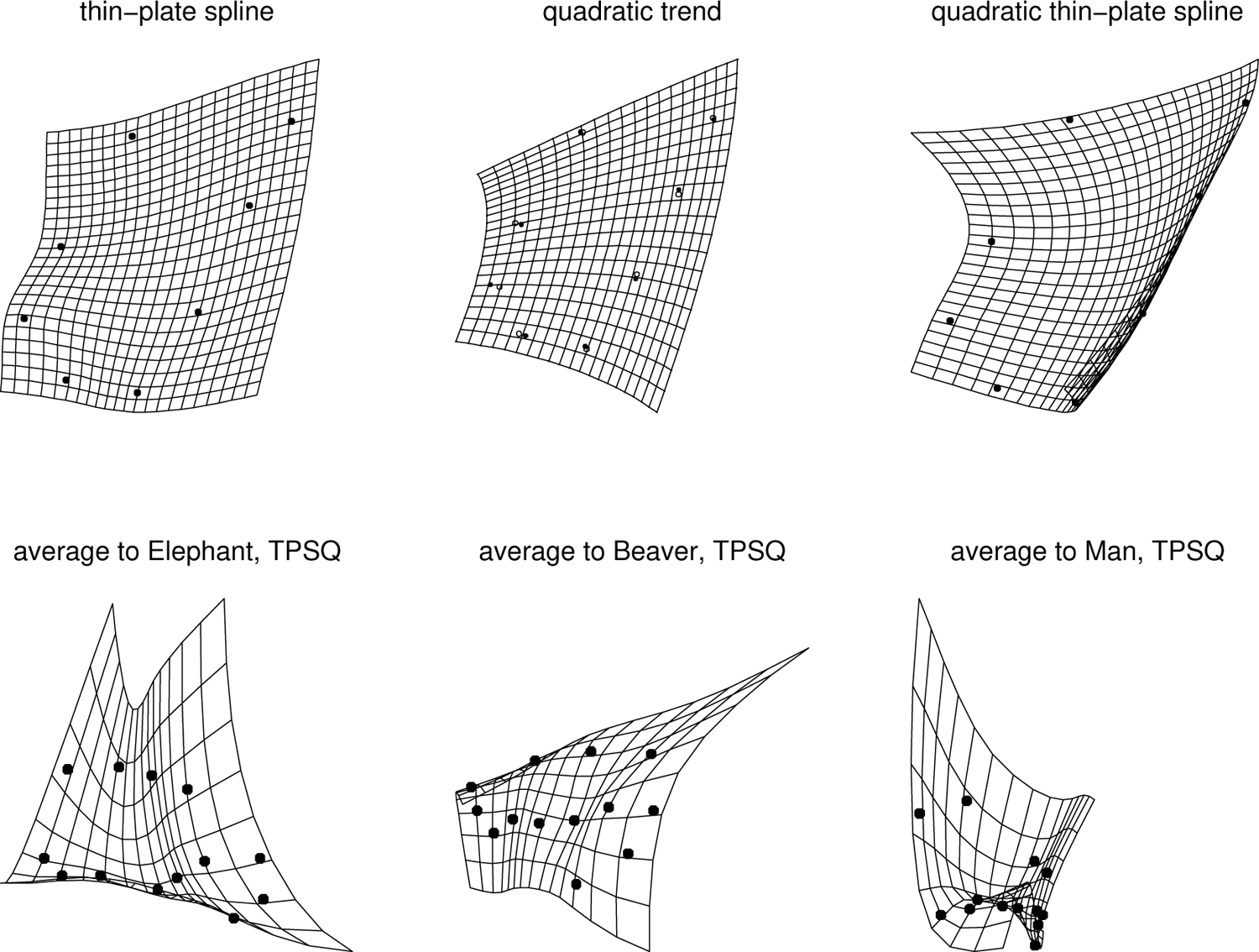
Above, three different versions of a transformation grid as applied to the fourth column of Figure 10 (the 13^◦^ rotation of a carefully chosen two-point registration of the Vilmann octagons). (left) Conventional thin-plate spline, relaxing toward linearity outside the form. (center) The quadratic trend recommended for this application, simplifying the pattern of grid lines into a vector of six parameters each a second derivative in some direction that is constant over the grid. (right) A different quadratic generated by an adjustment of the thin-plate spline formula itself. Below, examples of this quadratic thinplate spline (“TPSQ”) for three of the 55 cranial 13-gons in the revised Marcus mammalian data set. See text.

### Concluding comment: the relation of landmarks to grids

We are thereby brought face to face with a question that was implicit but mostly overlooked in the literature of thin-plate spline deformations from their first publication (Bookstein 1989) on: what is the epistemology of these grid diagrams — what reality do they so persuasively appear to be claiming? To the extent that GMM is imagined a component of the science of biology rather than a subtheme of the artificial intelligence of shape recognition or classification or the intellectual property of industrial biometric applications such as security or animal husbandry, I believe it is important to align with Thompson rather than the current GMM synthesis on this matter: to have the role of GMM be to *generate* hypotheses, not to test them, so that the job of the grid diagrams per se is specifically to serve as graphical metaphors of biological explanations that go on to be tested experimentally, as Przibram argued fully a century ago, or at least by explicit confrontation with data resources exogenous to morphology. The finding in Figure 10, repeated in the upper central panel of Figure 32, is thus a provocation to generate some explanation of why four of the six parameters of a quadratic trend fit are indeed close to zero in this growth data set. Likely their vanishing, and the resulting bilinearity of the growth deformation, is saying something important about the regulation of rodent neurocranial form; what might that bilinearity be announcing?

Similar to this concern about the biological meaning of the transformation grids is a concern for the meaning of the data that delimit them. Sneath (1967) proceeded past the six-coefficient stage of these quadratic transformations to consider the next level as well, the ten-coefficient stage that supplements the *Q* of equations (3) or (4) with terms in *x*^3^*, x*^2^*y*, *xy*^2^, and *y*^3^. However, I think it would be premature to proceed with parameterization of this model or other higher-order coefficient spaces. Rather, the GMM community should turn to a serious, principled probe into what we mean by a “configuration of landmarks” in the first place. How are these lists constructed, and when do we decide to stop adding landmarks to them, or even to delete some? The relation of landmark points to the theory of homology has hardly been explored since the pioneering work of Nicholas Jardine more than fifty years ago (Jardine, 1969). And what about semilandmarks, those representations of curving form that can number in the thousands or even higher now that powerful software can run almost autonomously on affordable multicore machines? Before extending the length of these coefficient vectors further I think the field ought to decide what it means to *be* a configuration, for instance, how and where boundaries are to be drawn between the global analysis of one composite form and a series of analyses of diverse component subforms followed by their synthesis using some non-morphometric method such as partial least squares or machine learning.

If *simplification of a grid report until it is comprehensible* is indeed the ultimate biological role of a revised GMM, none of today’s standard GMM tools — not Procrustes shape coordinates, not their principal components, not the thin-plate splines that purport to visualize their multivariate patterns — is capable of supplying the appropriate rhetoric for the announcement of empirical pattern findings in a language suggestive of future organismal explanations. Their occasional usefulness in tasks of discrimination or classification notwithstanding, none of this technology accommodates information about the spatial arrangement of landmarks along with the spatial disposition of the curves that summarize the comparisons of the anatomical regions they delineate. Neither the sums of squares that the Procrustes method mimimizes nor the sums of squared loadings that the method of principal components maximizes are the appropriate currency of a biometric pattern analysis; nor is the bending energy of a thin-plate spline. What is required instead is an interpretation of the landmark configurations as expressions of causes or effects of their anatomical organization in life, prior to being digitized. Because the current GMM toolkit is not capable of serving biological science in this way, I put forward the quadratic technology of this article as a possible first step in liberating GMM from the current strait-jacket of Procrustes shape coordinates and thin-plate splines into which the synthesis of the 1990’s has inadvertently cast it.

## Supporting information

supplement: 55 4-panel dashboards

## Acknowledgements

I am grateful to Joe Felsenstein (University of Washington), Norm Macleod (Nanjing University), and Jim Rohlf (Stony Brook University) for extensive comments on earlier versions of the argument here. There is no grant support associated with this work at present and no conflicts of interest to be reported. Thanks go as well to Erika Hingst-Zaher (Instituto Butantan, São Paulo) for the revised Marcus data set she shared with me ten years ago in connection with the preparation of the work cited here as Bookstein 2018.

## Supplement

Analogues to Figures 14 through 16 and 25 through 27 for all 55 of the specimens in the data set of Section IV.

## Footnotes

1 The symbol ∂, read “del,” stands for the partial derivative function, the derivative of a function of two or more variables when only one at a time is allowed to vary.

2 To convert a predicted set of cardinal diameter endpoints, such as these principal component loading vector, into a quadratic trend, one must produce the values of all six parameters of that trend. The coefficients of *x*^2^ and *y*^2^ for both of the target coordinates are explicit in the locations of the endpoints of the predicted cardinal diameter at angles 0^◦^ and 90^◦^ to the baseline. The remaining two coefficients, pertaining to the mixed partial 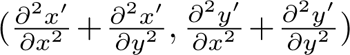 in the two dimensions of the target configuration, can be estimated as half the difference of the two directional derivatives in the (*x* + *y*) and (*x* − *y*) directions (Abramowitz and Stegun 1964, formula 25.3.26). The multiplier here of ½ adjusts the coefficient ¼ of their formula 25.3.26 to account for the facts that the radii of the circle of diameters here is only 1/√2 of the spacing of the grid points in that formula and that the dependence of the estimated derivative on these differences goes as the inverse square of their separation.

3 The operator Δ is sometimes written ∇^2^, where ∇ is the operator the lexicons name “nabla” that stands for either “gradient” (if it is applied to a scalar function) or “divergence” (if a vector). The Laplacian of a scalar function *f* is the divergence of its gradient, that is, Δ*f* ≡ ∇ · (∇*f*), which, by an abuse of notation, gets to be written as ∇^2^. roughness vector that is the same everywhere.)

