## supplement: 55 4-panel dashboards for "Quadratic trends: a morphometric tool both old and new"

Fred L. Bookstein

University of Vienna, University of Washington

**Supplement**

Four-panel figures on the design of Figure 13, 14, 15 for each of the 55 specimens of mammals in the data set of Section IV. Pages are in the order of the original file as sent by Erika Hingst-Zaher to the author in 2013.

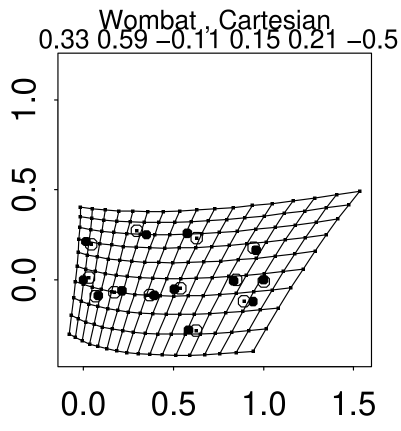
Wombat , Cartesian
0.33 0.59 −0.11 0.15 0.21 −0.5
1.0
0.5
0.0
0.0 0.5 1.0 1.5

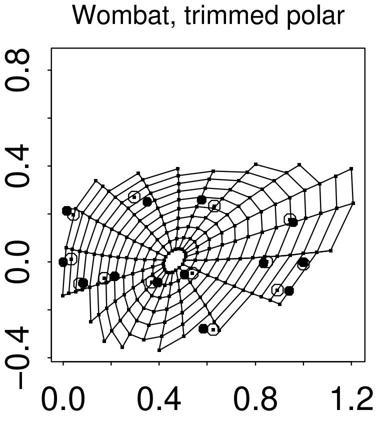
Wombat, trimmed polar
0.8
0.4
0.0
−0.4
0.0 0.4 0.8 1.2

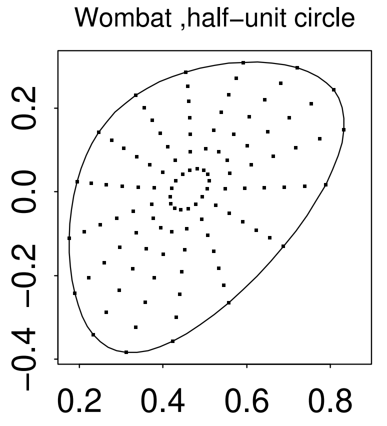
Wombat ,half−unit circle
0.2
0.0
−0.2
−0.4
0.2 0.4 0.6 0.8

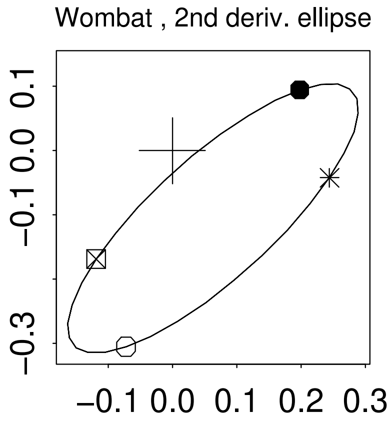
Wombat , 2nd deriv. ellipse
0.1
0.0
−0.1
−0.3
−0.1 0.0 0.1 0.2 0.3

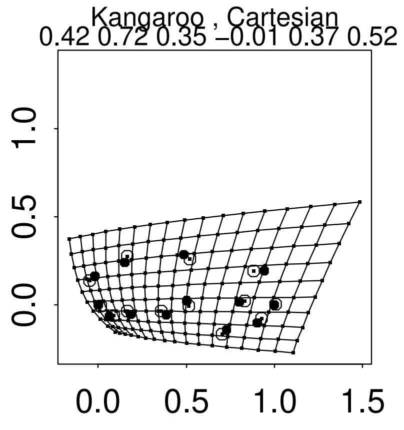
Kangaroo , Cartesian
0.42 0.72 0.35 −0.01 0.37 0.52
0.0 0.5 1.0 1.5
0.0 0.5 1.0 1.5

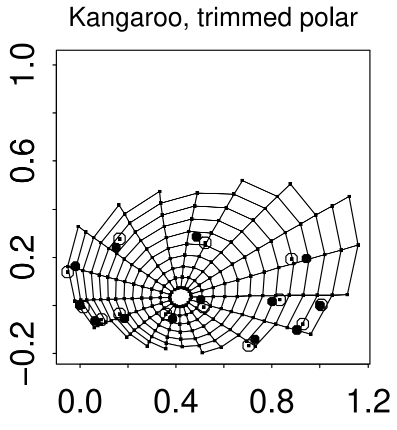
Kangaroo, trimmed polar
−0.2 0.2 0.6 1.0
0.0 0.4 0.8 1.2

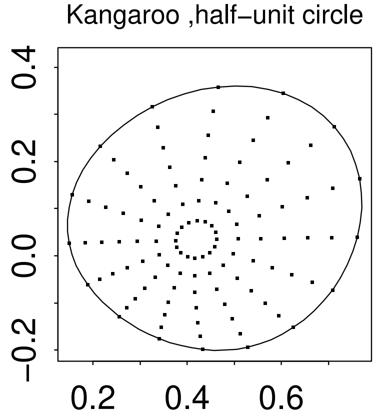
Kangaroo ,half−unit circle
−0.2 0.0 0.2 0.4
0.2 0.4 0.6

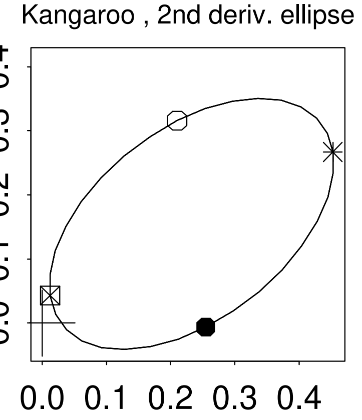
Kangaroo , 2nd deriv. ellipse
0.0 0.1 0.2 0.3 0.4
0.0 0.1 0.2 0.3 0.4

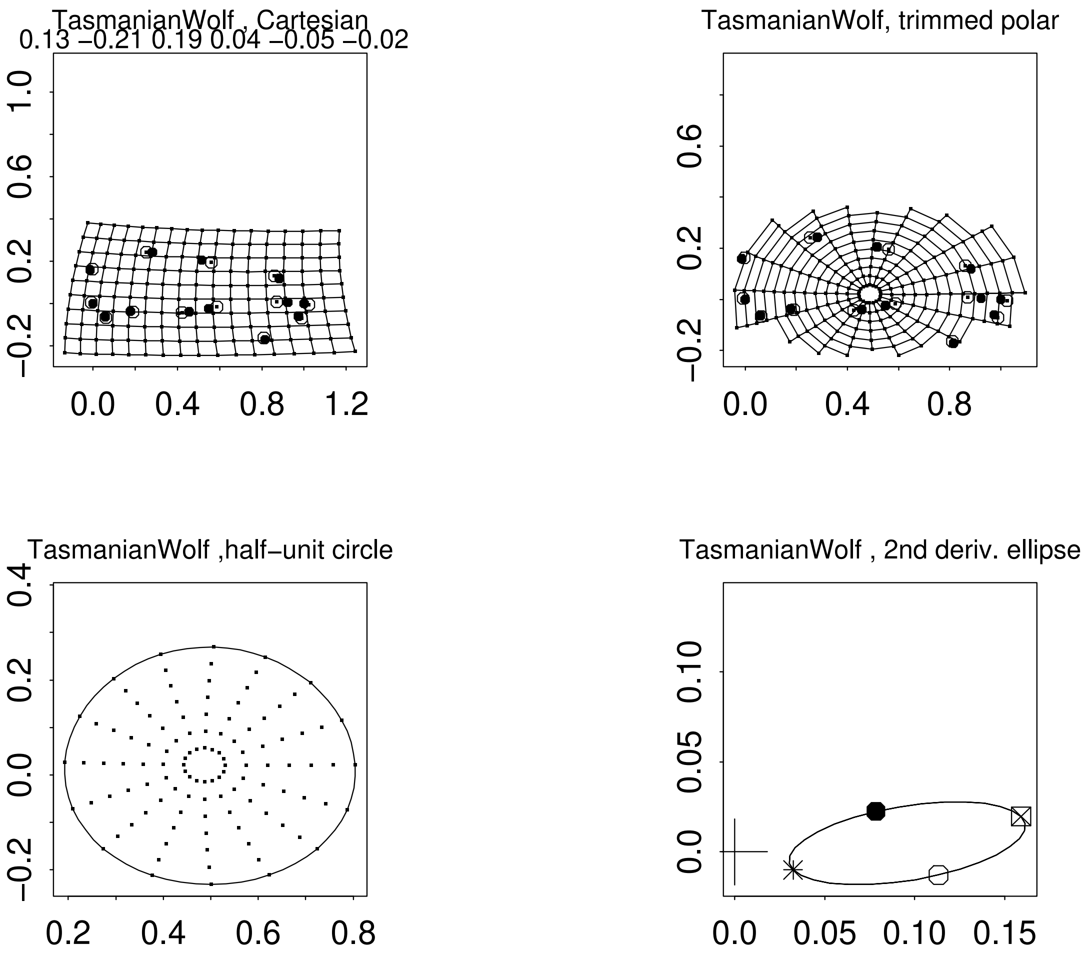
TasmanianWolf , Cartesian
0.13 −0.21 0.19 0.04 −0.05 −0.02
TasmanianWolf, trimmed polar
TasmanianWolf ,half−unit circle
TasmanianWolf , 2nd deriv. ellipse

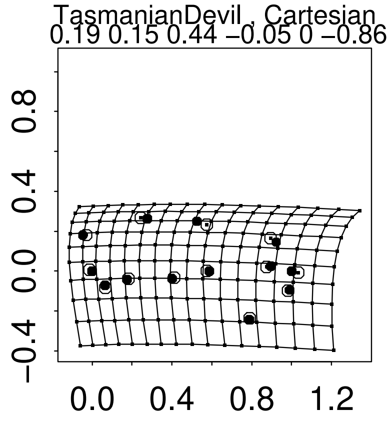
TasmanianDevil , Cartesian
0.19 0.15 0.44 −0.05 0 −0.86
0.8
0.4
0.0
−0.4
0.0 0.4 0.8 1.2

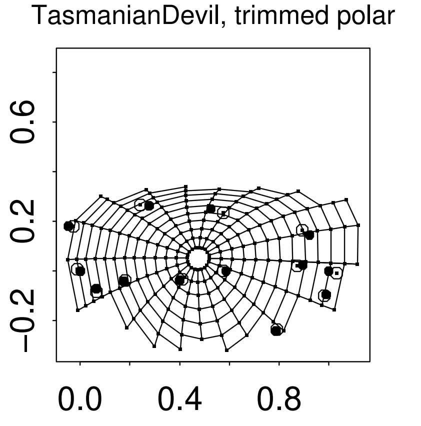
TasmanianDevil, trimmed polar
0.6
0.2
−0.2
0.0 0.4 0.8

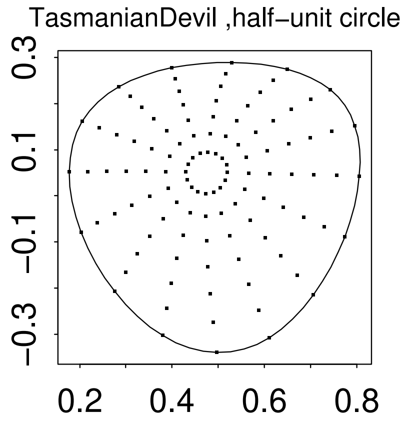
TasmanianDevil ,half−unit circle
0.3
0.1
−0.1
−0.3
0.2 0.4 0.6 0.8

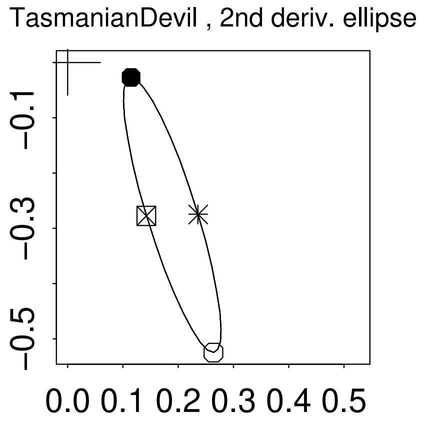
TasmanianDevil , 2nd deriv. ellipse
−0.1
−0.3
−0.5
0.0 0.1 0.2 0.3 0.4 0.5

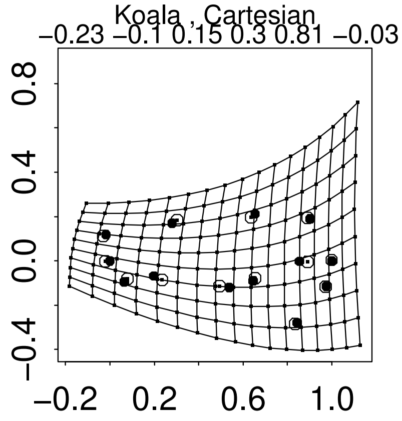
Koala , Cartesian
−0.23 −0.1 0.15 0.3 0.81 −0.03
−0.4 0.0 0.4 0.8
−0.2 0.2 0.6 1.0

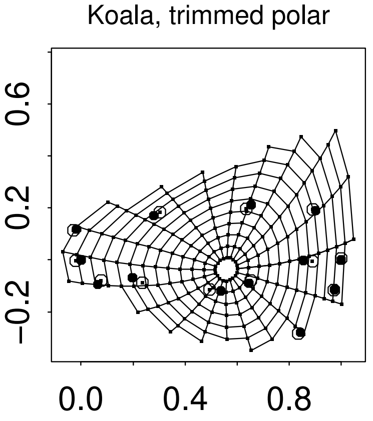
Koala, trimmed polar
−0.2 0.2 0.6
0.0 0.4 0.8

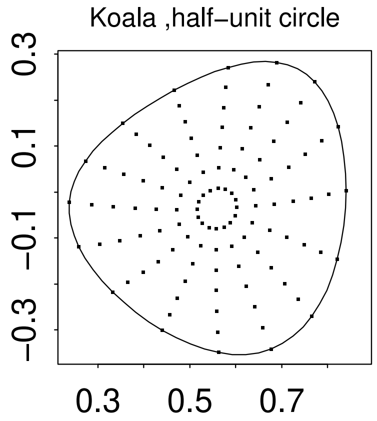
Koala ,half−unit circle
−0.3 −0.1 0.1 0.3
0.3 0.5 0.7

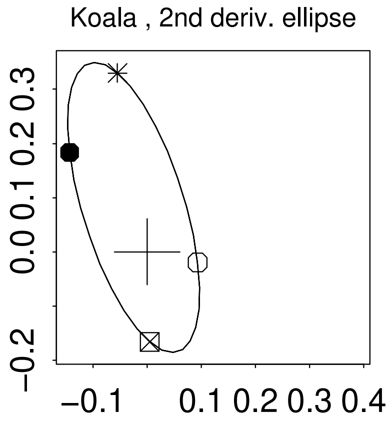
Koala , 2nd deriv. ellipse
−0.2 0.0 0.1 0.2 0.3
−0.1 0.1 0.2 0.3 0.4

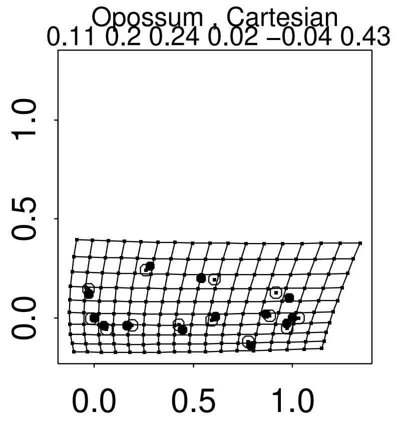
Opossum , Cartesian
0.11 0.2 0.24 0.02 −0.04 0.43
1.0
0.5
0.0
0.0
0.5
1.0

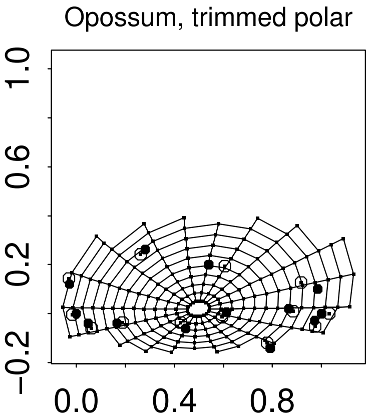
Opossum, trimmed polar
1.0
0.6
0.2
−0.2
0.0
0.4
0.8

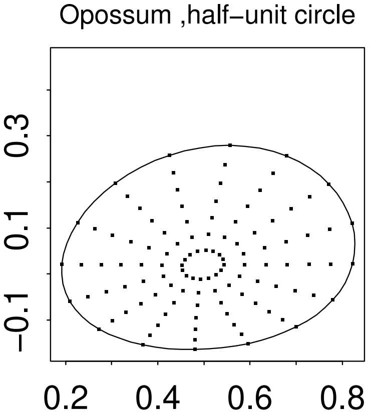
Opossum ,half−unit circle
0.3
0.1
−0.1
0.2
0.4
0.6
0.8

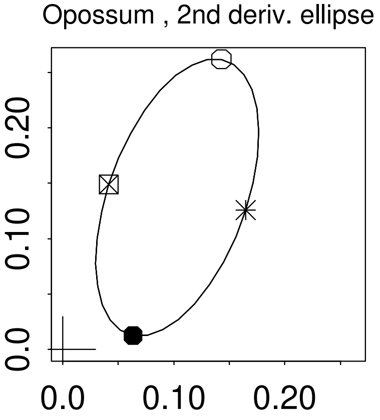
Opossum , 2nd deriv. ellipse
0.20
0.10
0.0
0.0
0.10
0.20

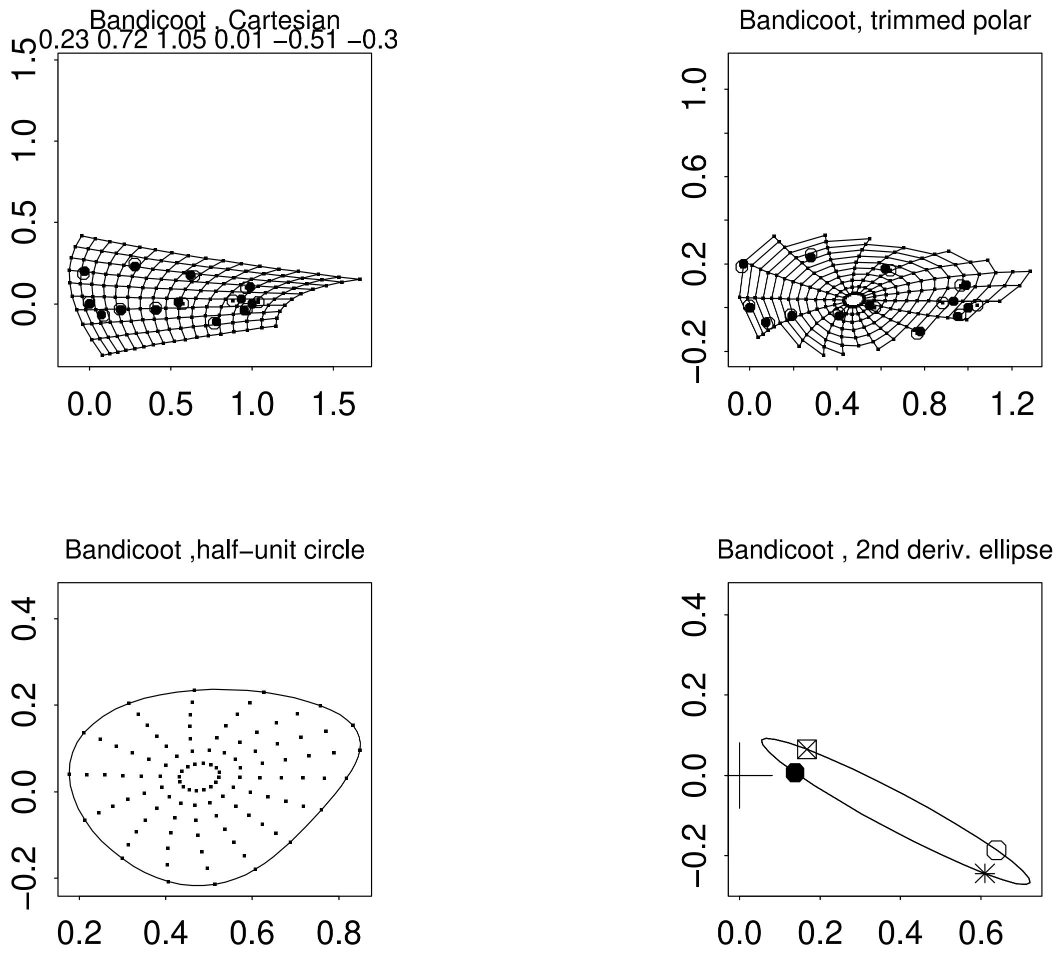
Bandicoot , Cartesian
0.23 0.72 1.05 0.01 −0.51 −0.3
Bandicoot, trimmed polar
Bandicoot ,half−unit circle
Bandicoot , 2nd deriv. ellipse

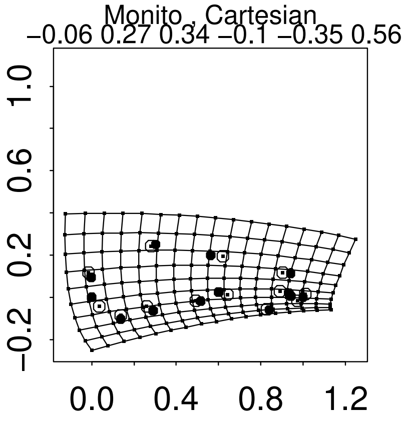
Monito , Cartesian
−0.06 0.27 0.34 −0.1 −0.35 0.56
−0.2
0.2
0.6
1.0
0.0
0.4
0.8
1.2

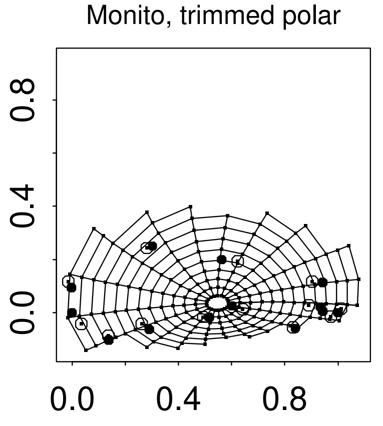
Monito, trimmed polar
0.0
0.4
0.8
0.0
0.4
0.8

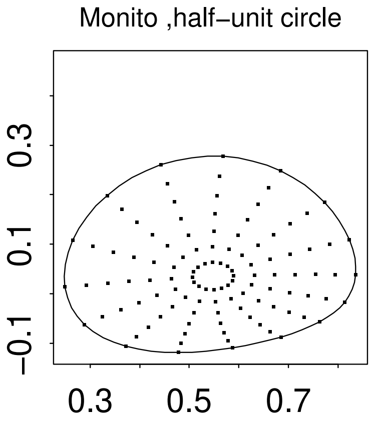
Monito ,half−unit circle
−0.1
0.1
0.3
0.3
0.5
0.7

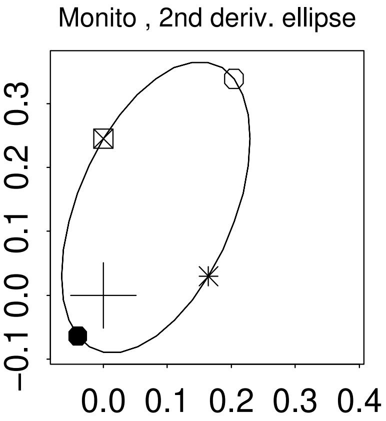
Monito , 2nd deriv. ellipse
−0.1
0.0
0.1
0.2
0.3
0.0
0.1
0.2
0.3
0.4

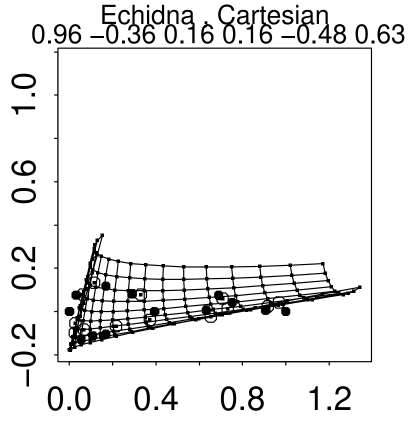
Echidna , Cartesian
0.96 −0.36 0.16 0.16 −0.48 0.63
1.0
0.6
0.2
−0.2
0.0
0.4
0.8
1.2

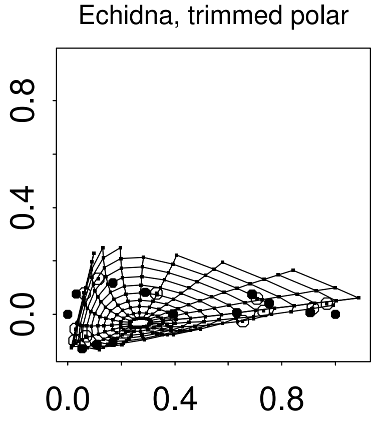
Echidna, trimmed polar
0.8
0.4
0.0
0.0
0.4
0.8

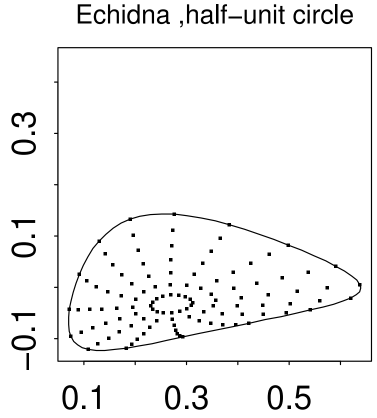
Echidna ,half−unit circle
0.3
0.1
−0.1
0.1
0.3
0.5

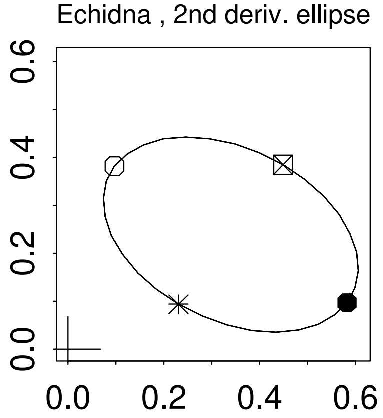
Echidna , 2nd deriv. ellipse
0.6
0.4
0.2
0.0
0.0
0.2
0.4
0.6

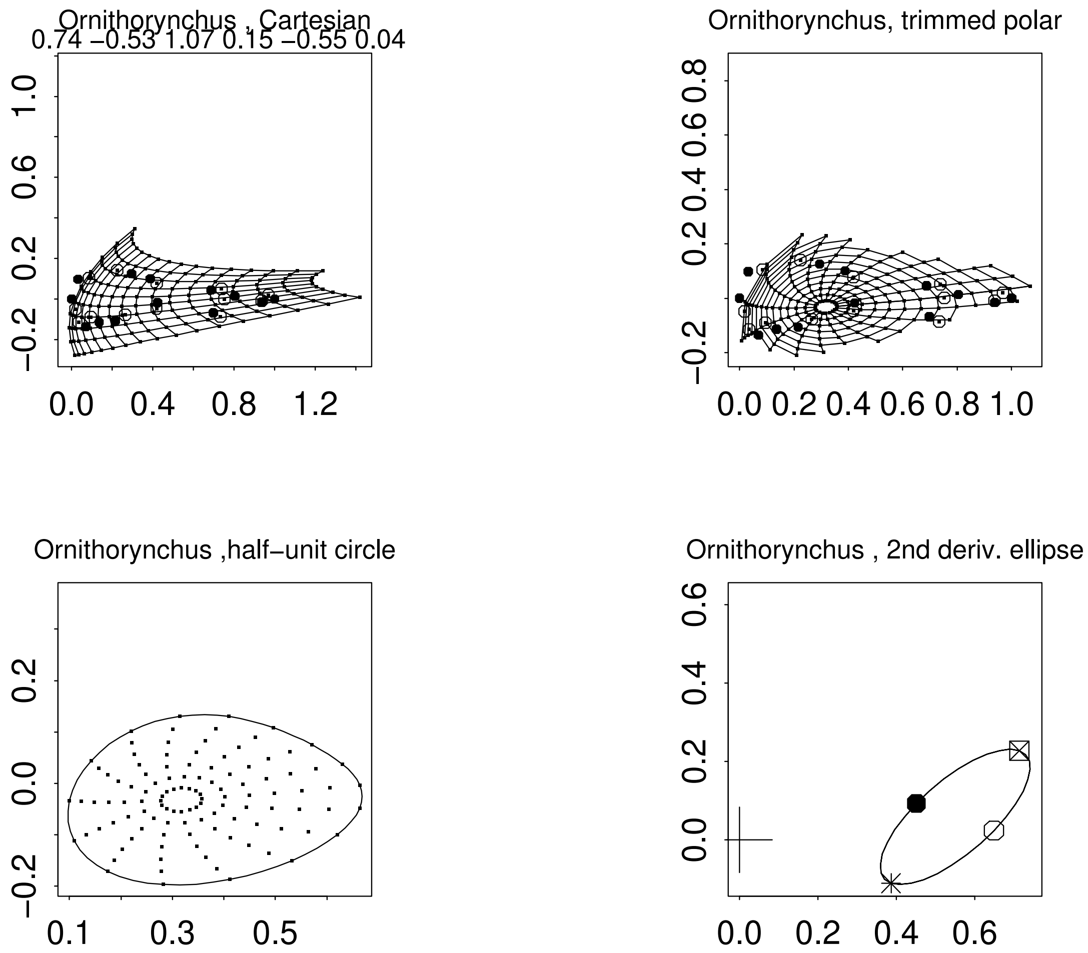

Ornithorynchus , Cartesian
0.74 −0.53 1.07 0.15 −0.55 0.04
Ornithorynchus, trimmed polar
Ornithorynchus ,half−unit circle
Ornithorynchus , 2nd deriv. ellipse

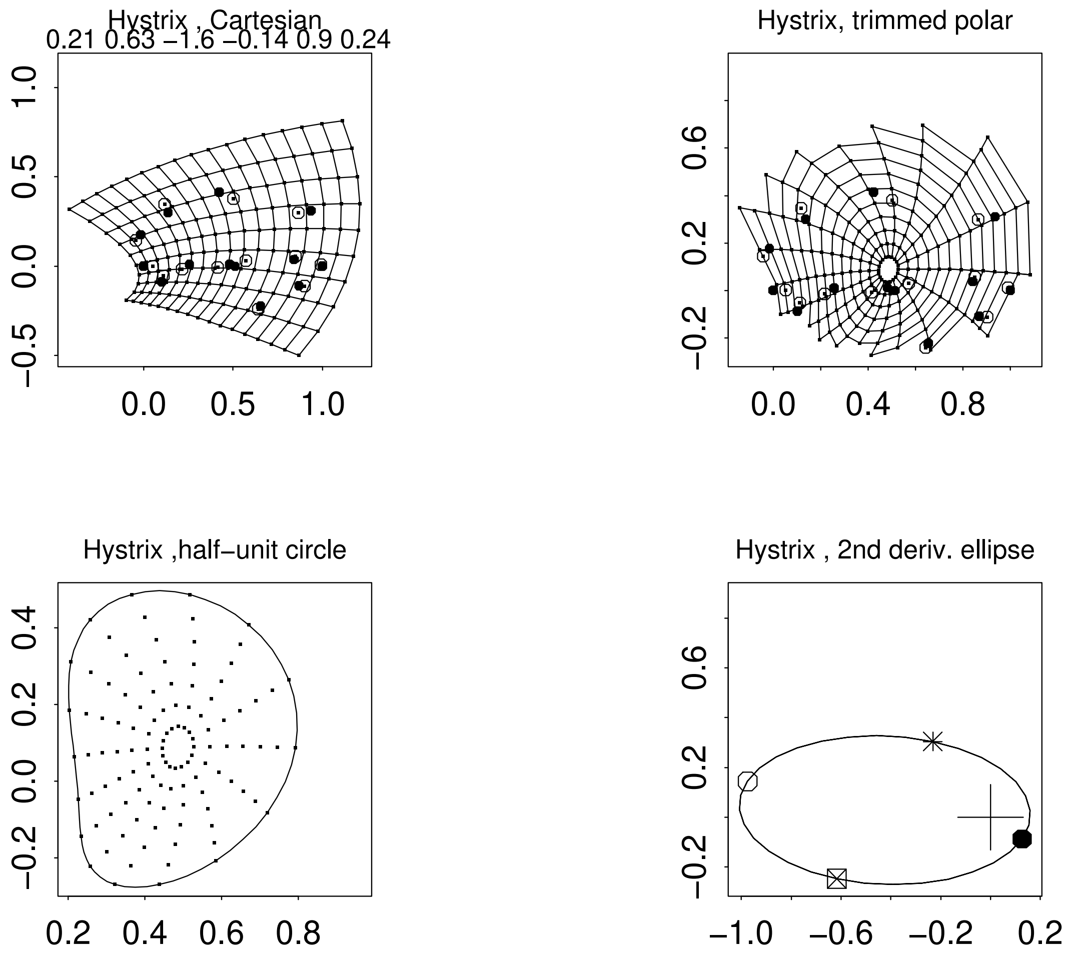
Hystrix , Cartesian
0.21 0.63 −1.6 −0.14 0.9 0.24
Hystrix, trimmed polar
Hystrix ,half−unit circle
Hystrix , 2nd deriv. ellipse

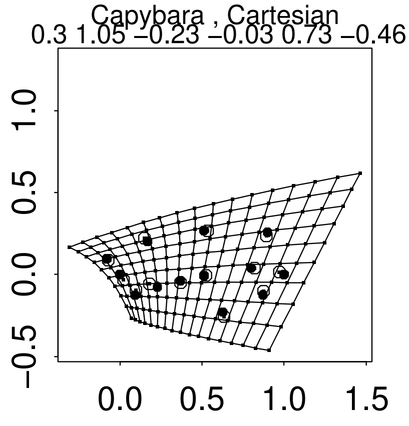
Capybara , Cartesian
0.3 1.05 −0.23 −0.03 0.73 −0.46
−0.5 0.0 0.5 1.0
0.0 0.5 1.0 1.5

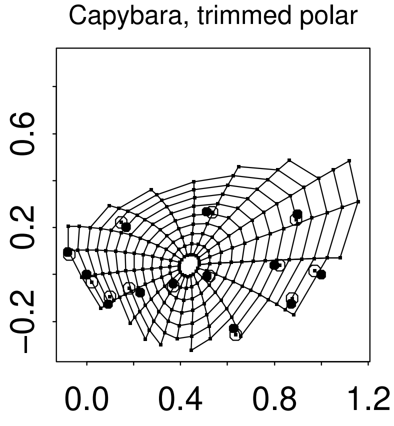
Capybara, trimmed polar
−0.2 0.2 0.6
0.0 0.4 0.8 1.2

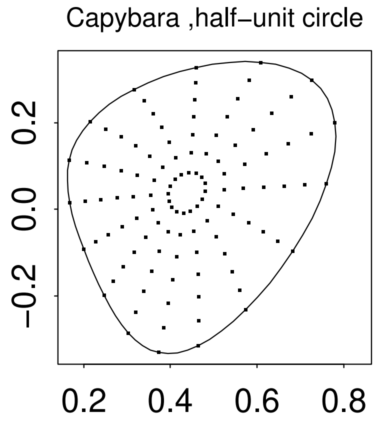
Capybara ,half−unit circle
−0.2 0.0 0.2
0.2 0.4 0.6 0.8

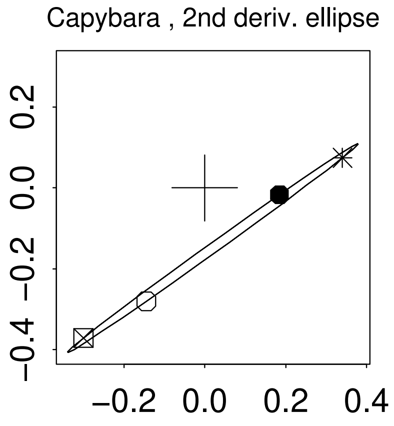
Capybara , 2nd deriv. ellipse
−0.4 −0.2 0.0 0.2
−0.2 0.0 0.2 0.4

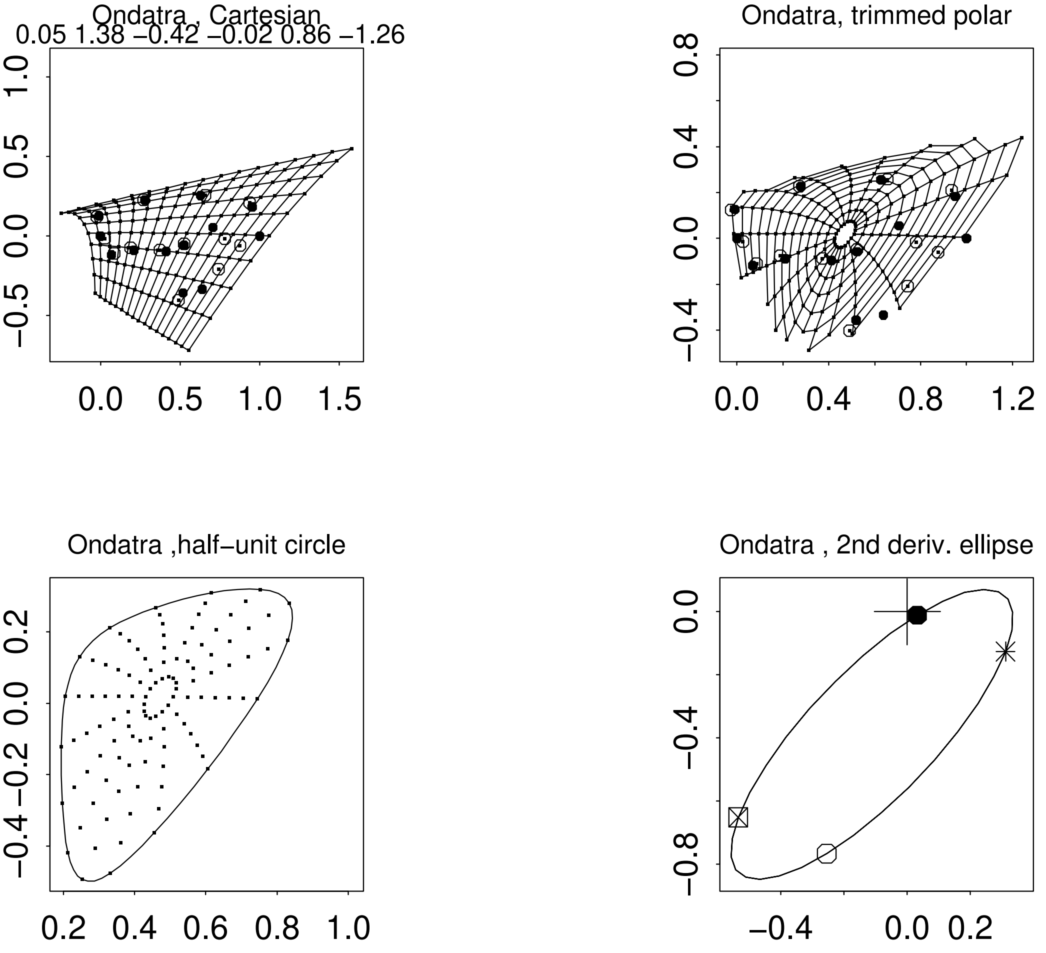
Ondatra , Cartesian
0.05 1.38 −0.42 −0.02 0.86 −1.26
−0.5 0.0 0.5 1.0
0.0 0.5 1.0 1.5
Ondatra, trimmed polar
−0.4 0.0 0.4 0.8
0.0 0.4 0.8 1.2
Ondatra ,half−unit circle
−0.4 −0.2 0.0 0.2
0.2 0.4 0.6 0.8 1.0
Ondatra , 2nd deriv. ellipse
−0.8 −0.4 0.0
−0.4 0.0 0.2

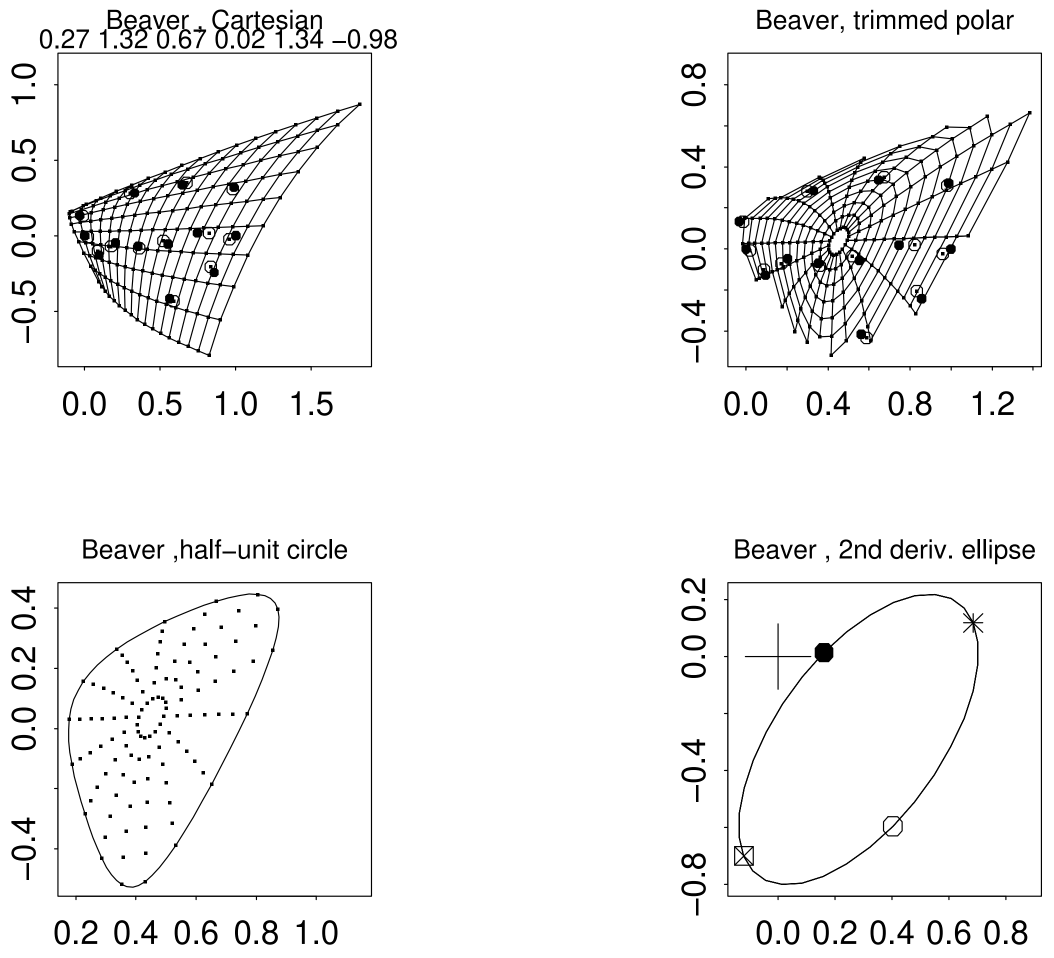

Beaver , Cartesian
0.27 1.32 0.67 0.02 1.34 −0.98
Beaver, trimmed polar
Beaver ,half−unit circle
Beaver , 2nd deriv. ellipse

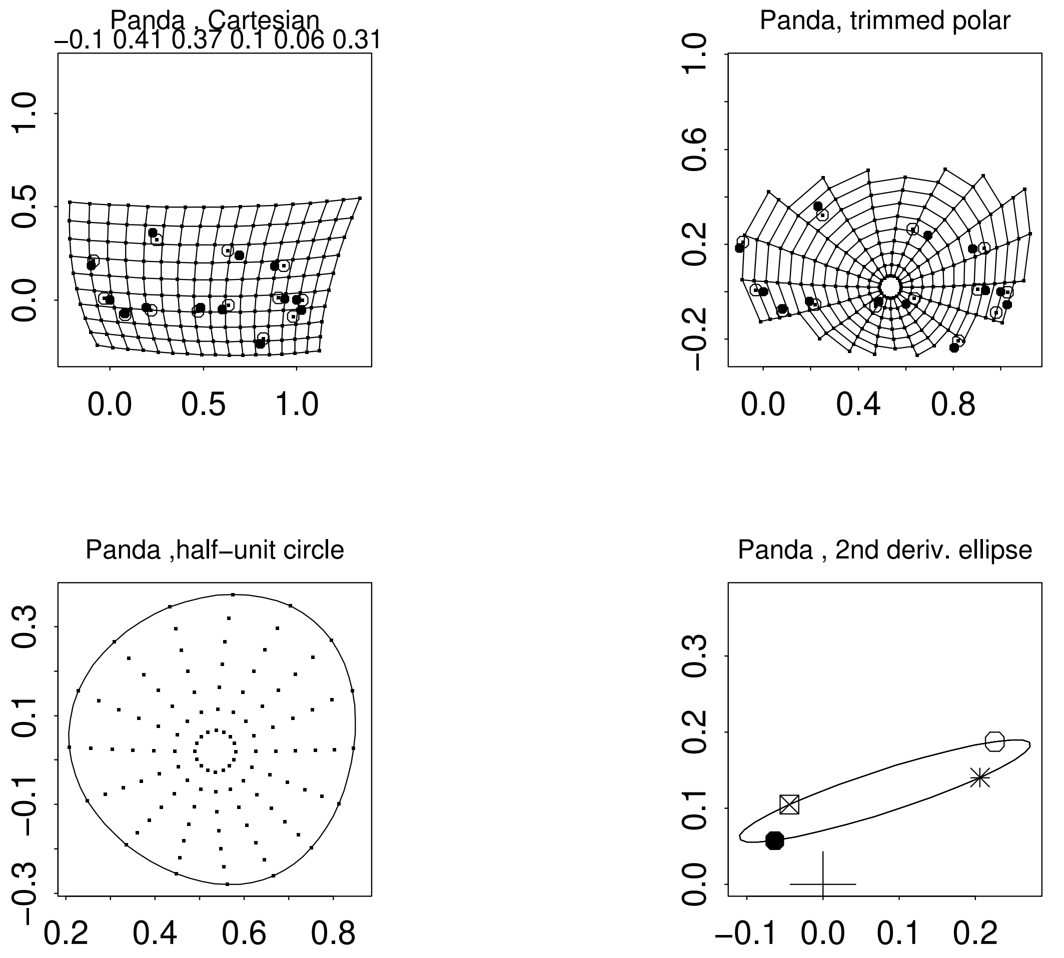

Panda , Cartesian
−0.1 0.41 0.37 0.1 0.06 0.31
0.0 0.5 1.0
Panda, trimmed polar
−0.2 0.2 0.6 1.0
0.0 0.4 0.8
Panda ,half−unit circle
−0.3 −0.1 0.1 0.3
0.2 0.4 0.6 0.8
Panda , 2nd deriv. ellipse
0.0 0.1 0.2 0.3
−0.1 0.0 0.1 0.2

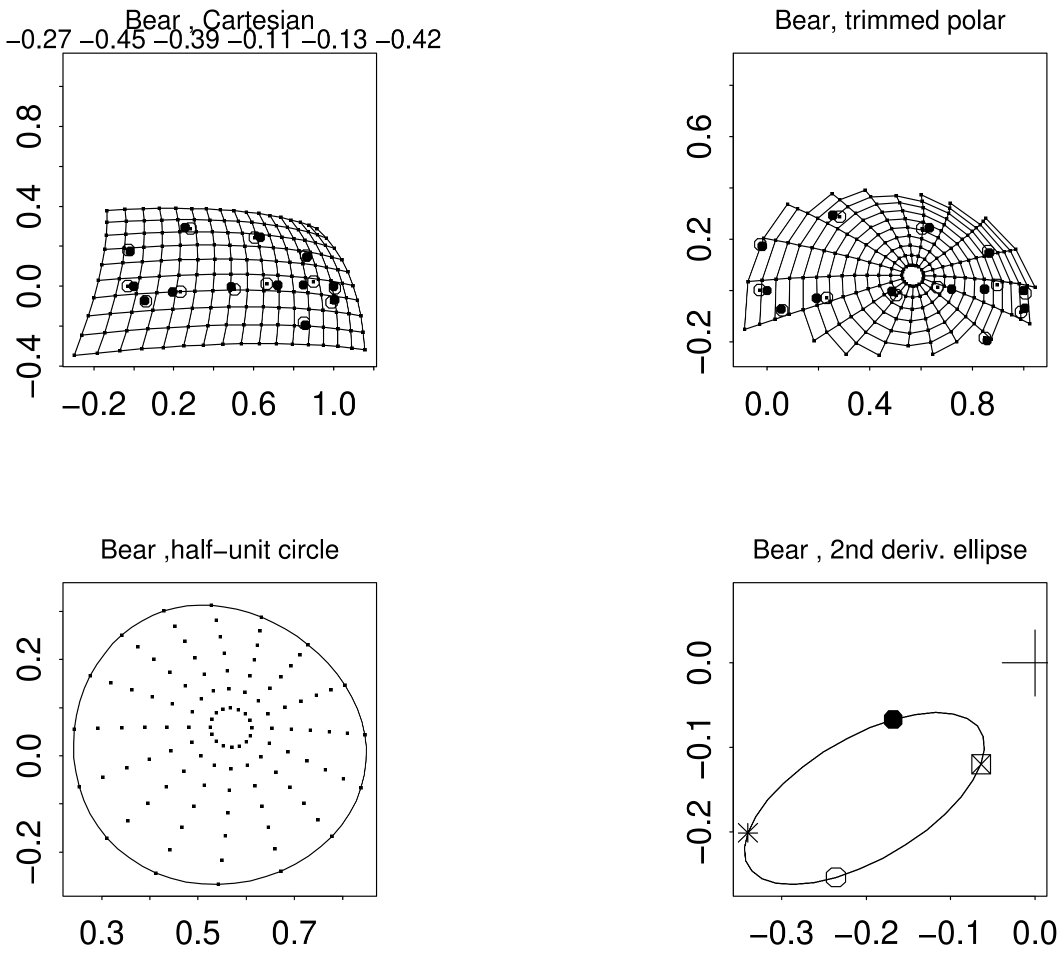
Bear , Cartesian
−0.27 −0.45 −0.39 −0.11 −0.13 −0.42
Bear, trimmed polar
Bear ,half−unit circle
Bear , 2nd deriv. ellipse

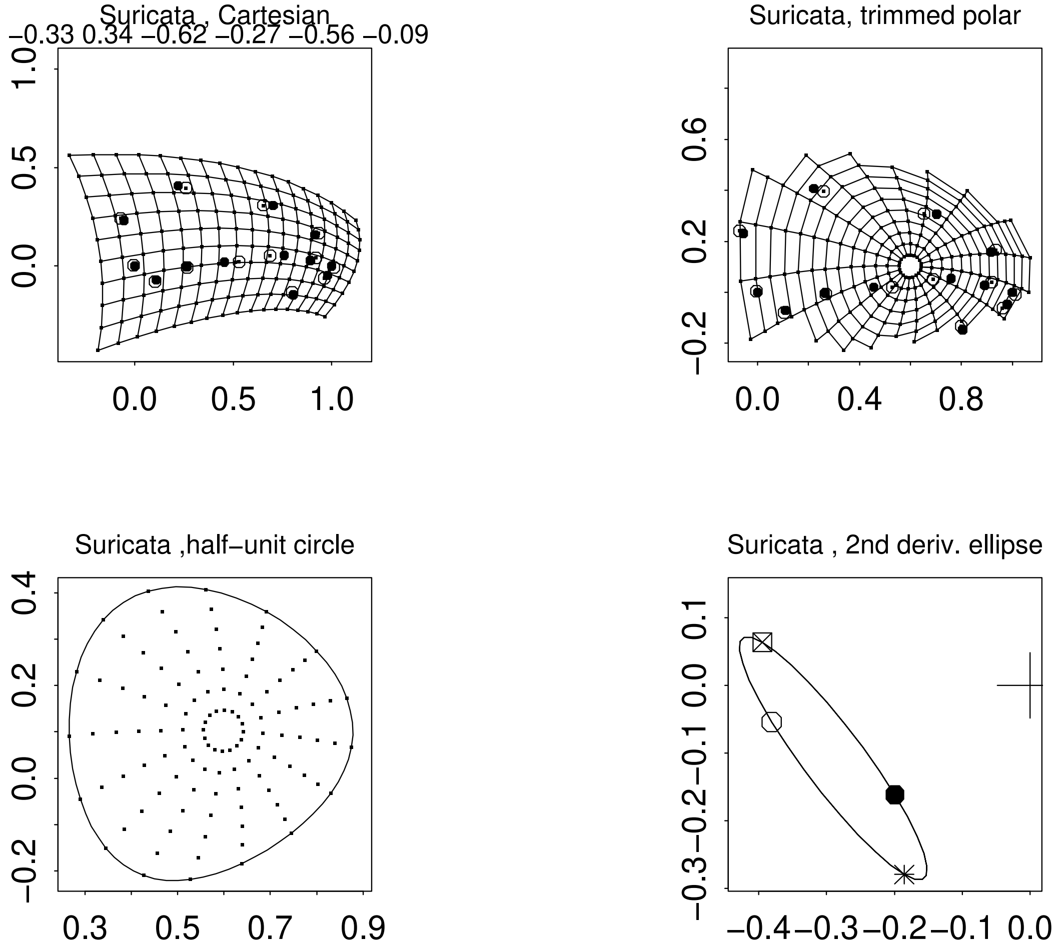
Suricata , Cartesian
−0.33 0.34 −0.62 −0.27 −0.56 −0.09
Suricata, trimmed polar
Suricata ,half−unit circle
Suricata , 2nd deriv. ellipse

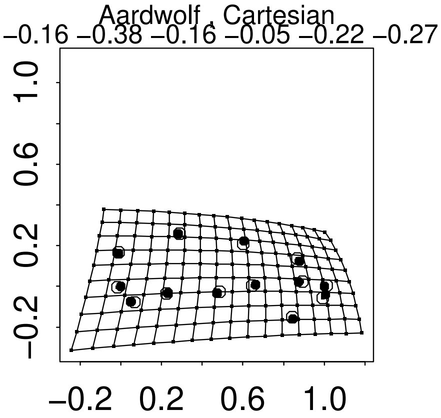
Aardwolf , Cartesian
−0.16 −0.38 −0.16 −0.05 −0.22 −0.27
1.0
0.6
0.2
−0.2
−0.2 0.2 0.6 1.0

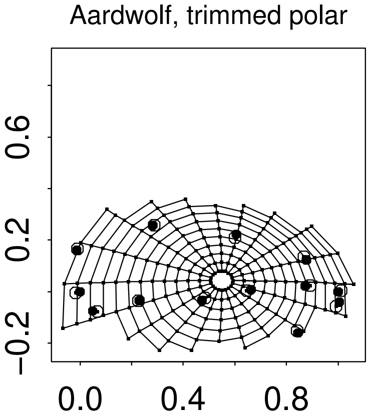
Aardwolf, trimmed polar
0.6
0.2
−0.2
0.0 0.4 0.8

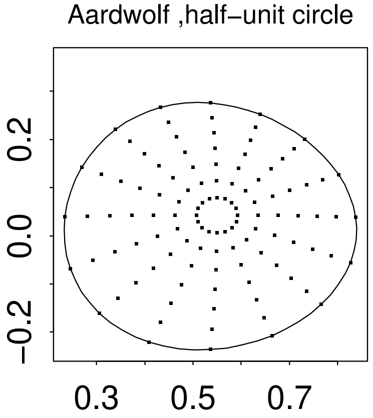
Aardwolf ,half−unit circle
0.2
0.0
−0.2
0.3 0.5 0.7

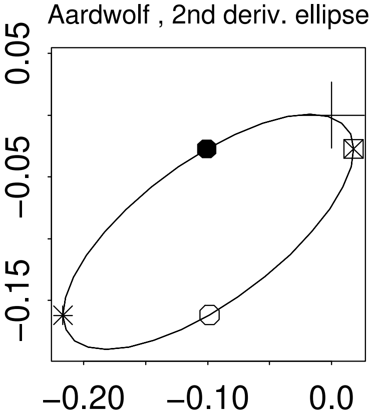
Aardwolf , 2nd deriv. ellipse
0.05
−0.05
−0.15
−0.20 −0.10 0.0

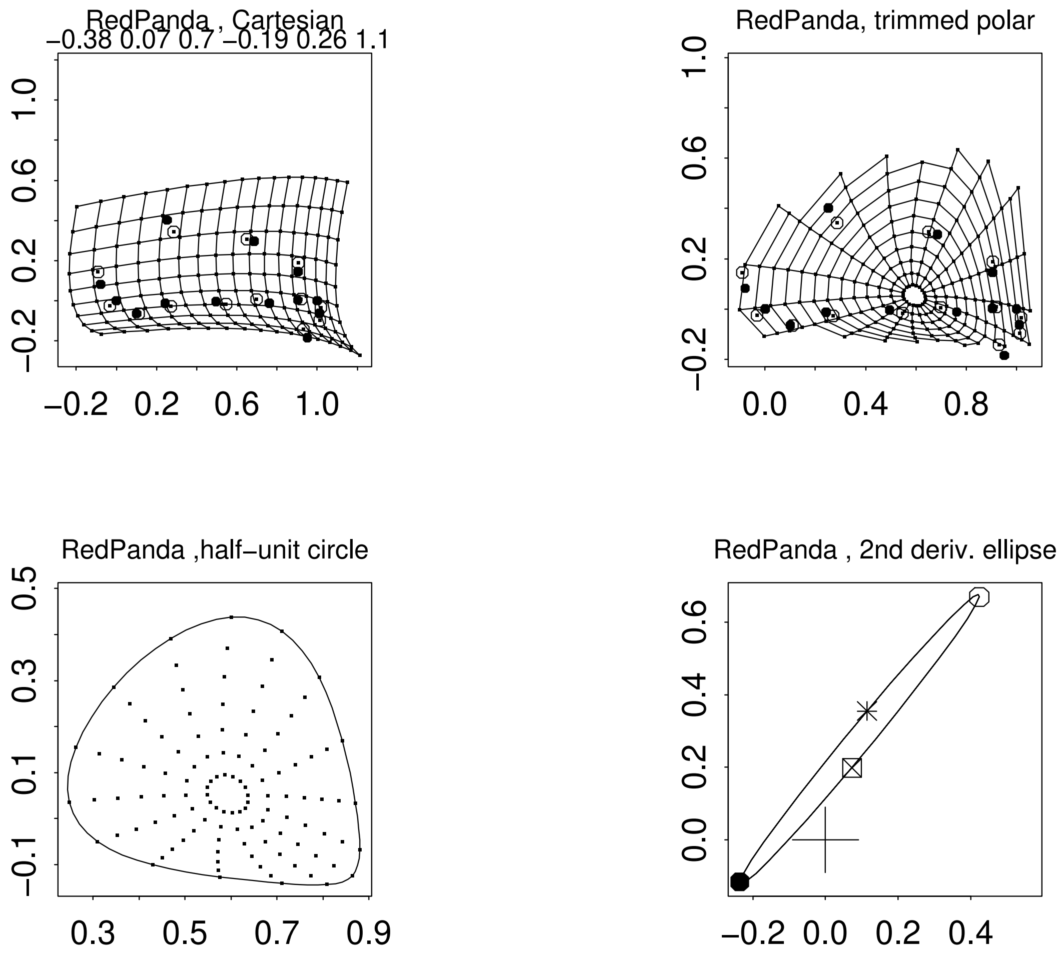
RedPanda , Cartesian
−0.38 0.07 0.7 −0.19 0.26 1.1
−0.2 0.2 0.6 1.0
−0.2 0.2 0.6 1.0
RedPanda, trimmed polar
−0.2 0.2 0.6 1.0
0.0 0.4 0.8
RedPanda ,half−unit circle
−0.1 0.1 0.3 0.5
0.3 0.5 0.7 0.9
RedPanda , 2nd deriv. ellipse
0.0 0.2 0.4 0.6
−0.2 0.0 0.2 0.4

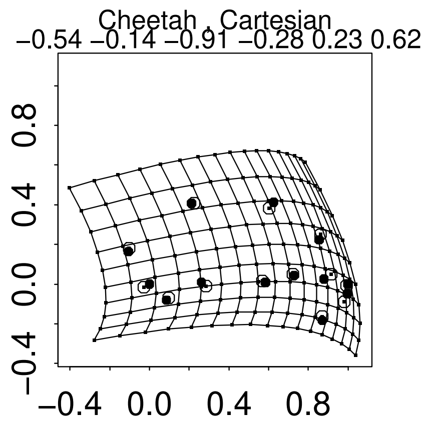
Cheetah , Cartesian
−0.54 −0.14 −0.91 −0.28 0.23 0.62
−0.4 0.0 0.4 0.8
−0.4 0.0 0.4 0.8

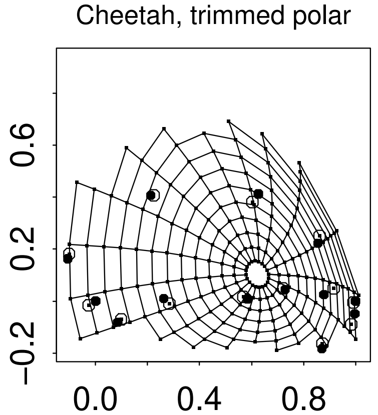
Cheetah, trimmed polar
−0.2 0.2 0.6
0.0 0.4 0.8

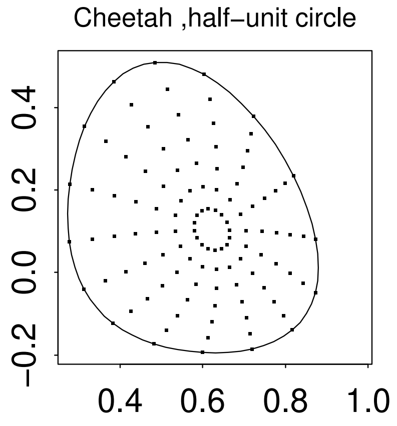
Cheetah ,half−unit circle
−0.2 0.0 0.2 0.4
0.4 0.6 0.8 1.0

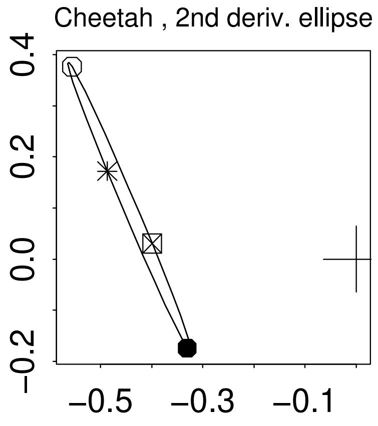
Cheetah , 2nd deriv. ellipse
−0.2 0.0 0.2 0.4
−0.5 −0.3 −0.1

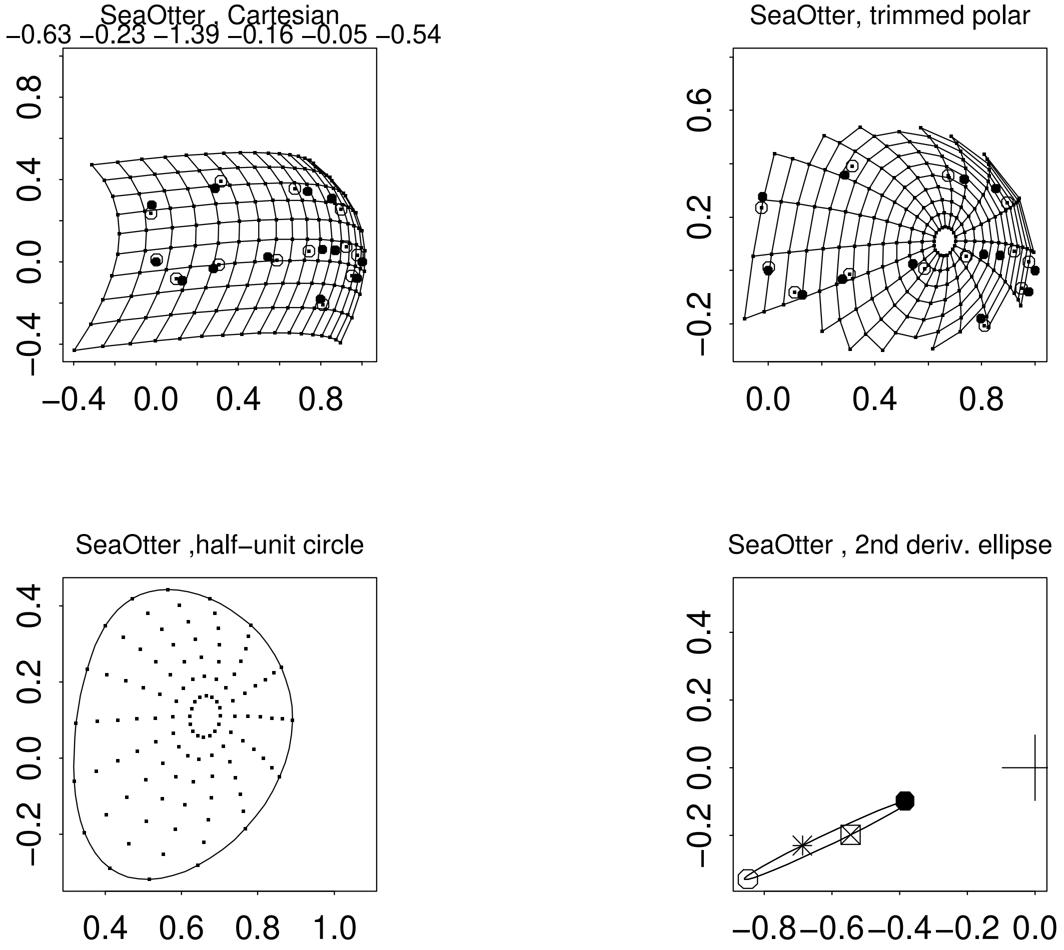
SeaOtter , Cartesian
−0.63 −0.23 −1.39 −0.16 −0.05 −0.54
−0.4 0.0 0.4 0.8
SeaOtter, trimmed polar
0.0 0.4 0.8
−0.2 0.2 0.6
SeaOtter ,half−unit circle
0.4 0.6 0.8 1.0
−0.2 0.0 0.2 0.4
SeaOtter , 2nd deriv. ellipse
−0.8 −0.6 −0.4 −0.2 0.0
−0.2 0.0 0.2 0.4

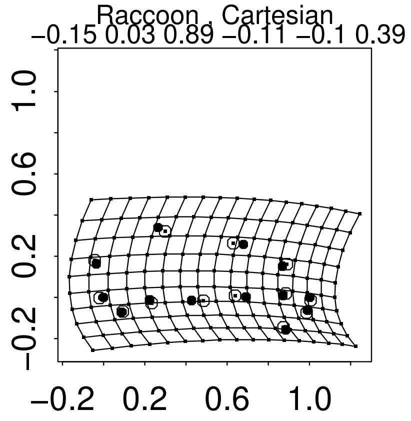
Raccoon , Cartesian
−0.15 0.03 0.89 −0.11 −0.1 0.39
−0.2 0.2 0.6 1.0
−0.2 0.2 0.6 1.0

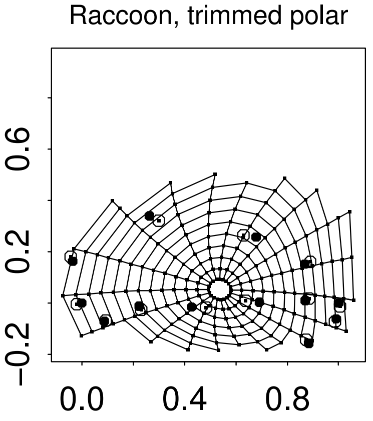
Raccoon, trimmed polar
−0.2 0.2 0.6
0.0 0.4 0.8

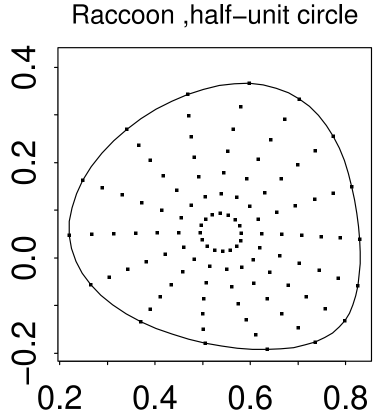
Raccoon ,half−unit circle
−0.2 0.0 0.2 0.4
0.2 0.4 0.6 0.8

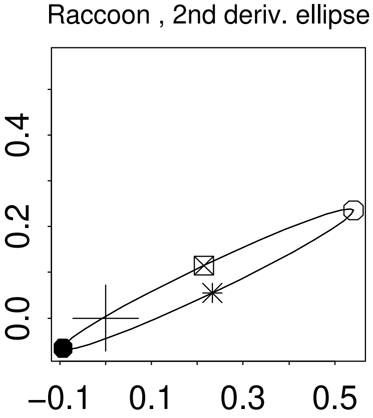
Raccoon , 2nd deriv. ellipse
0.0 0.2 0.4
−0.1 0.1 0.3 0.5

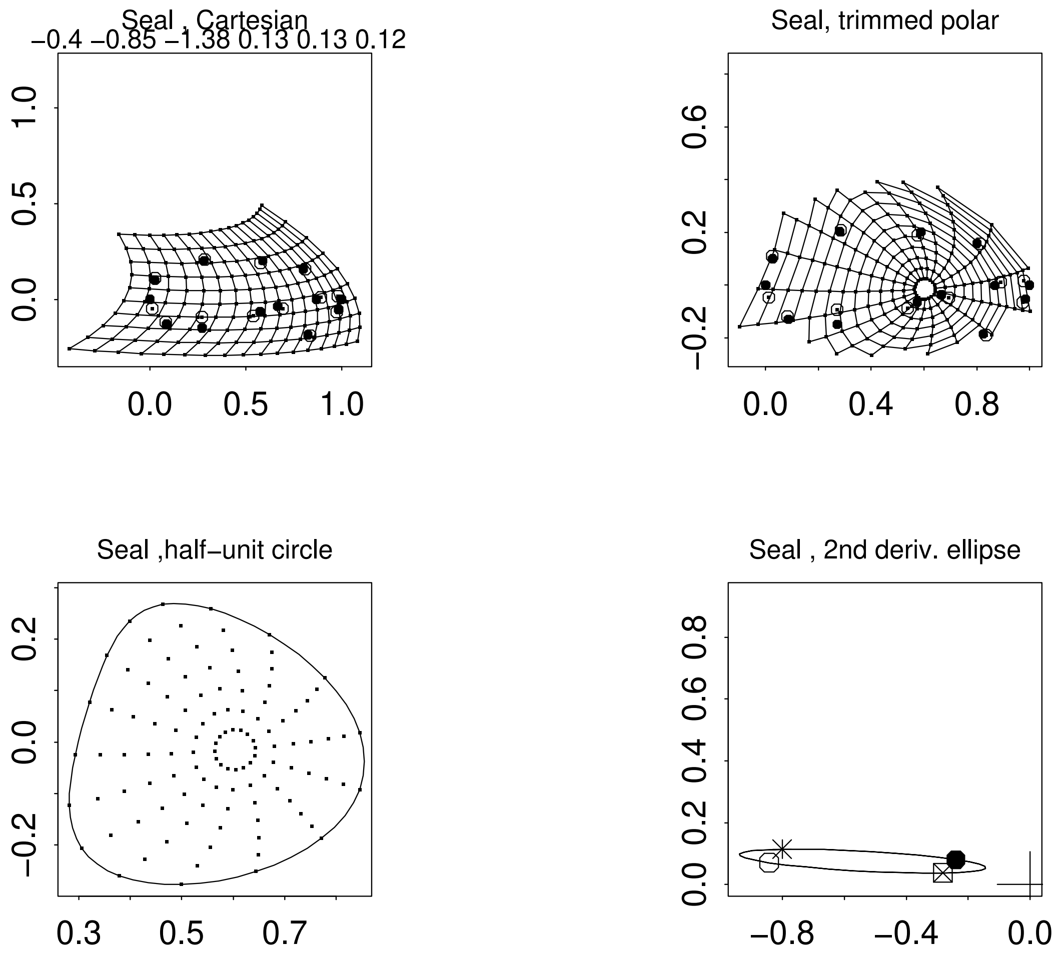
Seal , Cartesian
−0.4 −0.85 −1.38 0.13 0.13 0.12
Seal, trimmed polar
Seal ,half−unit circle
Seal , 2nd deriv. ellipse

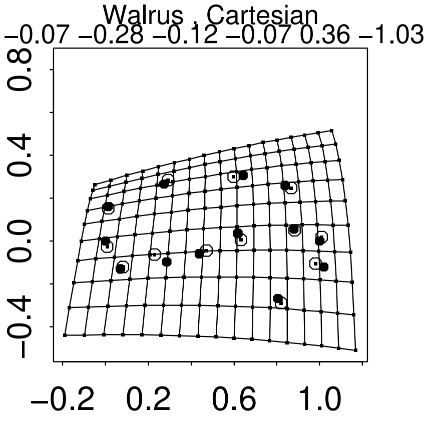
Walrus , Cartesian
−0.07 −0.28 −0.12 −0.07 0.36 −1.03
0.8
0.4
0.0
−0.4
−0.2 0.2 0.6 1.0

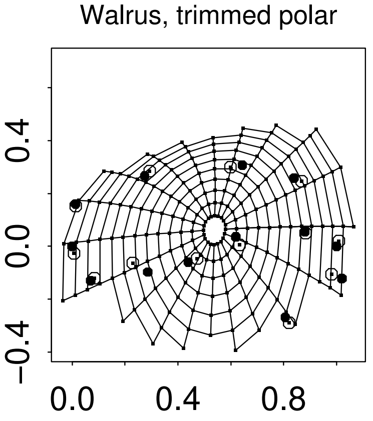
Walrus, trimmed polar
0.4
0.0
−0.4
0.0 0.4 0.8

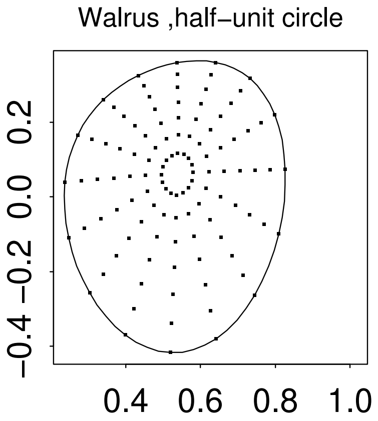
Walrus ,half−unit circle
0.2
0.0
−0.2
−0.4
0.4 0.6 0.8 1.0

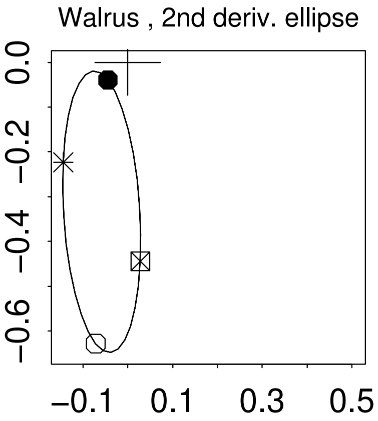
Walrus , 2nd deriv. ellipse
0.0
−0.2
−0.4
−0.6
−0.1 0.1 0.3 0.5

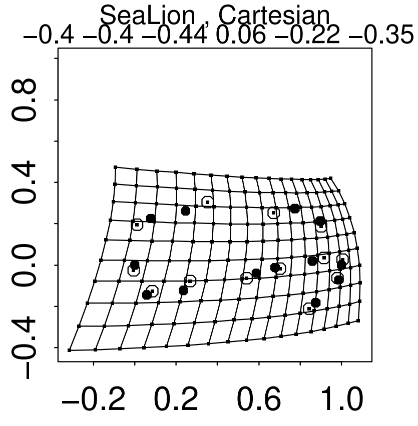
SeaLion , Cartesian
−0.4 −0.4 −0.44 0.06 −0.22 −0.35
−0.4
0.0
0.4
0.8
−0.2
0.2
0.6
1.0
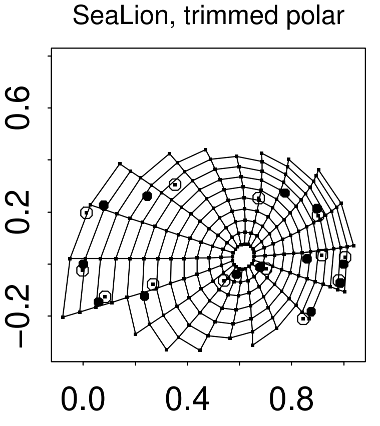
SeaLion, trimmed polar
−0.2
0.2
0.6
0.0
0.4
0.8
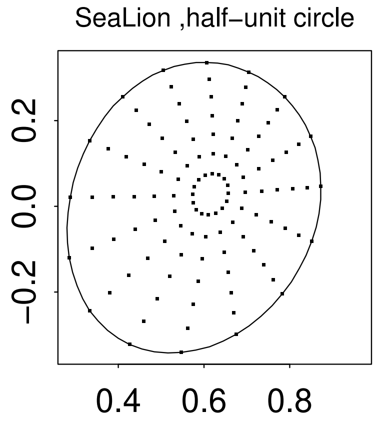
SeaLion ,half−unit circle
−0.2
0.0
0.2
0.4
0.6
0.8
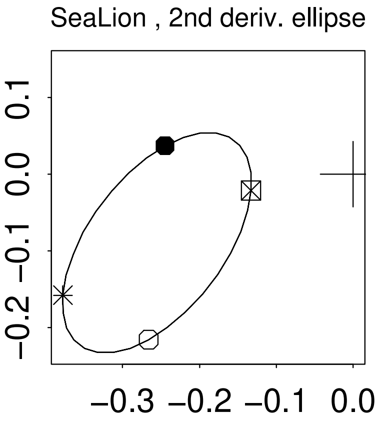
SeaLion , 2nd deriv. ellipse
−0.2
−0.1
0.0
0.1
−0.3
−0.2
−0.1
0.0

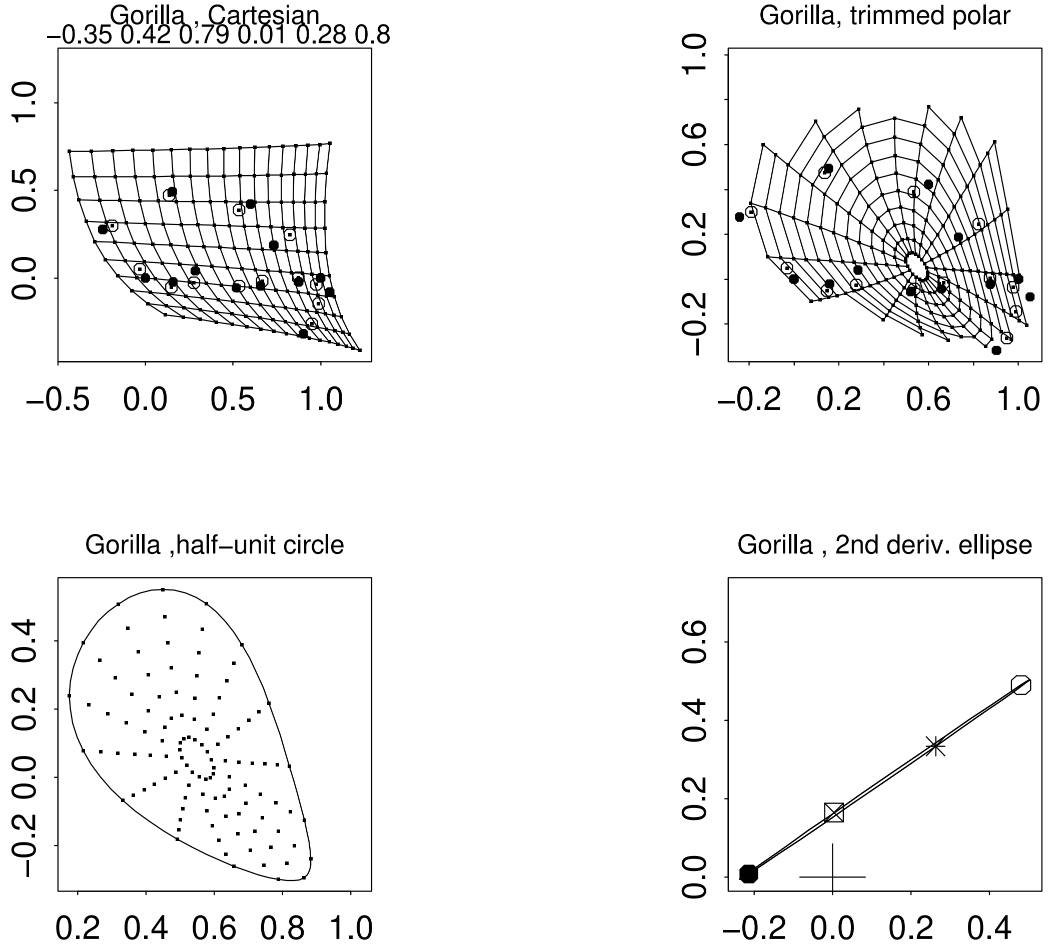
Gorilla , Cartesian
−0.35 0.42 0.79 0.01 0.28 0.8
−0.5 0.0 0.5 1.0
0.0 0.5 1.0
Gorilla, trimmed polar
−0.2 0.2 0.6 1.0
−0.2 0.2 0.6 1.0
Gorilla ,half−unit circle
0.2 0.4 0.6 0.8 1.0
−0.2 0.0 0.2 0.4
Gorilla , 2nd deriv. ellipse
−0.2 0.0 0.2 0.4
0.0 0.2 0.4 0.6

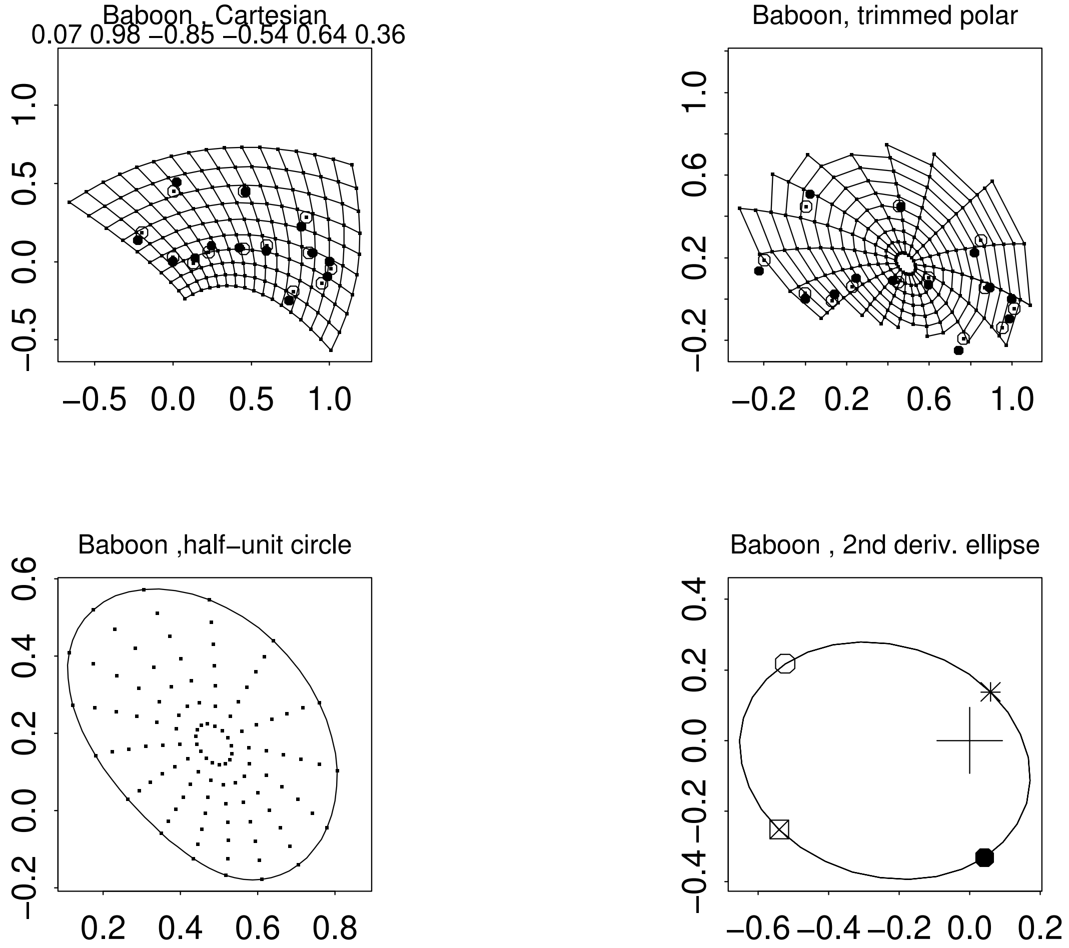
Baboon , Cartesian
0.07 0.98 −0.85 −0.54 0.64 0.36
Baboon, trimmed polar
Baboon ,half−unit circle
Baboon , 2nd deriv. ellipse

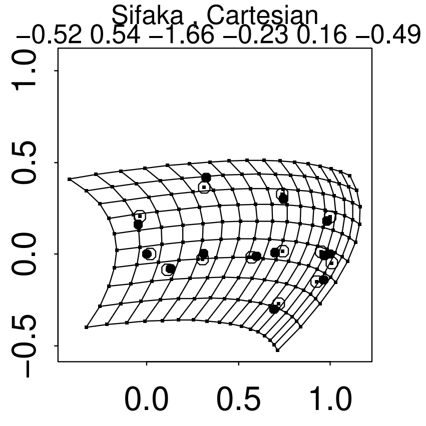
Sifaka , Cartesian
−0.52 0.54 −1.66 −0.23 0.16 −0.49
−0.5 0.0 0.5 1.0
0.0 0.5 1.0

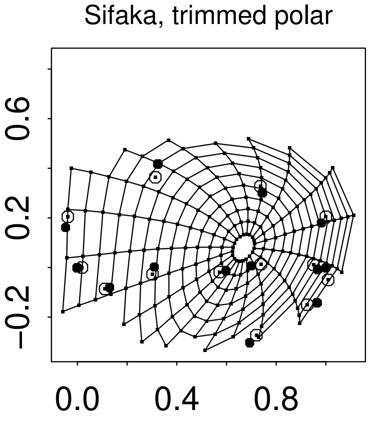
Sifaka, trimmed polar
−0.2 0.2 0.6
0.0 0.4 0.8

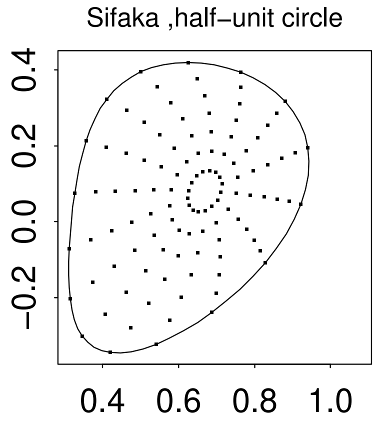
Sifaka ,half−unit circle
−0.2 0.0 0.2 0.4
0.4 0.6 0.8 1.0

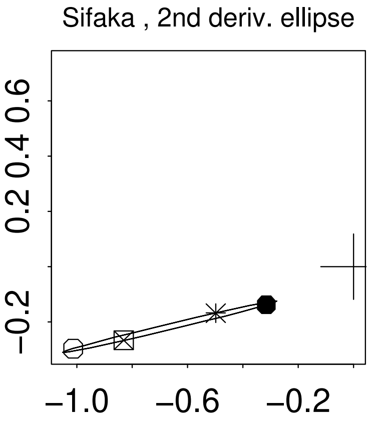
Sifaka , 2nd deriv. ellipse
−0.2 0.2 0.4 0.6
−1.0 −0.6 −0.2

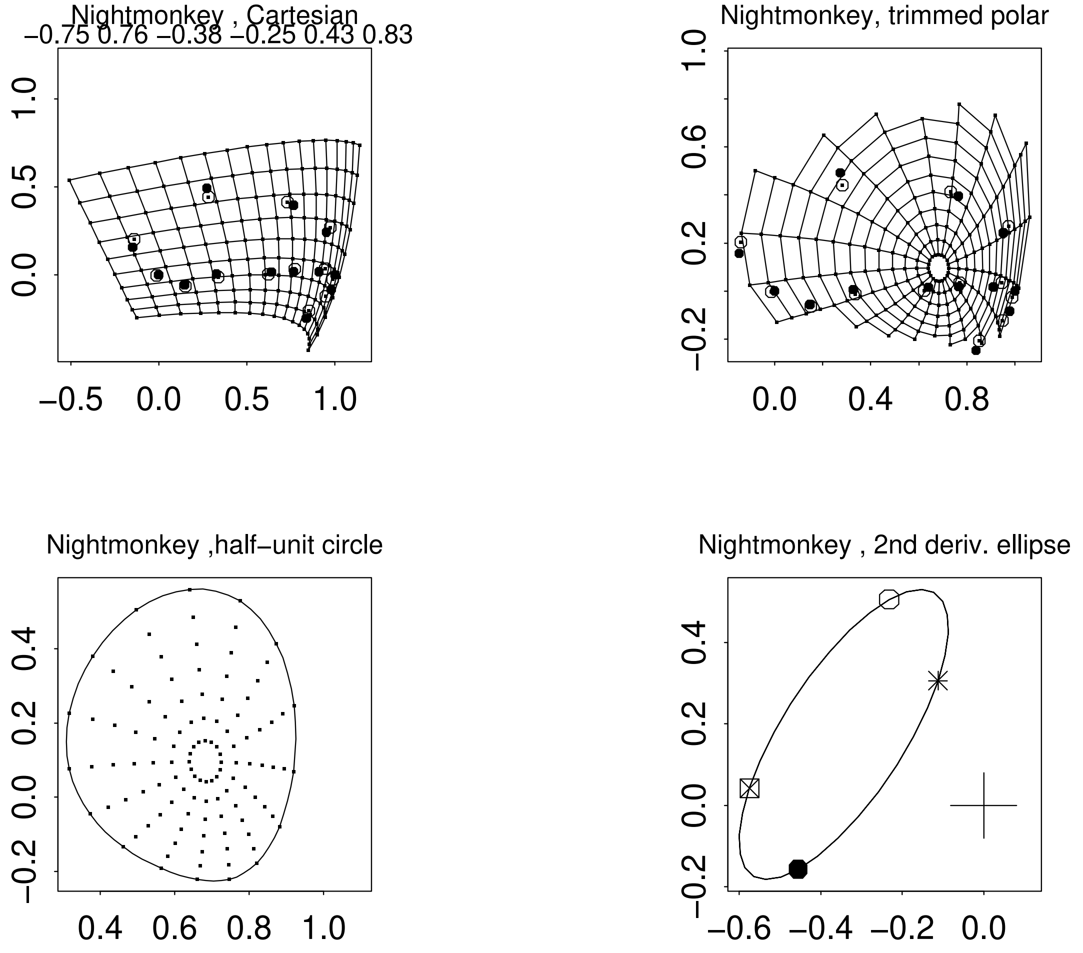

Nightmonkey , Cartesian
−0.75 0.76 −0.38 −0.25 0.43 0.83
Nightmonkey, trimmed polar
Nightmonkey ,half−unit circle
Nightmonkey , 2nd deriv. ellipse

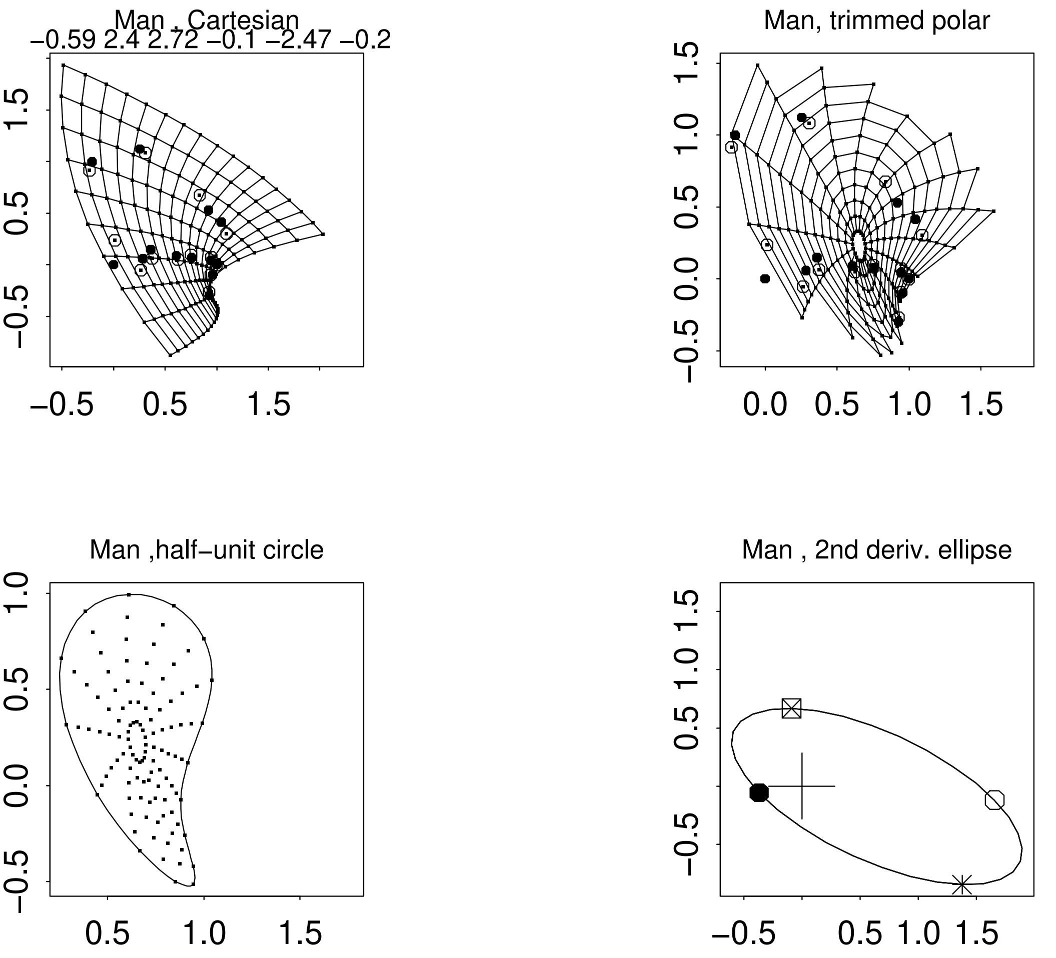
Man , Cartesian
−0.59 2.4 2.72 −0.1 −2.47 −0.2
Man, trimmed polar
Man ,half−unit circle
Man , 2nd deriv. ellipse

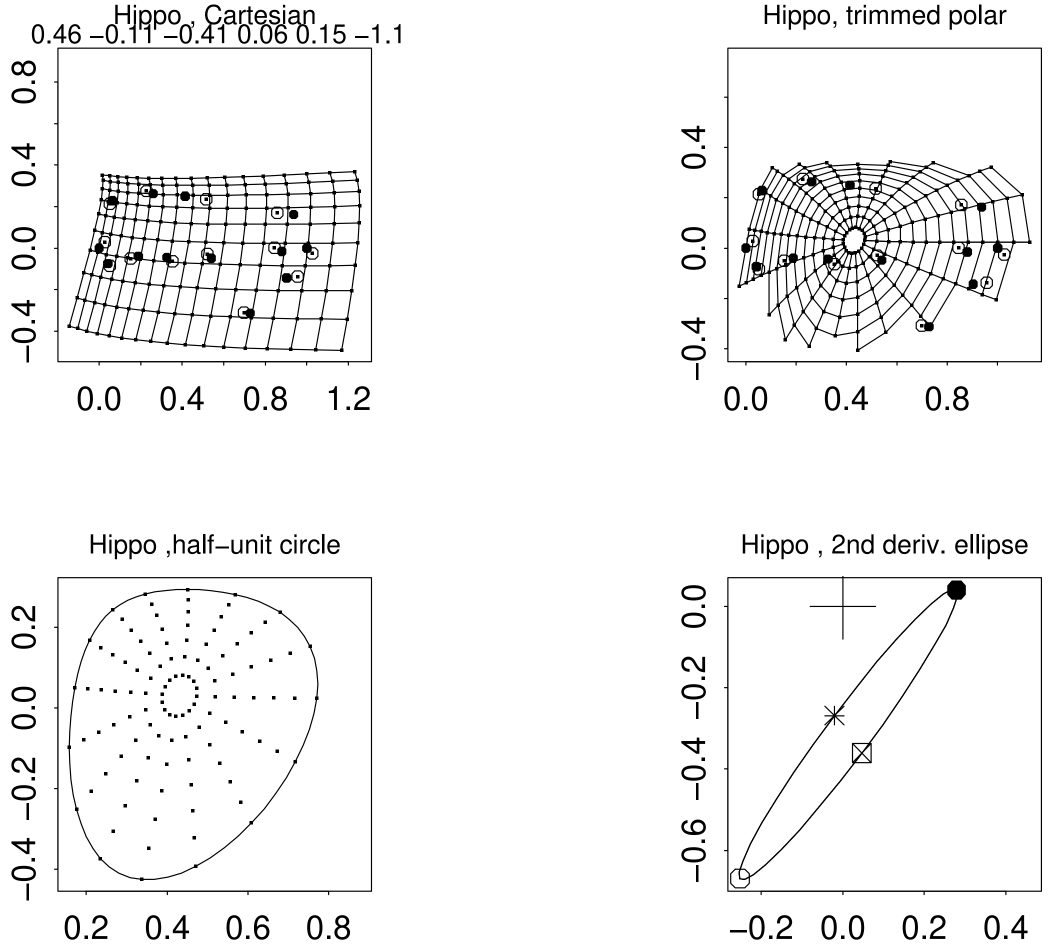

Hippo , Cartesian
0.46 −0.11 −0.41 0.06 0.15 −1.1
Hippo, trimmed polar
Hippo ,half−unit circle
Hippo , 2nd deriv. ellipse

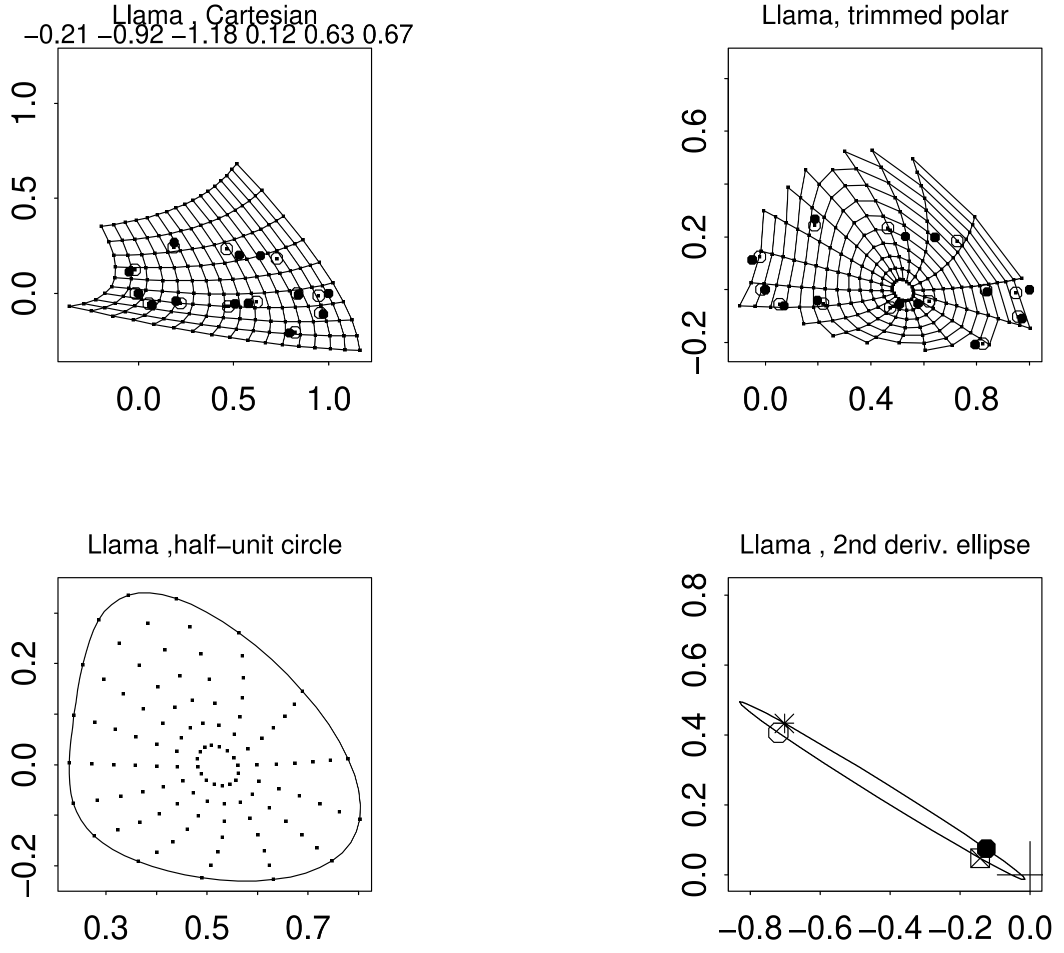
Llama , Cartesian
−0.21 −0.92 −1.18 0.12 0.63 0.67
Llama, trimmed polar
Llama ,half−unit circle
Llama , 2nd deriv. ellipse

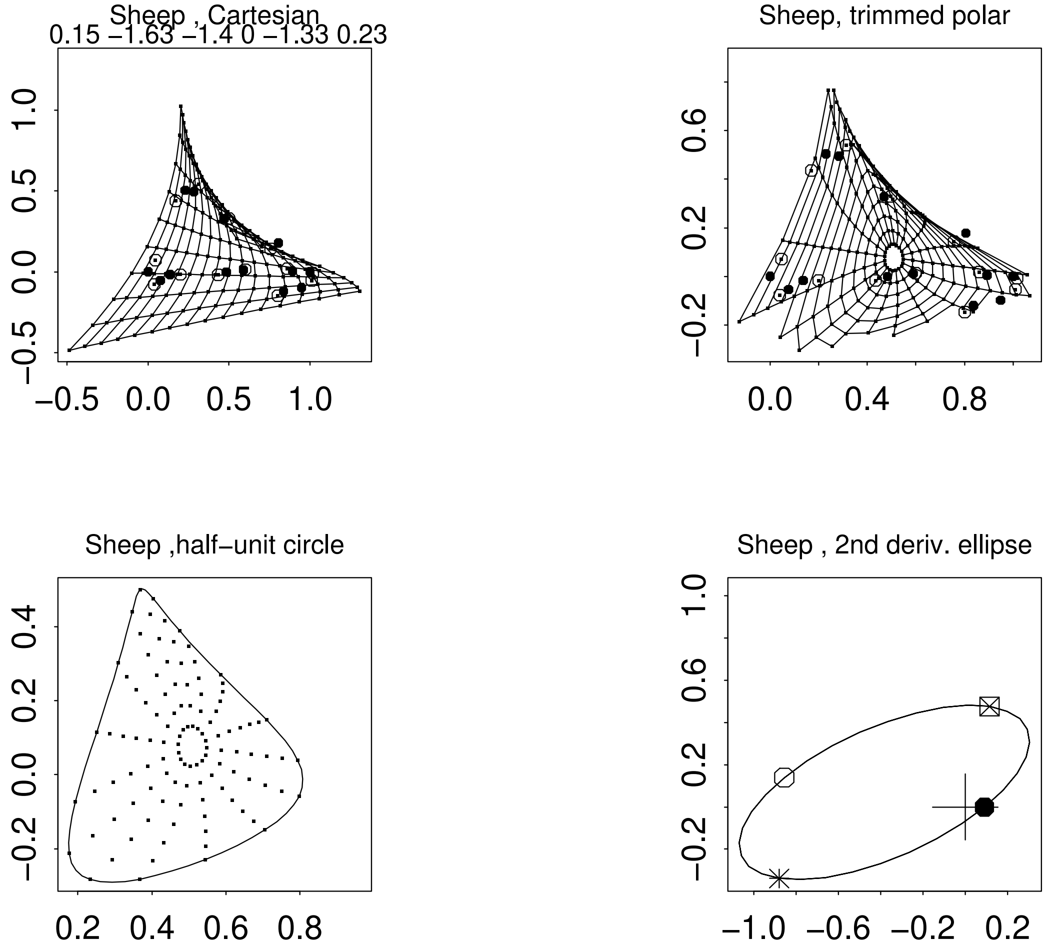
Sheep , Cartesian
0.15 −1.63 −1.4 0 −1.33 0.23
Sheep, trimmed polar
Sheep ,half−unit circle
Sheep , 2nd deriv. ellipse

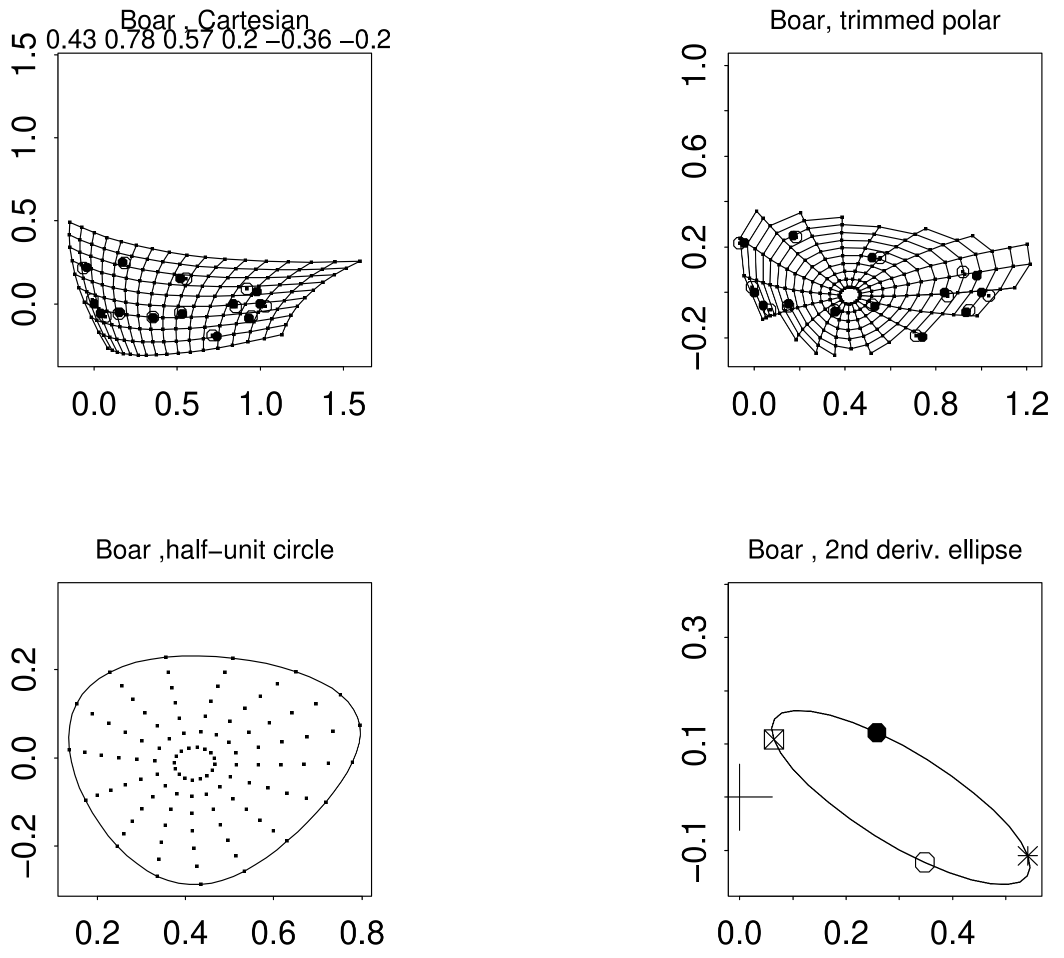
Boar , Cartesian
0.43 0.78 0.57 0.2 −0.36 −0.2
Boar, trimmed polar
Boar ,half−unit circle
Boar , 2nd deriv. ellipse

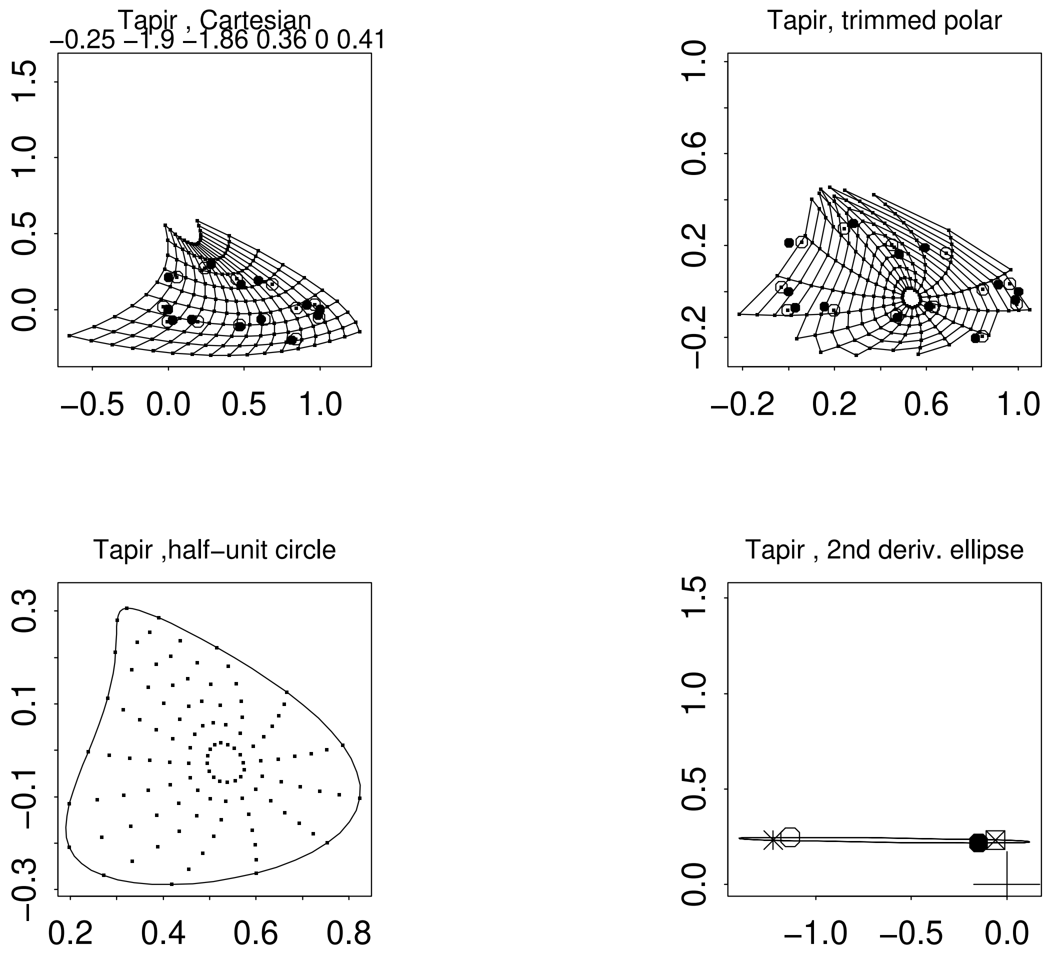
Tapir , Cartesian
−0.25 −1.9 −1.86 0.36 0 0.41
Tapir, trimmed polar
Tapir ,half−unit circle
Tapir , 2nd deriv. ellipse

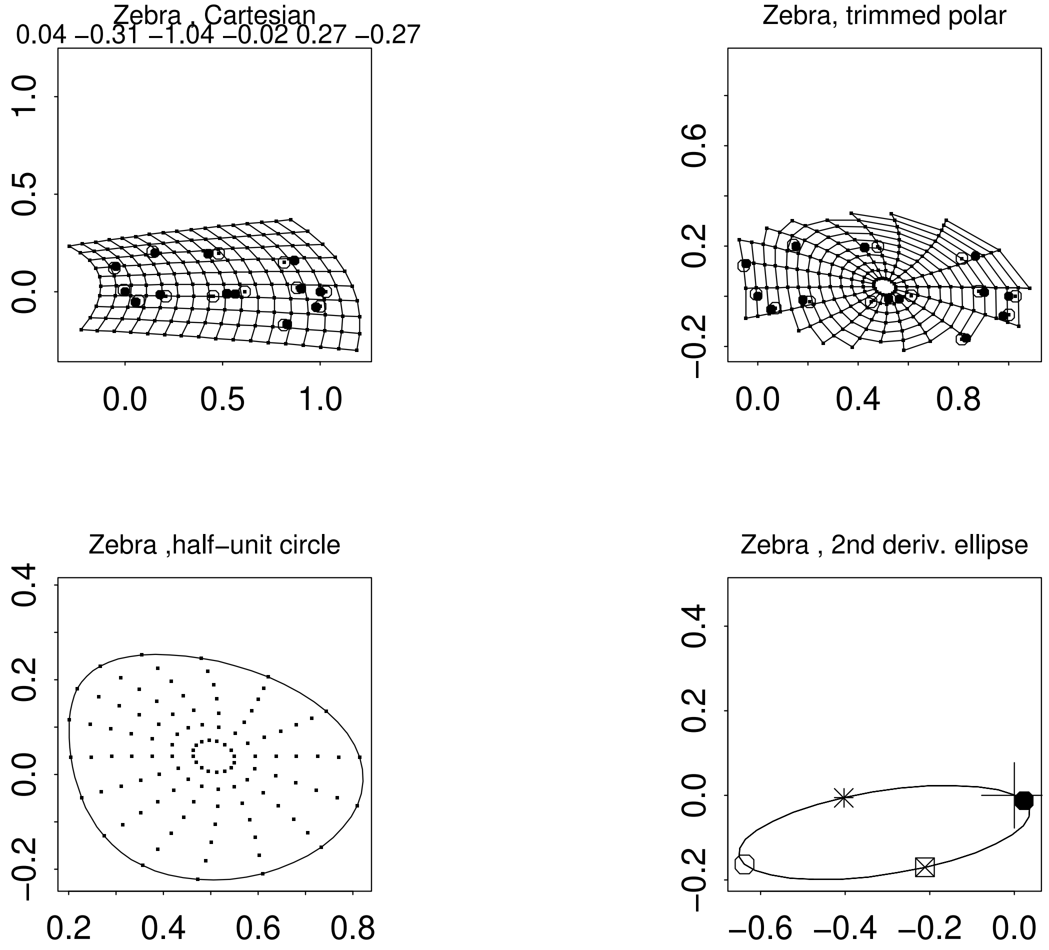
Zebra , Cartesian
0.04 −0.31 −1.04 −0.02 0.27 −0.27
Zebra, trimmed polar
Zebra ,half−unit circle
Zebra , 2nd deriv. ellipse

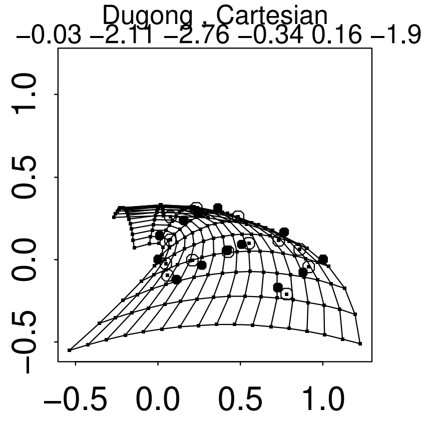
Dugong , Cartesian
−0.03 −2.11 −2.76 −0.34 0.16 −1.9

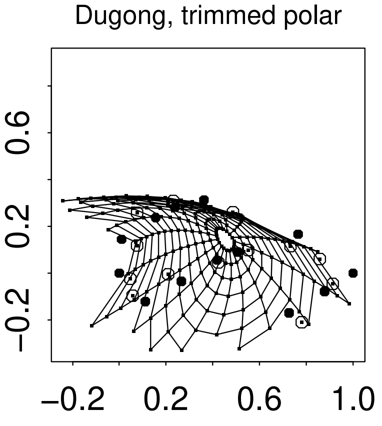
Dugong, trimmed polar

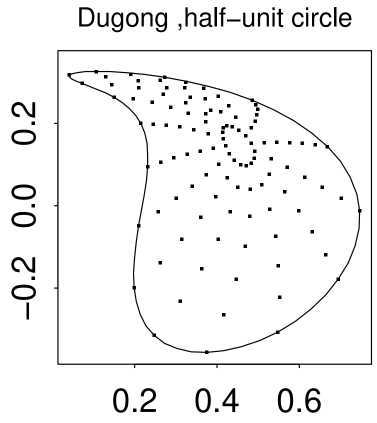
Dugong ,half−unit circle

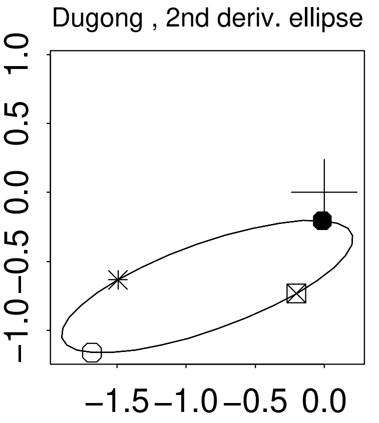
Dugong , 2nd deriv. ellipse

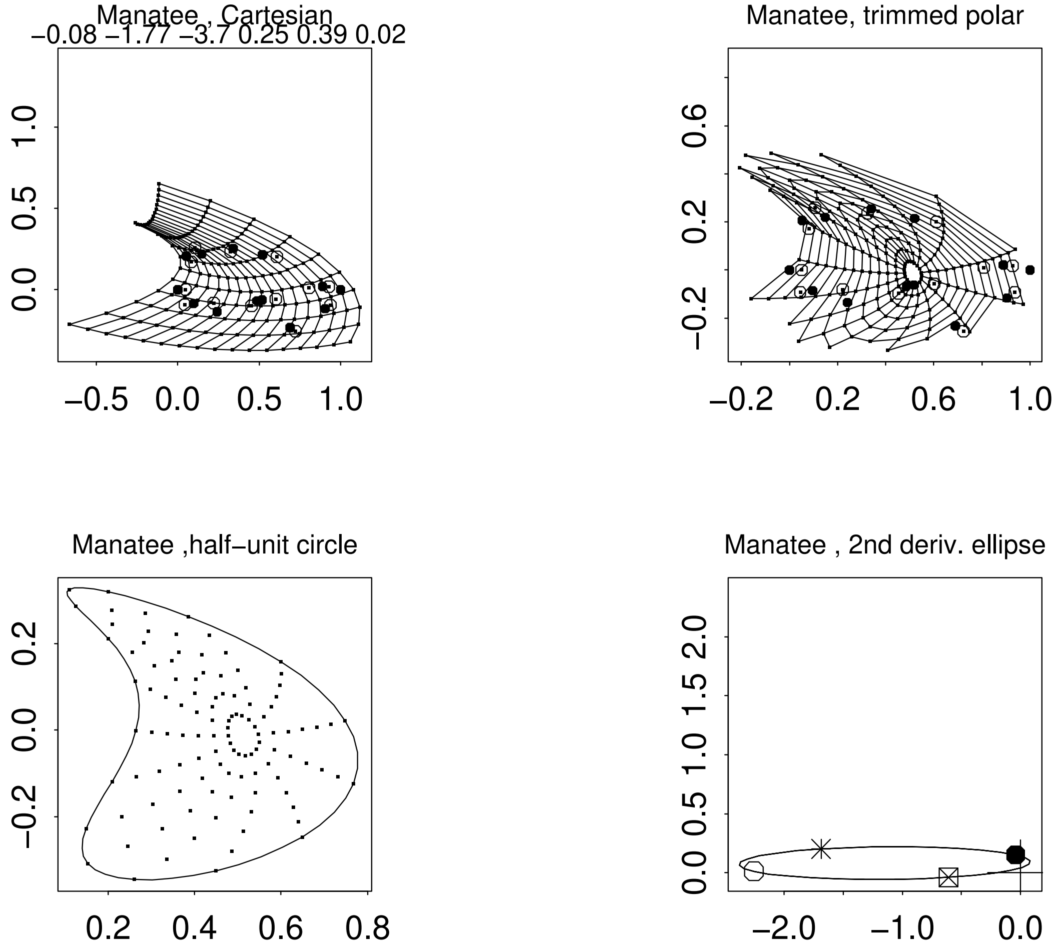
Manatee , Cartesian
−0.08 −1.77 −3.7 0.25 0.39 0.02
1.0
0.5
0.0
−0.5 0.0 0.5 1.0
Manatee, trimmed polar
0.6
0.2
−0.2
−0.2 0.2 0.6 1.0
Manatee ,half−unit circle
0.2
0.0
−0.2
0.2 0.4 0.6 0.8
Manatee , 2nd deriv. ellipse
0.0 0.5 1.0 1.5 2.0
−2.0 −1.0 0.0

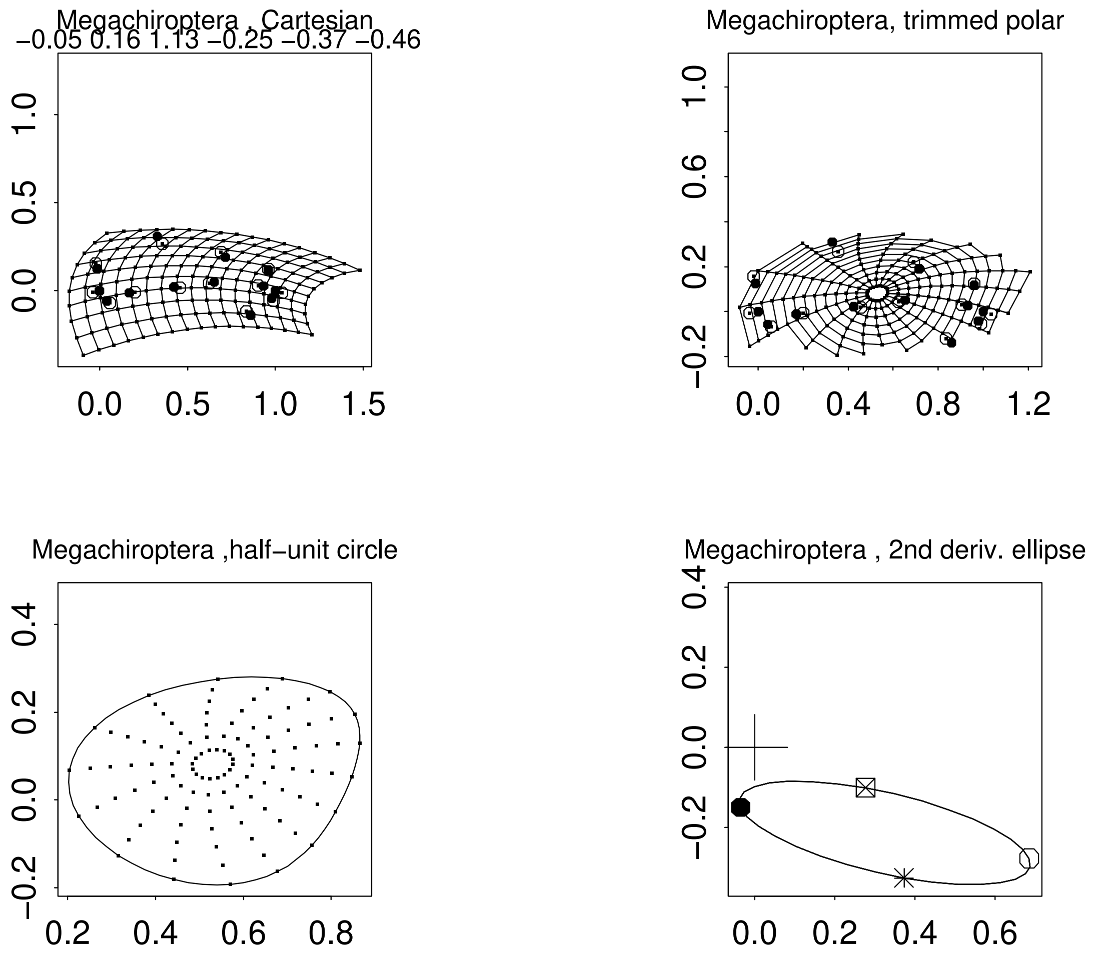
Megachiroptera , Cartesian
−0.05 0.16 1.13 −0.25 −0.37 −0.46
Megachiroptera, trimmed polar
Megachiroptera ,half−unit circle
Megachiroptera , 2nd deriv. ellipse

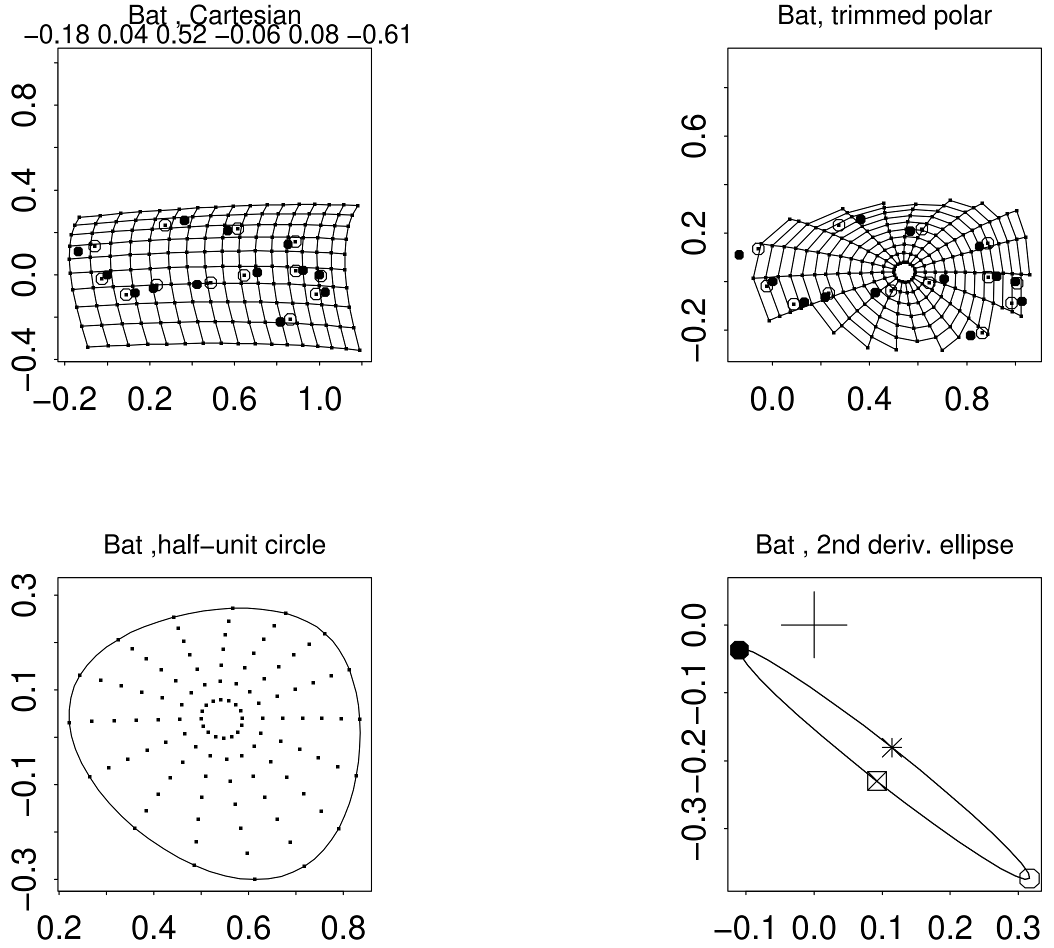
Bat , Cartesian
−0.18 0.04 0.52 −0.06 0.08 −0.61
Bat, trimmed polar
Bat ,half−unit circle
Bat , 2nd deriv. ellipse

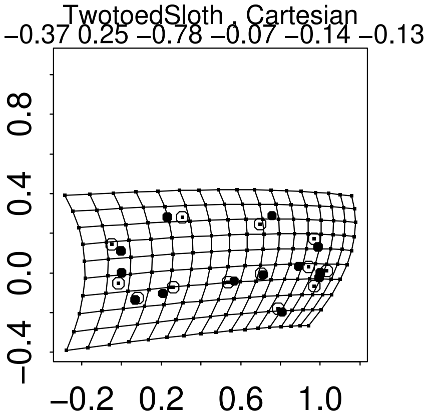
TwotoedSloth , Cartesian
−0.37 0.25 −0.78 −0.07 −0.14 −0.13
−0.4 0.0 0.4 0.8
−0.2 0.2 0.6 1.0

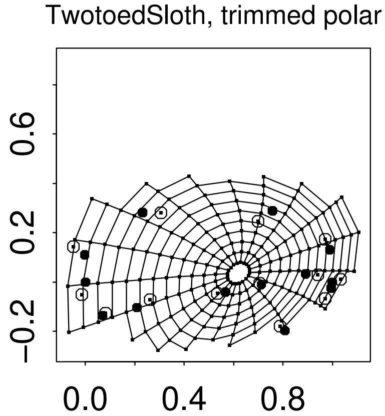
TwotoedSloth, trimmed polar
−0.2 0.2 0.6
0.0 0.4 0.8

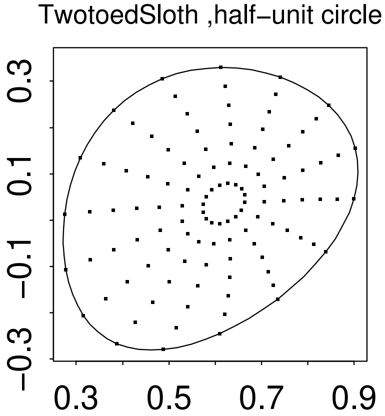
TwotoedSloth ,half−unit circle
−0.3 −0.1 0.1 0.3
0.3 0.5 0.7 0.9

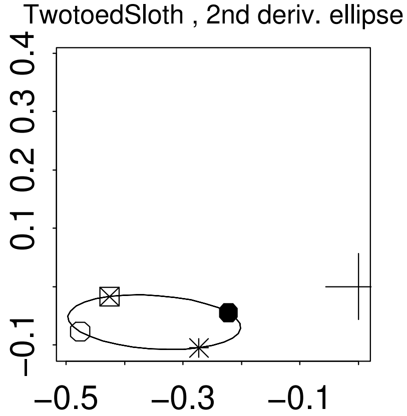
TwotoedSloth , 2nd deriv. ellipse
−0.1 0.1 0.2 0.3 0.4
−0.5 −0.3 −0.1

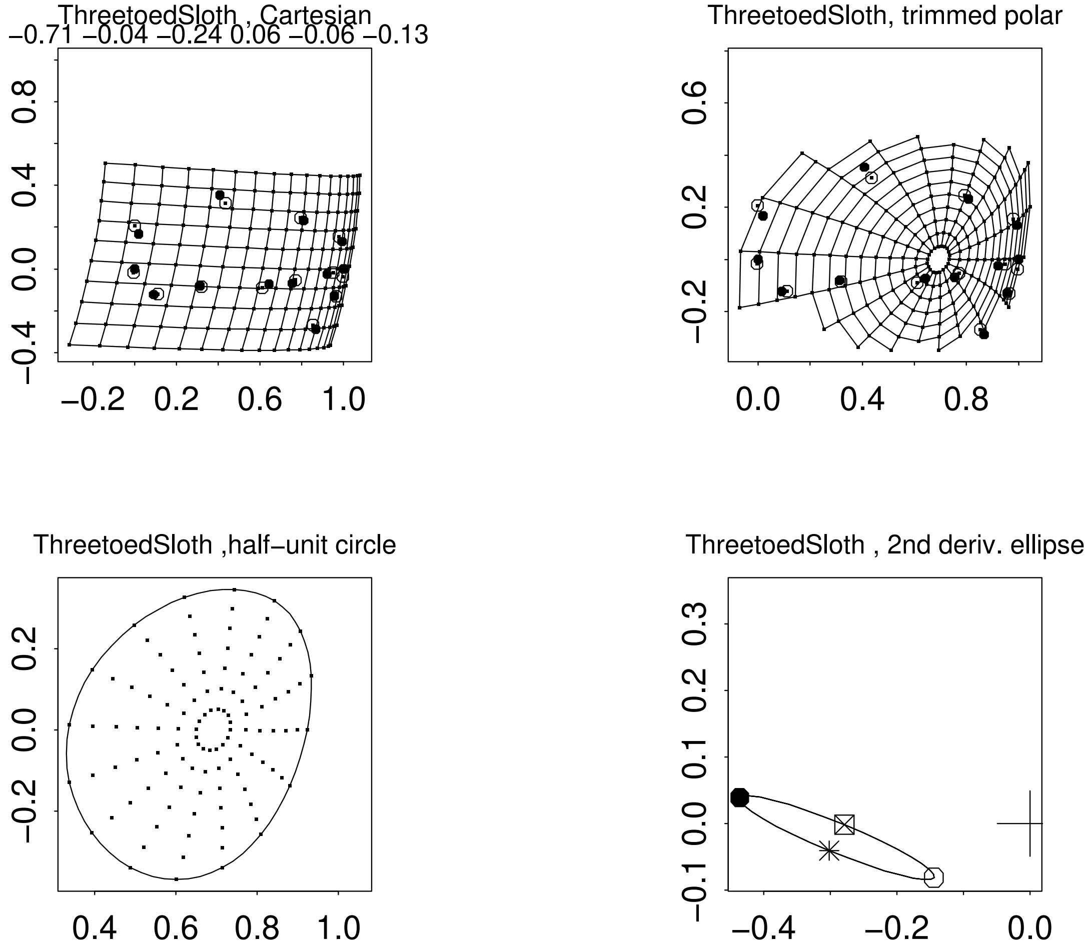
ThreetoedSloth , Cartesian
−0.71 −0.04 −0.24 0.06 −0.06 −0.13
ThreetoedSloth, trimmed polar
ThreetoedSloth ,half−unit circle
ThreetoedSloth , 2nd deriv. ellipse

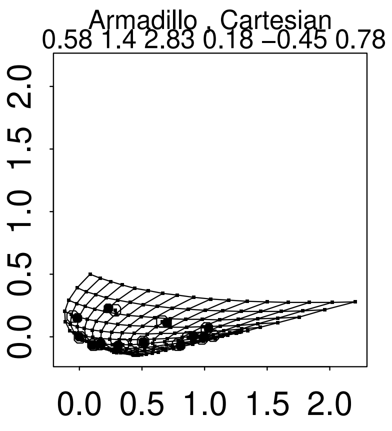
Armadillo , Cartesian
0.58 1.4 2.83 0.18 −0.45 0.78
0.0 0.5 1.0 1.5 2.0

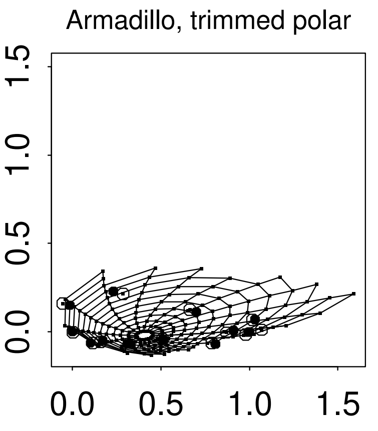
Armadillo, trimmed polar
0.0 0.5 1.0 1.5

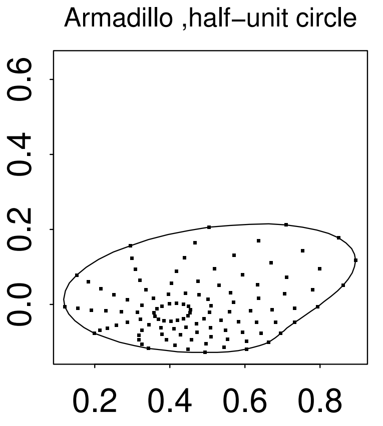
Armadillo ,half−unit circle
0.2 0.4 0.6 0.8
0.0 0.2 0.4 0.6

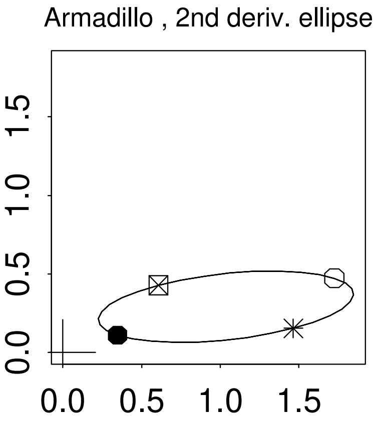
Armadillo , 2nd deriv. ellipse
0.0 0.5 1.0 1.5

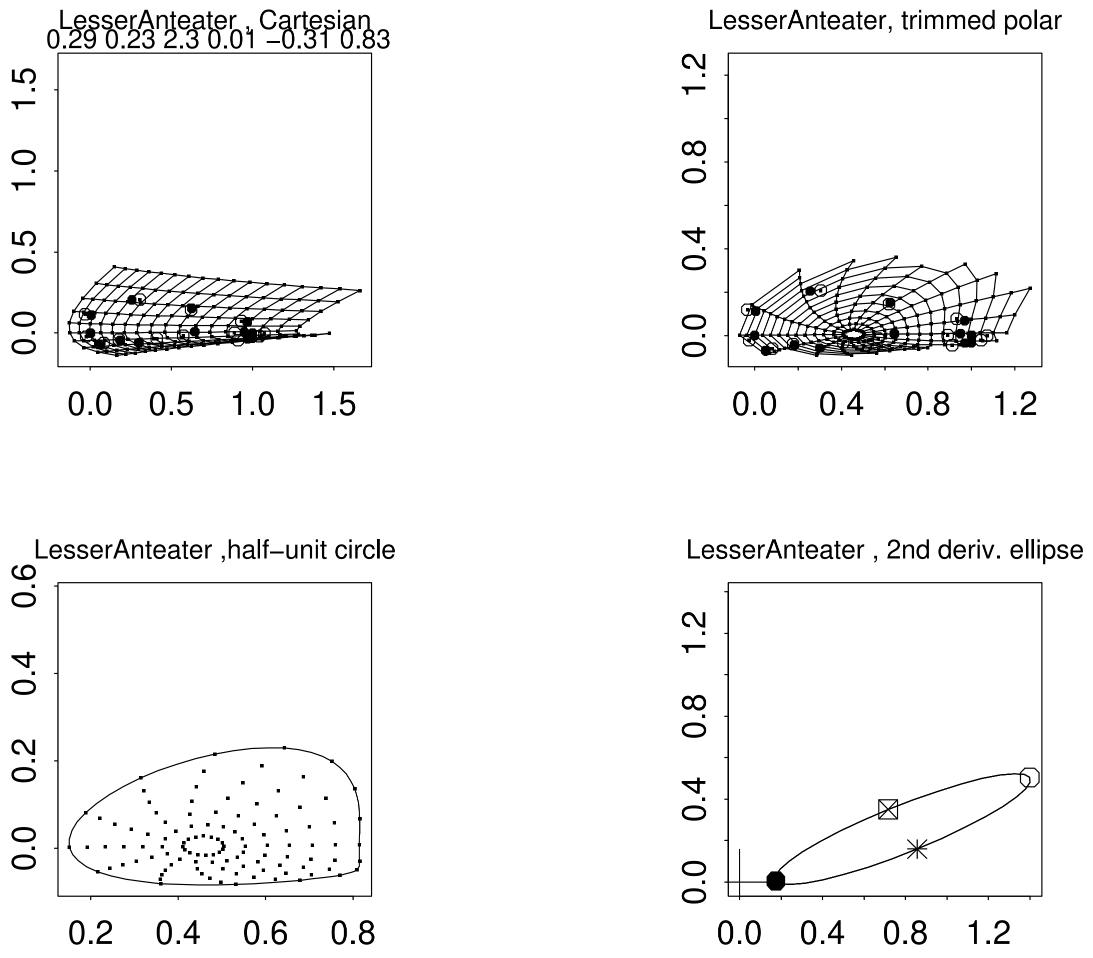
LesserAnteater , Cartesian
0.29 0.23 2.3 0.01 −0.31 0.83
0.0 0.5 1.0 1.5
LesserAnteater, trimmed polar
0.0 0.4 0.8 1.2
LesserAnteater ,half−unit circle
0.2 0.4 0.6 0.8
0.0 0.2 0.4 0.6
LesserAnteater , 2nd deriv. ellipse
0.0 0.4 0.8 1.2

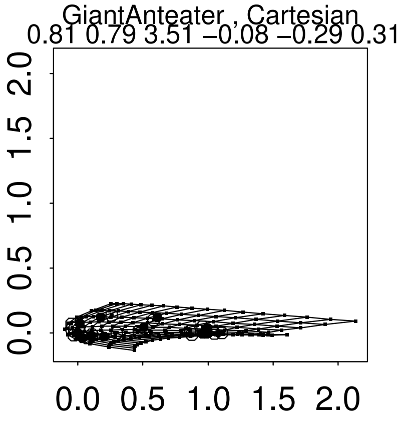
GiantAnteater , Cartesian
0.81 0.79 3.51 −0.08 −0.29 0.31
0.0 0.5 1.0 1.5 2.0
0.0 0.5 1.0 1.5 2.0

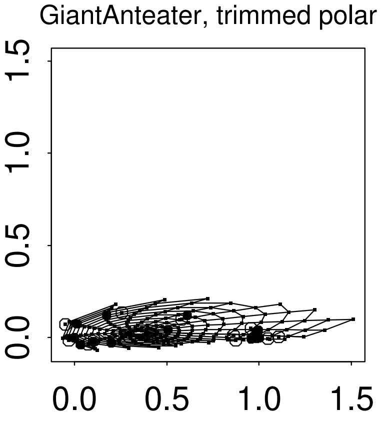
GiantAnteater, trimmed polar
0.0 0.5 1.0 1.5
0.0 0.5 1.0 1.5

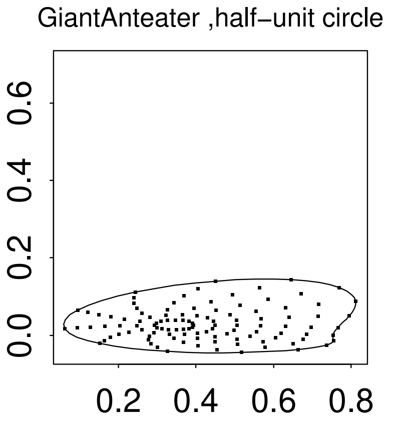
GiantAnteater ,half−unit circle
0.0 0.2 0.4 0.6
0.2 0.4 0.6 0.8

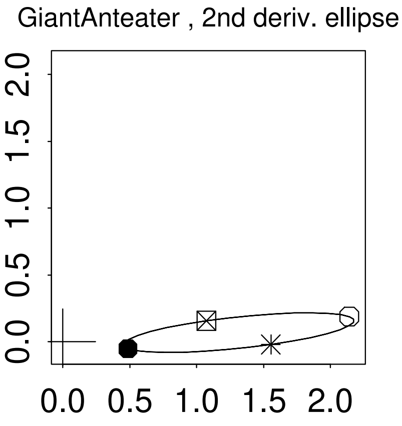
GiantAnteater , 2nd deriv. ellipse
0.0 0.5 1.0 1.5 2.0
0.0 0.5 1.0 1.5 2.0

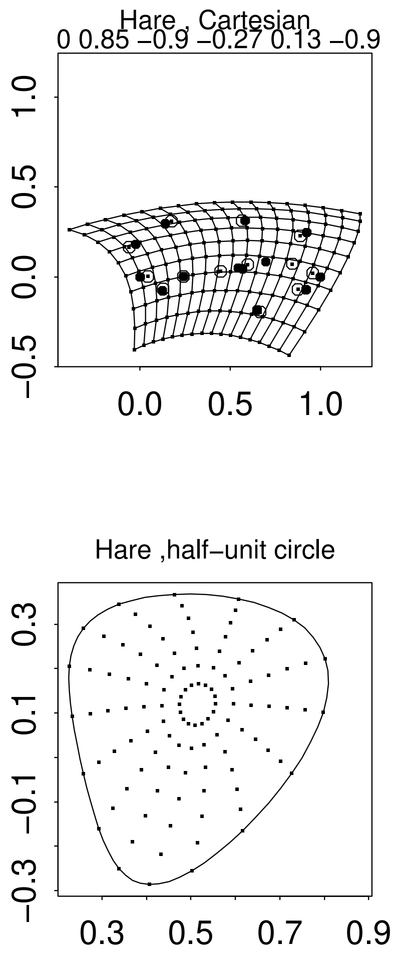

Hare , Cartesian
0 0.85 −0.9 −0.27 0.13 −0.9
Hare, trimmed polar
Hare ,half−unit circle
Hare , 2nd deriv. ellipse

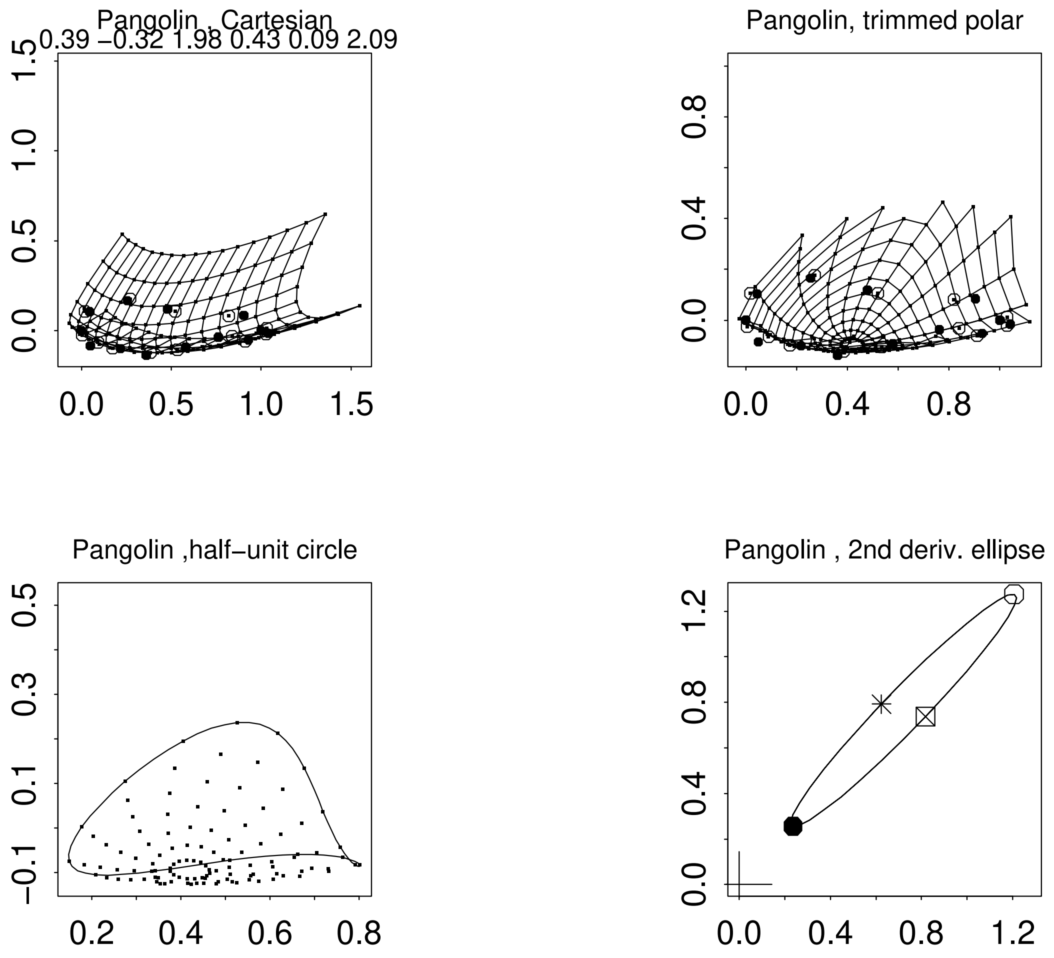
Pangolin , Cartesian
0.39 −0.32 1.98 0.43 0.09 2.09
Pangolin, trimmed polar
Pangolin ,half−unit circle
Pangolin , 2nd deriv. ellipse

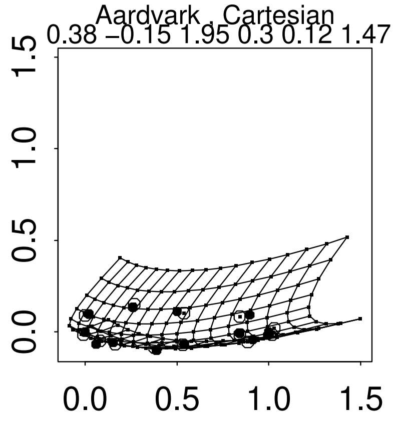
Aardvark , Cartesian
0.38 −0.15 1.95 0.3 0.12 1.47
0.0 0.5 1.0 1.5
0.0 0.5 1.0 1.5

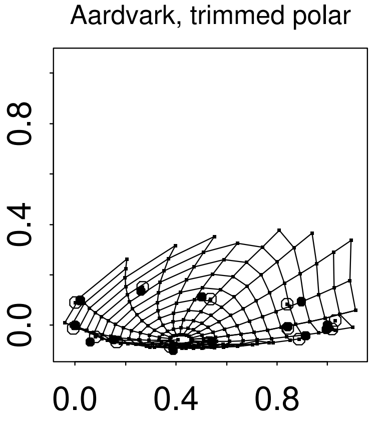
Aardvark, trimmed polar
0.0 0.4 0.8
0.0 0.4 0.8

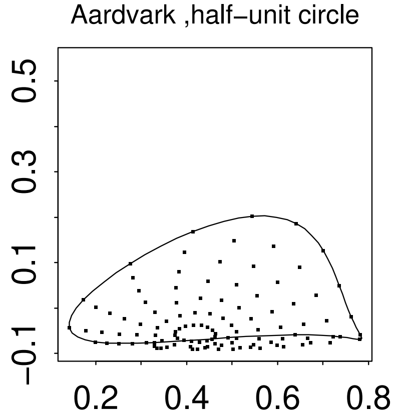
Aardvark ,half−unit circle
−0.1 0.1 0.3 0.5
0.2 0.4 0.6 0.8

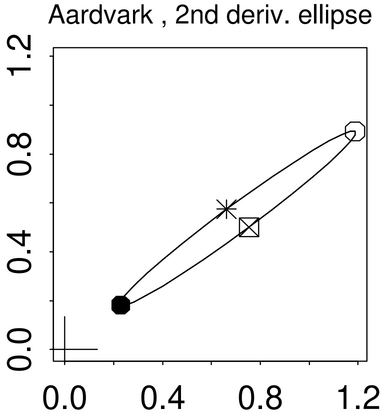
Aardvark , 2nd deriv. ellipse
0.0 0.4 0.8 1.2
0.0 0.4 0.8 1.2

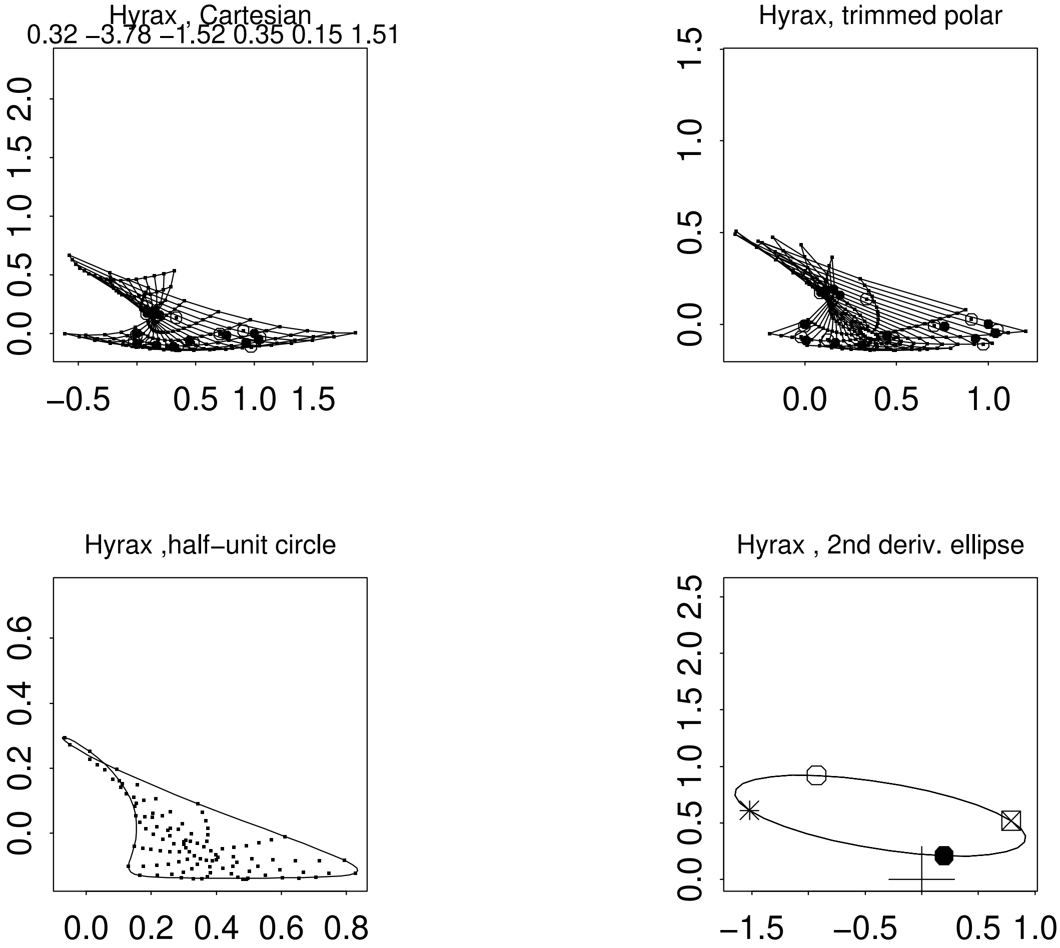

Hyrax , Cartesian
0.32 −3.78 −1.52 0.35 0.15 1.51
Hyrax, trimmed polar
Hyrax ,half−unit circle
Hyrax , 2nd deriv. ellipse

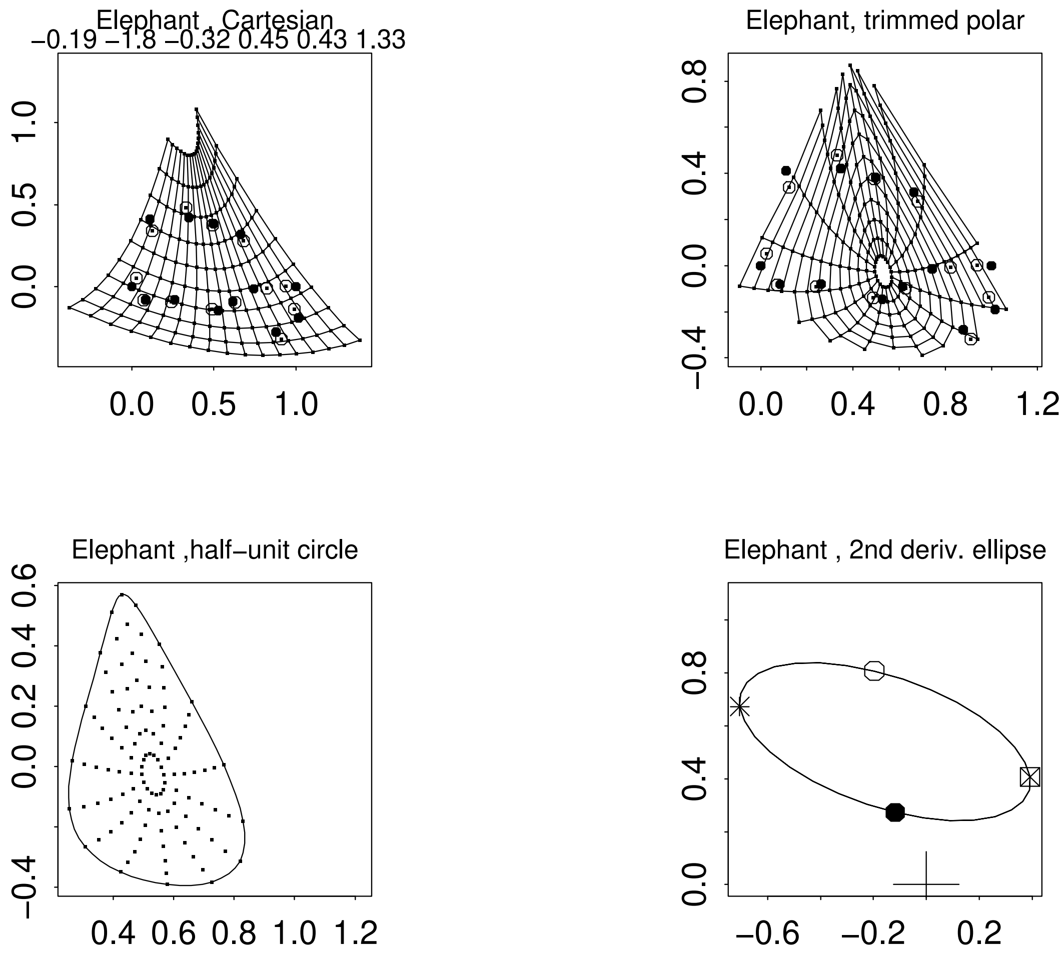
Elephant , Cartesian
−0.19 −1.8 −0.32 0.45 0.43 1.33
0.0 0.5 1.0
0.0 0.5 1.0
Elephant, trimmed polar
0.0 0.4 0.8 1.2
−0.4 0.0 0.4 0.8
Elephant ,half−unit circle
0.4 0.6 0.8 1.0 1.2
−0.4 0.0 0.2 0.4 0.6
Elephant , 2nd deriv. ellipse
−0.6 −0.2 0.2
0.0 0.4 0.8

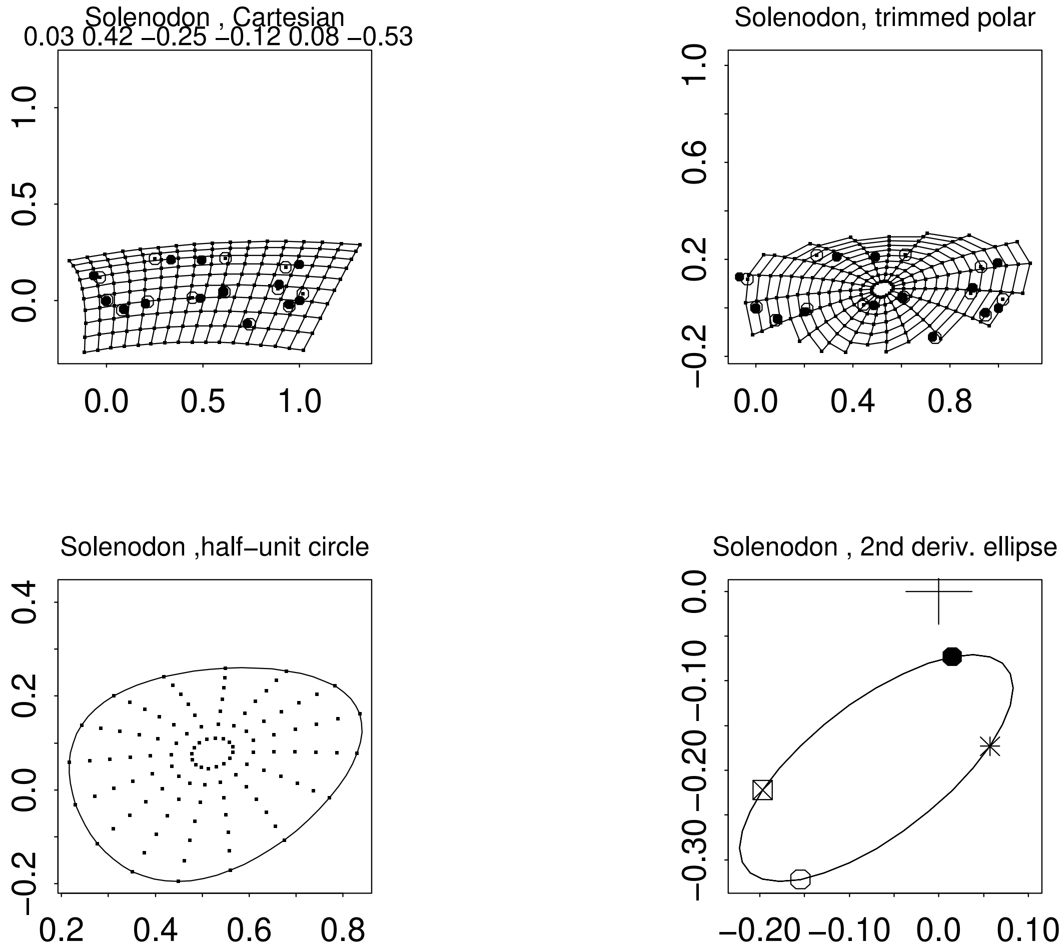
Solenodon , Cartesian
0.03 0.42 −0.25 −0.12 0.08 −0.53
Solenodon, trimmed polar
Solenodon ,half−unit circle
Solenodon , 2nd deriv. ellipse

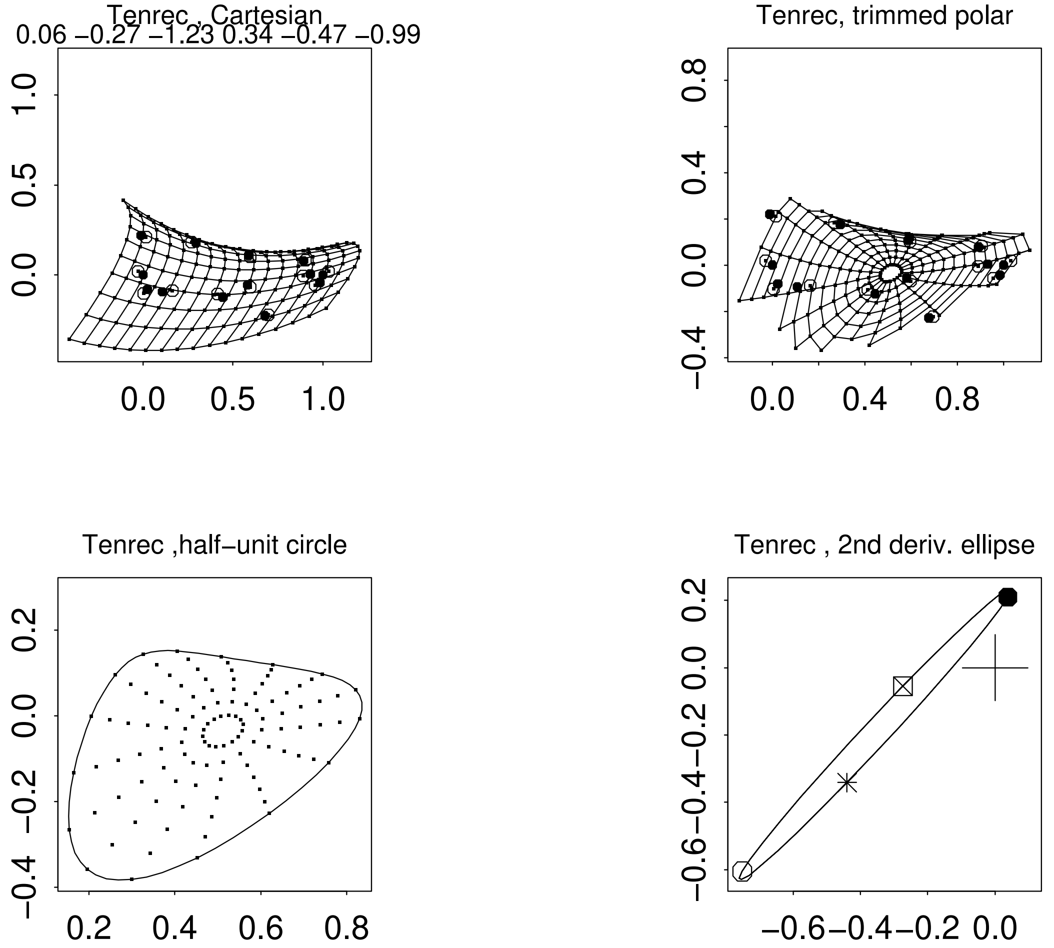
Tenrec , Cartesian
0.06 −0.27 −1.23 0.34 −0.47 −0.99
1.0
0.5
0.0
0.0
0.5
1.0
Tenrec, trimmed polar
0.8
0.4
0.0
−0.4
0.0
0.4
0.8
Tenrec ,half−unit circle
0.2
0.0
−0.2
−0.4
0.2
0.4
0.6
0.8
Tenrec , 2nd deriv. ellipse
0.2
0.0
−0.2
−0.4
−0.6
−0.6
−0.4
−0.2
0.0

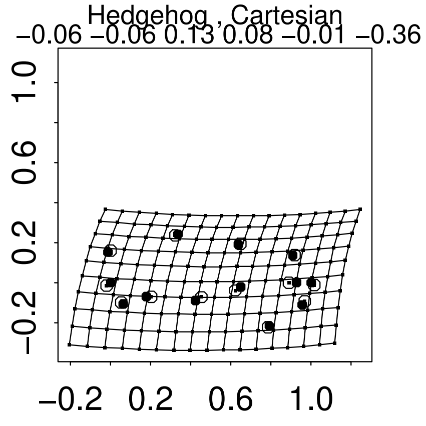
Hedgehog , Cartesian
−0.06 −0.06 0.13 0.08 −0.01 −0.36
−0.2 0.2 0.6 1.0
−0.2 0.2 0.6 1.0

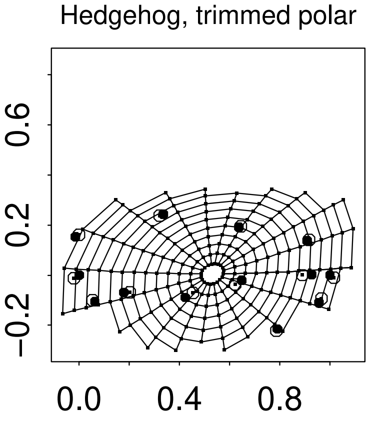
Hedgehog, trimmed polar
−0.2 0.2 0.6
0.0 0.4 0.8

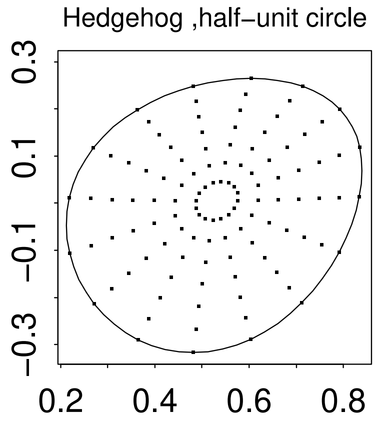
Hedgehog ,half−unit circle
−0.3 −0.1 0.1 0.3
0.2 0.4 0.6 0.8

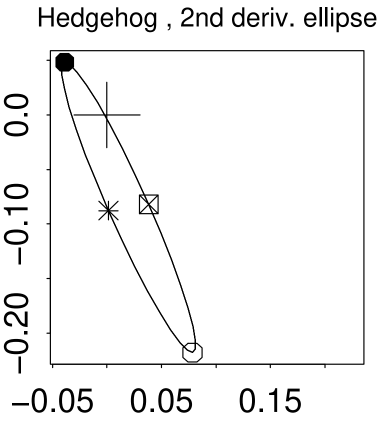
Hedgehog , 2nd deriv. ellipse
−0.20 −0.10 0.0
−0.05 0.05 0.15

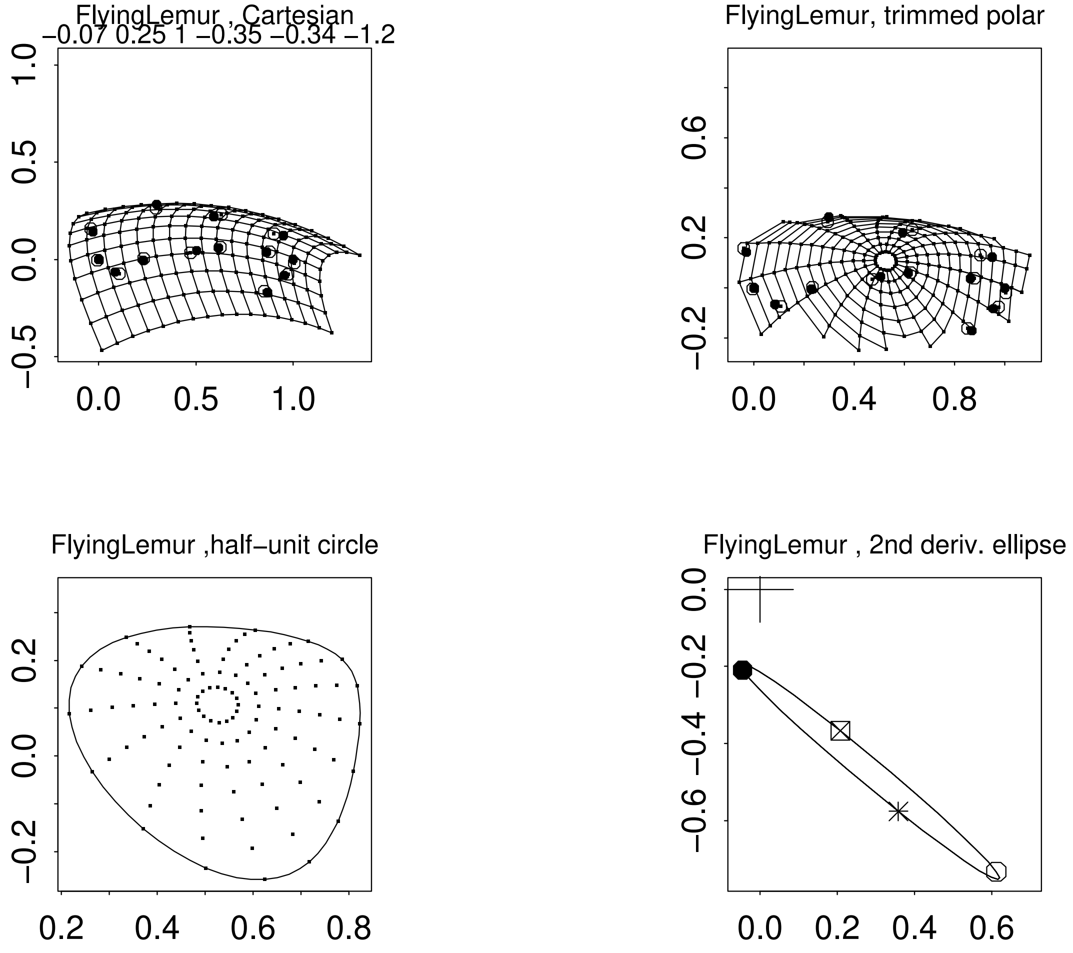
FlyingLemur , Cartesian
−0.07 0.25 1 −0.35 −0.34 −1.2
1.0
0.5
0.0
−0.5
0.0
0.5
1.0
FlyingLemur, trimmed polar
0.6
0.2
−0.2
0.0
0.4
0.8
FlyingLemur ,half−unit circle
0.2
0.0
−0.2
0.2
0.4
0.6
0.8
FlyingLemur , 2nd deriv. ellipse
0.0
−0.2
−0.4
−0.6
0.0
0.2
0.4
0.6

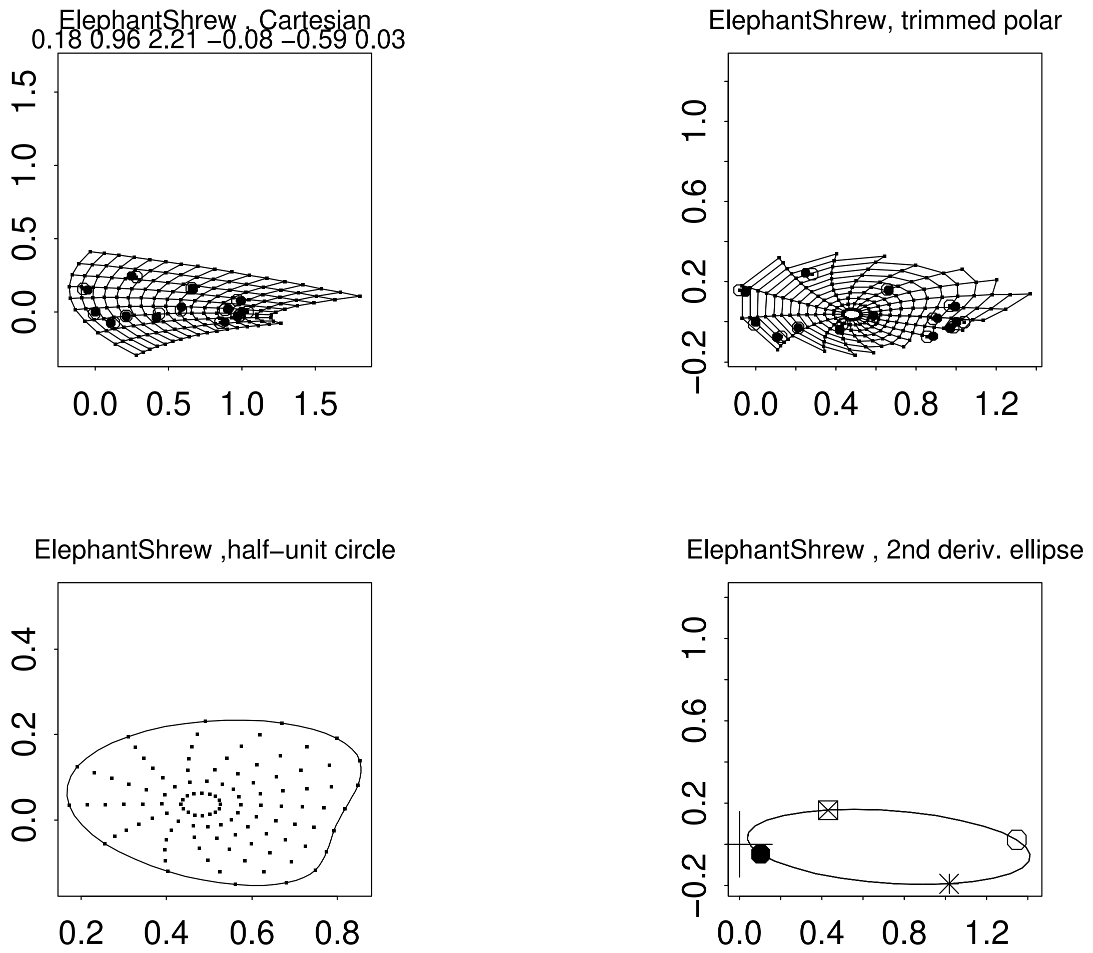
ElephantShrew , Cartesian
0.18 0.96 2.21 −0.08 −0.59 0.03
ElephantShrew, trimmed polar
ElephantShrew ,half−unit circle
ElephantShrew , 2nd deriv. ellipse
